# Detection of Antibodies Against the African Parasite *Trypanosoma brucei* Using Synthetic glycosylphosphatidylinositol oligosaccharide fragments

**DOI:** 10.1101/2023.12.31.573764

**Authors:** Maurice Michel, Benoît Stijlemans, Dana Michel, Monika Garg, Andreas Geissner, Peter H. Seeberger, Daniel Varón Silva

## Abstract

*Trypanosoma brucei* (*T. brucei*) parasites cause two major infectious diseases in Africa: African trypanosomiasis in humans (HAT) and Nagana in animals. Despite the enormous economic and social impact, vaccines and reliable diagnostic measures are still lacking for these diseases. The main obstacle to developing accurate diagnostic methods and an active vaccine is the parasite’s ability for antigenic variation and impairment of B cell maturation, which prevents the development of a long-lasting, effective immune response. The antigenic variation is sustained by random gene switching, segmental gene conversion, and altered glycosylation states of solvent-exposed regions of the corresponding variant surface glycoproteins (VSG). These glycoproteins use a glycosylphosphatidylinositol (GPI) anchor for attachment to the membrane. GPIs of *T. brucei* have specific branched structures that are further heterogeneously galactosylated. We synthesized a glycan fragment library containing *T. brucei* GPIs most prominent structural features and performed an epitope mapping using mice and human sera of infected specimens using glycan microarrays. The studies indicate that in contrast to VSG, *T. brucei* GPIs are recognized by both short-lived IgM and long-lasting IgG, indicating a specific immune response against GPI structures. These findings enable the development of diagnostic tests based on synthetic antigens for reliable diagnosis of human African trypanosomiasis and Nagana.

## Introduction

Subspecies of the African extracellular parasite *T. brucei* cause two infectious diseases in rural areas of Africa, human African Trypanosomiasis (HAT) and Nagana in animals. Limited diagnosis and treatment, a lack of trained point-of-care personnel, and restricted access to medical facilities have created a beneficial environment for the parasite and its vector, the tsetse fly.[1,2] A *T. brucei* infection is characterized by a haemolymphatic phase that displays symptoms such as weakness and fever and a neurologic phase associated with severe anemia, sleep cycle disruption and progressive mental deterioration.[3] The symptoms of the first phase are not uncommon in sub-Saharan Africa and often leave the infection undiagnosed in animals and humans. Thereby, HAT can progress to the second stage and becomes lethal if not treated by chemotherapy.[4,5]

*T. brucei* is one of the most persistent parasites infecting humans. The parasite’s long-term survival of the hostile innate and adaptive human immune system is achieved by several mechanisms involving heterogeneity and structural organization of cell surface antigens.[6,7] The outer surface of *T. brucei* is covered by a dense coat of a single phenotype of a variant surface glycoprotein (VSG) that is attached via a glycosylphosphatidylinositol (GPI) anchor.[8,9] VSGs participate in the complement system inhibition, installation of a diffusion barrier, antibody scavenging, masking of other surface proteins and induce the production of autoantibodies.[10–14] Furthermore, *T. brucei* uses VSGs for antigen variation by randomly expressing one VSG construct out of several hundred genes in the genome. The possibility of switching the responsible gene between generations increases the probability of immune system evasion over several parasite generations.[15–17] Segmental gene conversion and mosaicism translate to a new unique phenotype in the solvent-exposed N-terminal and C-terminal domains, respectively.[18–23] Recent reports also showed VSG sequences displaying a third glycosylation site at the top of the solvent-exposed N-terminus covering the amino acid sequence.[21]

The heterogeneity of surface antigens hinders their application for developing consistent diagnostic methods and vaccines against *T. brucei* infections. Currently, trypanosomiasis diagnosis is divided into three stages.[3,22–24] The first stage includes screening for infections by serological tests and analysis of clinical signs, i.e., swollen lymph nodes. The second and third stages involve microscopic confirmation of parasite presence in the blood (infection in phase 1) and the cerebrospinal fluid (infection in phase 2). A commonly used serological test is the card agglutination test for trypanosomiasis (CATT). This test shows improved thermostability, selectivity and specificity but is limited to detecting infections of only a subset of *T. brucei gambiense (Tbg)* strains expressing the VSG LiTaT1.3 variant.[25] Furthermore, the test is not objectively reproducible and is based on fixed *Tbg* parasites, which demand a constant supply of cultivated parasites and trained medical personnel. Determination of the infection phase is essential to select the appropriate therapy and the need to use crossing blood-brain barrier drugs for a phase 2 infection. In recent years, the number of intravenous infusions has been considerably reduced by combining chemotherapeutic drugs for trypanosome infections, thereby enhancing patients’ quality of life.[26–28]

Current efforts to improve infection diagnosis focus on serological methods and antibody detection. Numerous lateral flow tests based on singular parasite-derived VSGs have recently been developed.[29] A dual-antigen lateral flow test using sVSG117 in combination with a cell lysate protein, rISG65,[30] as immunodiagnostic antigens were reported for detection of trypanosome infections in humans with good specificity.[31] Considering heterogeneity-based antigenic variation, this study suggests sVSG117 as either a dominant mother gene in segmental gene conversion and mosaicism or an existing reactive epitope at the C-terminal domain of the VSGs.

In contrast to VSGs, the structure of GPIs only depends on the glycosylation machinery of the cell. The structural variations include the presence or absence of galactoses attached to the conserved GPI pseudopentasaccharide core (Figure 1a).[32] We recently described the synthesis of diverse GPI glycolipids using a general convergent strategy and a set of fully orthogonal protecting groups.[33] We expanded this strategy to obtain a series of galactosylated GPI fragments from the GPI of *Trypanosoma brucei*.[34,35] Introduction of α-galactoses required additional ester-based protecting groups and adapted synthetic strategies. Synthetic GPI derivatives have shown potential application in serodiagnosis by determining anti-GPI antibodies in patients infected with *Toxoplasma gondii* and *Plasmodium falciparum*.[36–39]

**Figure 1:**
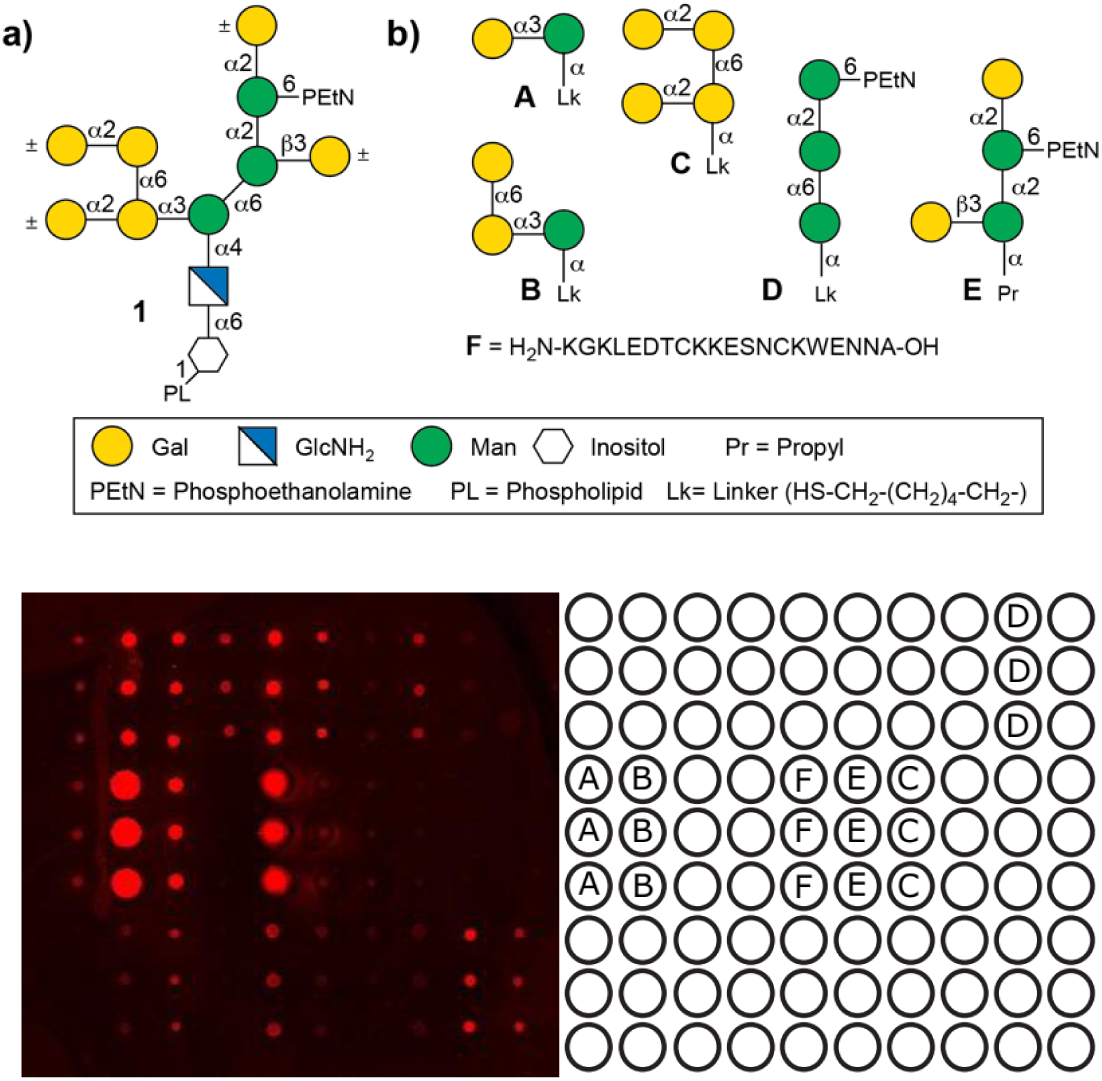
*Trypanosoma brucei* GPI derivatives: a) Representation of the *T. brucei* GPI; b) *T. brucei* GPI fragments containing mannose with different galactosylation patterns **A**-**E**; c) low variant CTD-peptide of VSG117 (**F)**; d) Representative Scan of a microarray indicating the positions of structures **A** - **F**; and e) Glycan-array printing pattern

We hypothesized that the host immune system recognizes the C-terminal of VSGs and galactosylated structures of the trypanosome GPI glycolipid, inducing specific antibodies that bind to synthetic structures and can be used to detect an infection. The main advantage of investigating chemically synthesized GPI derivatives over VSGs derived from parasitic cultures is the higher accessibility and availability of greater homogeneous amounts. Here, we evaluate a series of synthetic GPI fragments with specific modifications of the natural *T. brucei* GPIs (Structures **A**-**F**, Figure 1b) to determine HAT infections. We printed the synthetic compounds on glass slides and used the obtained glycan array to detect anti-GPI antibodies in the sera of mice infected with trypanosome parasites. We show that GPI glycan-specific recognition depends on the presence of α-galactose and demonstrate that these structures can be used for the detection of short-term IgM and long-term IgG. Furthermore, we performed an analysis of sera from *T. brucei gambiense* and *T. brucei rhodesiense* infected humans using the GPI array and demonstrated for the first time the presence of specific antibodies recognizing synthetic GPI glycans and a peptide present in the VSG-GPI interface.

## Materials and Methods

### General Synthetic Methods

All chemicals were reagent grade and all solvents anhydrous high-purity grade and used as supplied except where noted otherwise. Unless noted otherwise, reactions were performed in oven-dried glassware under an inert argon atmosphere. Reaction molarity was 0.1 molar except where noted otherwise. Reagent-grade thiophene was dried over activated molecular sieves before use. Pyridine was distilled over CaH_2_ before use. Sodium hydride suspension was washed with hexane and THF and stored in an anhydrous environment. Benzyl bromide was passed through activated basic aluminum oxide before use. Before use, molecular sieves were powdered and activated by heating under a high vacuum. Analytical thin layer chromatography (TLC) was performed on Merck silica gel 60 F_254_ plates (0.25mm). All compounds were visualized by UV irradiation and/or heating the plate after dipping into a staining solution. The compounds were stained with cerium sulfate-ammonium molybdate (CAM) solution, basic potassium permanganate solution, acidic ninhydrin-acetone solution or a 3-methoxyphenol-sulfuric acid solution. Flash column chromatography was carried out using a forced flow of the indicated solvent on Fluka silica gel 60 (230-400 mesh, for preparative column chromatography).

^1^H, ^13^C and ^31^P-NMR spectra were recorded on a Varian 400 (400 MHz), a Varian 600 (600 MHz), a Bruker 400 (400 MHz) and a Bruker Ascend 400 (400 MHz) spectrometer in CDCl_3_ (7.26 ppm ^1^H, 77.1 ppm ^13^C), D_2_O (4.79 ppm ^1^H), MeOD (4.87 ppm and 3.31 ppm ^1^H, 49.00 ppm ^13^C), acetone-d6 (2.05 ppm and 2.84 ppm ^1^H, 206.26 ppm and 29.84 ppm ^13^C) unless otherwise stated. Coupling constants are reported in Hertz (Hz). Splitting patterns are indicated as s, singlet; d, doublet; t, triplet; q, quartet; br, broad singlet; dd, doublet of doublets; m, multiplet; dt, doublet of triplets; h, sextet for ^1^H NMR data. Signals were assigned using ^1^H-^1^H COSY, ^1^H-^1^H TOCSY, ^1^H-^1^H NOESY, ^1^H-^1^H ROESY, ^1^H-^13^C HSQC, ^1^H-^13^C HMBC spectra and version thereof. ESI mass spectral analyses were performed by the MS-service at the Institute for Chemistry and Biochemistry at the Free University of Berlin using a modified MAT 711 spectrometer, the MS-service at the Institute for Chemistry at the University of Potsdam using an ESI-Q-TOF micro spectrometer and a Waters Xevo G2-XS QTof coupled with an Acquity H-class UPLC. Infrared (FTIR) spectra were recorded as thin films on a Perkin Elmer Spectrum 100 FTIR spectrophotometer. Optical rotations were measured with a Schmidt & Haensch UniPol L 1000 at a concentration (c) expressed in g/100 mL. HPLC-supported purifications were conducted using Agilent 1100 and Agilent 1200 systems. Supercritical fluid chromatography was carried out using a Waters Investigator System.

### Ethics statement

All experiments concerning the mice complied with the ECPVA guidelines (CETS n° 123) and were approved by the VUB Ethical Committee (Permit Number: 14-220-29). Human serum samples used for this research were obtained from the WHO HAT Specimen Biobank and stored at the Pasteur Institute. WHO acquired patient consent, and serum samples were collected to develop new diagnostic tests.[40]

### Patient sera

Patient infection status was determined by applying the CATT test, subsequent parasitological analysis and examination of clinical symptoms of HAT.[40] Serum samples were stored in the WHO HAT Specimen Biobank at −80°C, shipped to Potsdam on dry ice, divided into aliquots and stored at −20°C.

### Parasites, mice and infections

Clonal pleomorphic *T. brucei* AnTat 1.1E parasites were a kind gift from N. Van Meirvenne (Institute for Tropical Medicine, Belgium) and stored at −80°C. Female wild-type (WT) C57Bl/6 mice (7–8 weeks old) were obtained from Janvier and infected with 5×10^3^ AnTat1.1E trypanosomes (intraperitoneally (i.p.) in 200 µL HBSS (Hanks’ balanced salt solution, ThermoFisher Scientific).

### Mice Sera

Blood was collected from CO_2_ euthanized non-infected and *T.brucei* infected mice via cardiac puncture, centrifuged (15 minutes, 10.000xg, 4°C), and serum was kept at -80°C.

### Preparation of glycan microarray

The synthetic glycans were dissolved in sodium phosphate buffer (50 mM, pH 8.5 for amine linker compounds) or PBS buffer (pH 7.4 for thiols, including an equimolar amount of TCEP·HCl). The compounds were immobilized in four copies employing a piezoelectric spotting device (S3, Scienion) on maleimide-functionalized slides or epoxy slides (sciCHIPEPOXY, Scienion), in 50% relative humidity at 23°C. The printed slides were stored for 18 h in a humidified chamber to complete the immobilization reaction. Afterwards, the slides were stored in a cooled environment. Before the experiment, the slides were washed three times with water, and the remaining maleimide or epoxy groups were quenched by incubating the slides in an aqueous solution of 100 mM ethanolamine in sodium phosphate buffer (50 mM, pH 9.01) for 1 h at 25°C. The slides were rinsed three times with water and dried by centrifugation. Microarrays were blocked with BSA (2.5%, w/v) in PBS for 1 h at room temperature. Blocked slides were washed twice with PBS, centrifuged and incubated with a 1:15 dilution of mouse or human sera in PBS for 1 h. After washing with PBS, microarrays were incubated with goat anti-mouse IgG H+L Alexa 645 (Molecular Probes, 1:400), donkey anti-mouse IgM Alexa 594 (Dianova, 1:200), goat anti-human IgG-Fc Alexa488 (Dianova, 1:400) or goat antihuman IgM Alexa 594 (Molecular Probes, 1:200) in PBS containing 1% BSA for 1 h. The slides were then washed with PBS and double-distilled water, subsequently dried by centrifugation and analyzed using a fluorescence microarray scanner (Genepix® 4300A, Molecular Devices).

### Data processing and Statistical analysis

Data were imported to GenePix Pro, and a mask was superimposed on the fluorescent area separating it from the background. The difference between fluorescence (inside) and background (outside) was calculated for each position and normalized against the mean of all samples corresponding to the same synthetic structure. The resulting data of duplicate scans were imported to GraphPad Prism, and outliers were removed. ANOVA with multiple comparisons and an initial significance test by Tukey was performed. Then, an unpaired T-test with the following definition of significance was performed: not significant (ns) = p > 0,05; * = p < 0,05; ** = p < 0,01; *** = p < 0,001; **** = p < 0,0001. Receiver operating characteristic (ROC) curves were generated when p < 0,05 existed for a synthetic structure in either murine IgM or IgG scan and for one disease stage for human IgM or IgG infected with *Tbg* and *Tbr,* respectively.

## Results

### Selection of synthetic VSG-GPI fragments

We previously reported on the synthesis of a series of synthetic fragments of the GPI of *T. brucei*.[34] We used the synthetic precursors of these molecules to obtain a second series of fragments with specific GPI modifications and a linker for immobilization and production of glycan microarray (Figure 1a-c, SI). The most prominent structural feature in GPIs from *T. brucei* is the presence of α-galactosylation on the C-3 position of Man-I. Interestingly, GPIs of VSG117, VSG221 and VSG121 variants bear additional galactosylation of up to two galactosides along a conserved 1→6 linkage.[41] Consequently, these modifications were essential for designing the substructures **A-C** to perform a GPI epitope mapping. The trimannose (**D)** was designed to lack the mammalian phosphorylation of Man-I at the C2 position to mimic the oligomannose part present in parasitic GPIs and to complete the variable part of GPIs. Earlier reports suggested that the absence of this phosphorylation leads to recognition by antibodies generated during an immune response.[36,37] As a final GPI fragment, the tetrasaccharide **E**,[35] was considered with two galactose residues to cover the heterogeneity of trypanosome VSG 221 and VSG121.[34,41] A final structure, the peptide (KGKLEDTCKKESNCKWENNA) **F** was designed to cover the VSG117 CTD that connects the protein to GPI and is also a part with expected low structural diversity.[31]

### IgM and IgG antibodies derived from *Trypanosoma brucei-*infected mice specifically recognize synthetic GPI fragments

Glycan microarrays using the synthetic compounds were prepared and incubated first with sera derived from 10 naïve and 40 AnTat1.1E infected C57Bl/6 mice. Glycan recognition analysis using a fluorescent secondary antibody (Figure 1c) showed low fluorescence levels for all antibody classes, indicating low anti-glycan antibody levels (SI). A quantification and statistical evaluation showed a three-fold increase in fluorescence level for IgM and IgG antibodies from infected mice sera at days 7, 14, 21, and 28 post infection (Figure 2, SI). These findings indicated the presence of antibodies recognizing GPI structures from VSGs.[7] Further data evaluation was necessary to determine whether synthetic **A**-**F** structures are antigens suitable for detecting a trypanosome infection.

**Figure 2:**
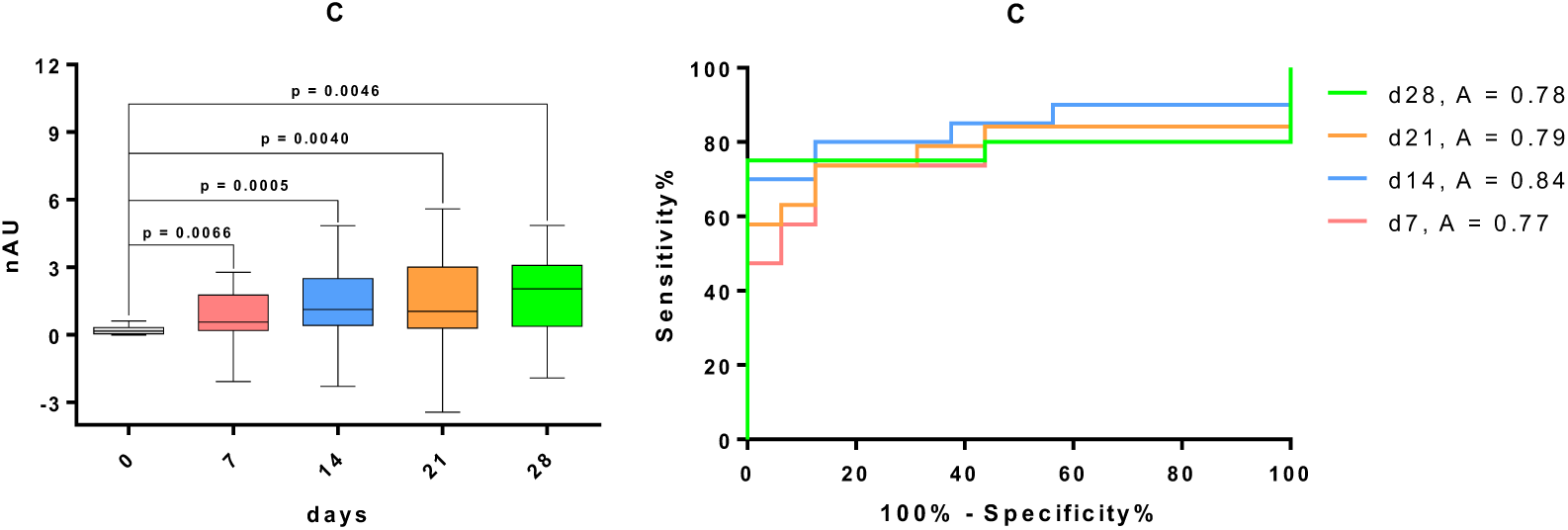
Statistical evaluation of fluorescence levels obtained by the interaction of IgM antibodies in mice sera with synthetic structure C. : *left*: Box-Plot indicating an increase in GPI fragment recognition after infection; *right:* ROC curve for the same synthetic structure

First, the grouped IgM response was analyzed against the naïve control using ANOVA and an unpaired t-test (Figure 2, SI). For structures showing significant recognition, a receiver operating characteristic (ROC) curve and the corresponding confidence intervals were calculated for the whole sera set (Figure 2, SI). ROC curves of **A** (average A = 0.79), **C** (A = 0.80) and **D** (A = 0.75) clearly showed these structures as potential markers for diagnosis. The in-lab trial test sensitivity and specificity determined for **A** (74.85%/94.74%), **C** (66.51%/ 93.75%) and **D** (66.84%/94.74%) are strong and suggest a high probability ratio for distinguishing infected and healthy specimen by serological determination of specific anti-GPI antibodies of trypanosome-infected mice (SI).

Synthetic glycan recognition by IgG was similarly analyzed (SI). The ROC analysis of this study delivered significant values for **A** (average A = 0.70; 61.20%/95%) and **D** (A = 0.83; 76.20%/94.44%), confirming their suitability to be recognized by antibodies present only in sera of the infected specimens (SI). With these promising results, we were motivated to investigate whether these results were transferable to the analysis of patient sera and endemic controls from sub-Saharan Africa and lead to the diagnosis of Human African Trypanosomiasis based on antibody binding of small trypanosome surface antigens.

### Synthetic GPI fragments are diagnostic antigens of Human African Trypanosomiasis

Sera from *T. brucei ghambiense* (Tbg) and *T. brucei rhodesiense* (Tbr) infected humans were provided by the WHO from the trypanosome database at Institute Pasteur.[40] Datasets comprised ten samples each for endemic control, a stage 1 and a stage 2 of the two distinct parasite infections. Initial observation showed higher structure recognition and fluorescence levels than in the mice sera analysis. A five-fold fluorescence increase between endemic controls and infected specimen indicated recognition of synthetic GPI structures by human antibodies (Figure 3). Analogous to analysis of mice sera, the IgM antibodies levels where statistically evaluated and allowed stage independent detection of *Tbg* infection by recognition of structures **A** (stage 1: A = 0.82; 63.16%/94.84%; stage 2: A = 0.89; 64.71%/94.74%), **C** (stage 1: A = 0.85; 52.63%/94.44%; stage 2: A = 0.82; 65.00%/94.44%), **E** (stage 1: A = 0.81; 65.00%/94.12%; stage 2: A = 0.87; 80.00%/94.12%) and **F** (stage 1: A = 0.81; 35.00%/95.00%; stage 2: A = 0.81; 55.00%/90.00%) (Figure 4, Table 3, SI). In contrast, IgM-mediated detection of *Tbr* infection was only possible in sera from patients in the second disease stage, with glycan **A** (A = 0.72; 44.44%/95.00%) and **C** (A = 0.82; 57.89%/94.44%) again giving the best results.

**Figure 3:**
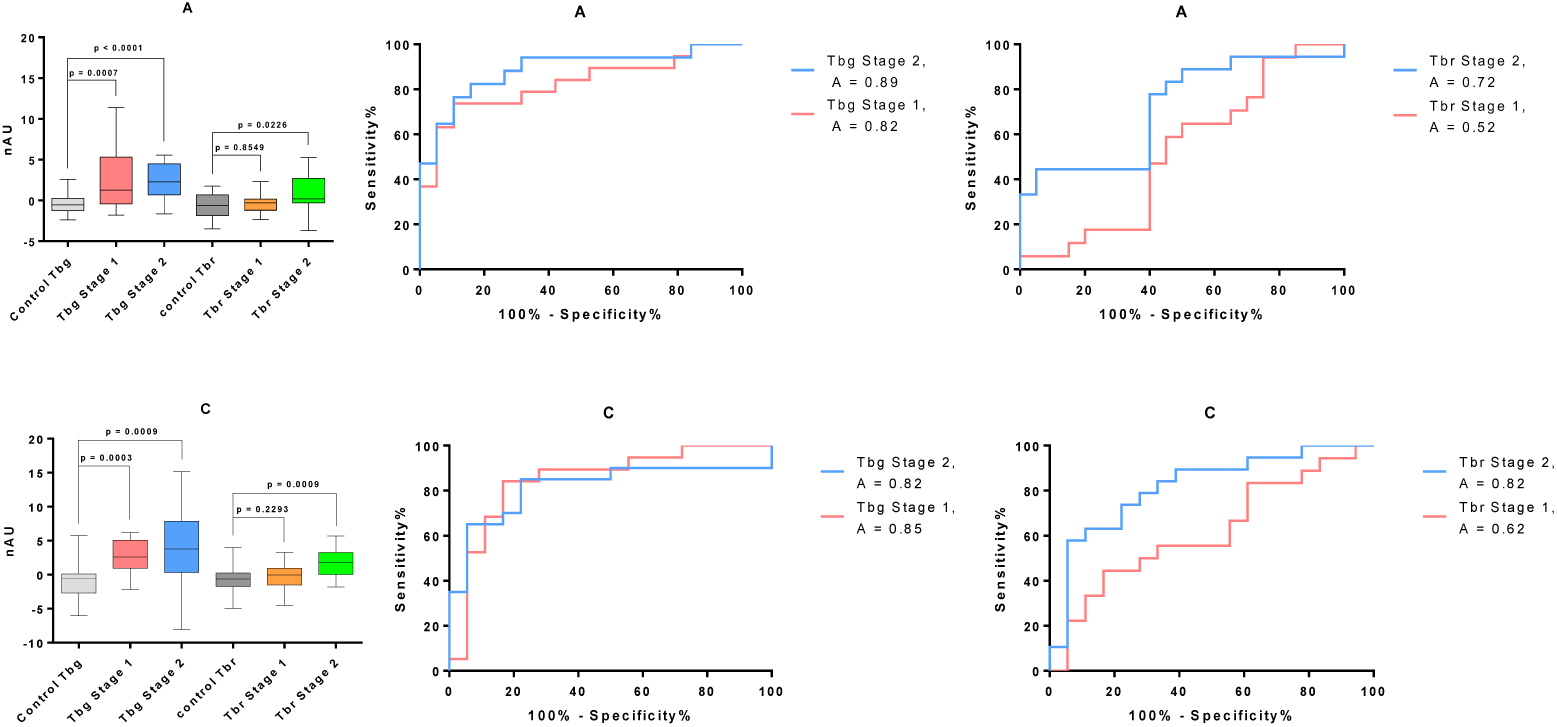
Statistical evaluation of fluorescence levels caused by interaction of IgM originating from human sera with an infection of *Tbg* or *Tbg* and endemic control with synthetic structures A and C: 1^st^ row: Box-Plot and ROC curves indicating an increase in GPI fragment **A** recognition in specimen with *Tbg* and a weaker recognition with focus on stage 2 with *Tbr* infection; 2^nd^ row: GPI fragment **C**

**Table 1:** Sensitivity, specificity, corresponding confidence intervals and alternative likelihood ratios of a test using IgM originating from human sea with *Tbg* or *Tbr* infection and endemic control and synthetic substances A and C.

| Structure/Stage |  | Sensitivity% | 95% CI | Specificity% | 95% CI | Likelihood ratio |
| --- | --- | --- | --- | --- | --- | --- |
| Tb gambiense |  |  |  |  |  |  |
| A | Stage 1 | 63.16 | 38.36% to 83.71% | 94.74 | 73.97% to 99.87% | 12.00 |
|  | Stage 2 | 64.71 | 38.33% to 85.79% | 94.74 | 73.97% to 99.87% | 12.29 |
| C | Stage 1 | 52.63 | 28.86% to 75.55% | 94.44 | 72.71% to 99.86% | 9.47 |
|  | Stage 2 | 65.00 | 40.78% to 84.61% | 94.44 | 72.71% to 99.86% | 11.70 |
| Tb rhodesiense |  |  |  |  |  |  |
| A | Stage 1 | 52.94 | 27.81% to 77.02% | 55.00 | 31.53% to 76.94% | 1.18 |
|  | Stage 2 | 44.44 | 21.53% to 69.24% | 95.00 | 75.13% to 99.87% | 8.89 |
| C | Stage 1 | 22.22 | 6.409% to 47.64% | 94.44 | 72.71% to 99.86% | 4.00 |
|  | Stage 2 | 57.89 | 33.50% to 79.75% | 94.44 | 72.71% to 99.86% | 10.42 |

**Figure 4:**
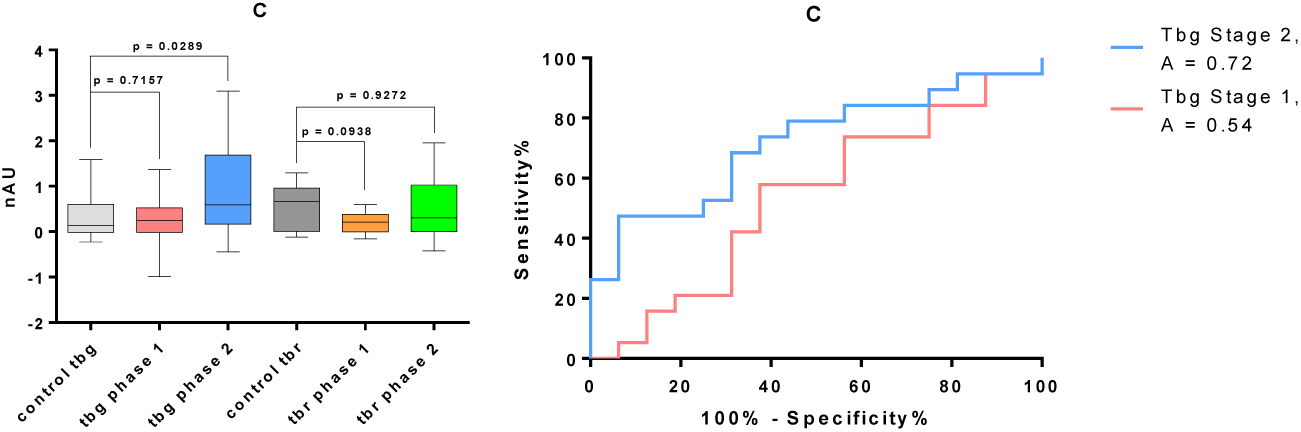
Statistical evaluation of fluorescence levels caused by interaction of IgG originating from human sera with an infection of *Tbg* and endemic control with synthetic structure C: Box-Plot and ROC curves indicating an increase in GPI fragment **C** recognition in specimen with *Tbg* phase 2 infection

**Table 2:** Sensitivity. specificity. corresponding confidence intervals and alternative likelihood ratios of a test using IgG originating from human sea with *Tbg* infection and endemic control and synthetic substance C.

| Structure/Stage |  | Sensitivity% | 95% CI | Specificity% | 95% CI | Likelihood ratio |
| --- | --- | --- | --- | --- | --- | --- |
| *Tb gambiense* |  |  |  |  |  |  |
| C | Stage 1 | 57.89 | 33.50% to 79.75% | 62.50 | 35.43% to 84.80% | 1.54 |
|  | Stage 2 | 47.37 | 24.45% to 71.14% | 93.75 | 69.77% to 99.84% | 7.58 |

An analysis of the glycan recognition by IgG antibody to detect *Tbg* of infections showed antigens **C** (A = 0.72; 47.37%/93.75%) and **E** (A = 0.75; 60.00%/93.75%) as the best structures to distinguish infected from healthy individuals in disease stage 2 only (Supporting Info). None of the synthetic antigens synthesized in this study showed a potential in detecting infection of *Tbr* based on anti-glycan IgG determinations.

## Discussion and Conclusion

In contrast to other protozoan parasites presenting free GPIs on the cell membrane. *T. brucei* GPIs are covered by a dense layer of glycoproteins.[42] Thus, it was essential to determine whether a host immune system can induce an immune response against the buried glycolipid anchor and which structural domains are involved. This study focused on determining antibodies specific to galactosylated GPI structures, a *T. brucei GPI-*specific modification.[34,35] VSG-GPI conjugates may be released by phosphoinositide phospholipase C-mediated hydrolysis or during phagocytosis. While only a limited number of *T. brucei* GPI anchors exist,[41] diverse VSG sets are processed[15] in both mechanisms. These differences may result in a polyclonal immune response to the few GPI modifications and allow diagnosis of a trypanosome infection. We evaluated this response by printing glycan microarrays and determining the recognition by antibodies in sera.

Antibodies from *T. brucei-infected* mice recognized the small synthetic GPI structures. Sera of infected mice presented anti-glycan IgM antibodies binding the structures **A, C** and **D.** The IgM levels were suitable to distinguish sera of naïve mice from infected animals several weeks after infection (Figures 2 and 3). The infection occurred in four groups ranging from one to four weeks before serum collection. Thus, the murine immune response likely encountered many VSG-GPI variants but only beginning impairment of B cell maturation.[15] Evaluation of the response by IgG antibodies showed the recognition of structures **A** and **D** and a distinction between infected and healthy specimens. The sensitivity for both IgM and IgG antibodies was comparable and moderate, with values between 66.51% - 74.85% for IgM and 61.20% - 76.20% for IgG. Although these levels are commonly reached in early-stage antibody-based diagnosis of trypanosome infection,[30,31] the specificity observed in our test reaches current field standards (85 - 97% for CATT, 96 - 99% LATEX).[45, 49] Synthetic structures **A, C** and **D** show specificity values between 93.75% - 94.74% for IgM and 94.44% - 95.00% for IgG. The overlap of the structures giving the best serostatus differentiation suggests using galactosylated structures and the non-mammalian GPI backbone to detect trypanosome infections in mice.

Encouraged by these results, the glycan array of structures **A-F** was used to screen a set of human sera from endemic controls and *Tbg* and *Tbr-infected* patients in disease stages 1 and 2. Sera from *Tbg* patients showed IgM antibody recognition of structures **A, C, E** and **F**. suggesting them as antigens for diagnosis independent of the infection stage. The detection was of moderate sensitivity between 35% - 80% and high specificity between 90% - 95%. In contrast, only compounds **A** and **C** were diagnostic antigens for infections with *Tbr* and only in the second disease stage. The test sensitivity was low compared to IgM, with values between 44% and 58%, but it maintained high specificity between 94.5% and 95.0%. Using IgG as a readout for *Tbg* infection, substances **C** and **E** were applicable with sensitivity between 47% and 67% and a specificity of 94%.

The results suggest applying fully synthetic GPI structures as diagnostic antigens of HAT, especially caused by *Tbg* infections. These preliminary data and the recognition of multiple structures indicate the possibility of improving the test sensitivity by structure optimization. In contrast, the high test specificity indicates the suitability of GPI antigens to exclude false negatives. Summarizing all different analyses, the tetra-α-galactoside structure **C** emerges as the most suitable structure for diagnosing infected individuals from distinct geographical areas. Analyzing structures **A** and **C,** it becomes apparent that **C** bears the terminal α-galactose structure twice. Thus, it can be seen as a multivalent display of this precursor structure **A** with enhanced binding affinity.

Further, the positive results obtained with compound **D** in the analysis of human sera do not necessarily reflect an exclusive response against trypanosome-derived GPI. This structure is conserved in other protozoan parasites, such as *Toxoplasma gondii* and *Plasmodium falciparum,* and the binding may derive from cross-reactivity with antibodies from other infections.[25.27] Further characterization of the sera and record of previous patient infections should help to clarify the recognition’s origin.

Compound **E** was immobilized using the phosphoethanolamine moiety; consequently, the structure was closer to the glass surface and reversed to the natural orientation of GPIs on the cell membrane. This orientation did not affect the binding by antibodies, and a broad recognition by antibodies was still observed. These results suggest a certain level of VSG-GPI release through hydrolysis, a phagocytosis mechanism or overlapping epitope recognition. Future applications in a field trial may prioritize diagnosis of infection over stage determination. Under these conditions, compounds **A** and **C** are key to determining a serostatus and IgM-based HAT diagnosis. One limitation of the analysis performed in this study is the number of sera used. Future investigations may elaborate on this test and increase the subject number to overcome the sensitivity limitations. Establishing a diagnostic test based on fully synthetic antigens may replace serological tests using recombinant antigens and parasite cultivation-dependent requirements.[48,49]

## Authors Contributions

Conceived and designed experiments: MM, BS, DM, AG; performed experiments: MM, BS, MG, DM, AG; analyzed data: MM, DM, AG; contributed reagents, materials, and analysis tools: MM, BS, MG, DM, AG, PHS, DVS; wrote the manuscript: MM, DM, DVS. Supervised the project PHS and DVS.

## Acknowledgments

We thank the WHO and the ICAReB platform in the Institute Pasteur for providing samples of human specimen. This project was supported by the Max Planck Society and the RIKEN-Max Planck Joint Research Center for Systems Chemical Biology. Dr. Maurice Michel is thankful for a doctoral scholarship from the Beilstein-Institut zur Förderung der chemischen Wissenschaften and a scholarship from the federal state of Lower Saxony. Prof. Dr. Benoît Stijlemans was supported by the Strategic Research Program (SRP#47. VUB). Dr. Monika Garg received a doctoral scholarship from the International Max Planck Research School on Multiscale Biosystems of the MPIKG. We thank Olaf Niemeyer for NMR Service, Eva Settels for technical support and Dr. Martin Scobie for carefully proofreading the manuscript.

## Supporting Information

### General Synthetic Methods

All chemicals were reagent grade and all solvents anhydrous high-purity grade and used as supplied except where noted otherwise. Reactions were performed in oven-dried glassware under an inert argon atmosphere unless noted otherwise. Reaction molarity was 0.1 molar except where noted otherwise. Reagent grade thiophene was dried over activated molecular sieves prior to use. Pyridine was distilled over CaH_2_ prior to use. Sodium hydride suspension was washed with hexane and THF and stored in an anhydrous environment. Benzyl bromide was passed through activated basic aluminum oxide prior to use. Molecular sieves were powdered and activated by heating under high vacuum prior to use. Analytical thin layer chromatography (TLC) was performed on Merck silica gel 60 F_254_ plates (0.25mm). All compounds were visualized by UV irradiation and/or heating the plate after dipping into a staining solution. The compounds were stained with cerium sulfate-ammonium molybdate (CAM) solution, basic potassium permanganate solution, acidic ninhydrin-acetone solution or a 3-methoxyphenol-sulfuric acid solution. Flash column chromatography was carried out using a forced flow of the indicated solvent on Fluka silica gel 60 (230-400 mesh, for preparative column chromatography).

^1^H, ^13^C and ^31^P-NMR spectra were recorded on a Varian 400 (400 MHz), a Varian 600 (600 MHz), a Bruker 400 (400 MHz) and a Bruker Ascend 400 (400 MHz) spectrometer in CDCl_3_ (7.26 ppm ^1^H, 77.1 ppm ^13^C), D_2_O (4.79 ppm ^1^H), MeOD (4.87 ppm and 3.31 ppm ^1^H, 49.00 ppm ^13^C), acetone-d6 (2.05 ppm and 2.84 ppm ^1^H, 206.26 ppm and 29.84 ppm ^13^C) unless otherwise stated. Coupling constants are reported in Hertz (Hz). Splitting patterns are indicated as s, singlet; d, doublet; t, triplet; q, quartet; br, broad singlet; dd, doublet of doublets; m, multiplet; dt, doublet of triplets; h, hextet for ^1^H NMR data. Signals were assigned by means of ^1^H-^1^H COSY, ^1^H-^1^H TOCSY, ^1^H-^1^H NOESY, ^1^H-^1^H ROESY, ^1^H-^13^C HSQC, ^1^H-^13^C HMBC spectra and version thereof. ESI mass spectral analyses were performed by the MS-service at the Institute for Chemistry and Biochemistry at the Free University of Berlin using a modified MAT 711 spectrometer, the MS-service at the Institute for Chemistry at the University of Potsdam using an ESI-Q-TOF micro spectrometer and a Waters Xevo G2-XS QTof coupled with an Acquity H-class UPLC. Infrared (FTIR) spectra were recorded as thin films on a Perkin Elmer Spectrum 100 FTIR spectrophotometer. Optical rotations were measured with a Schmidt & Haensch UniPol L 1000 at a concentration (c) expressed in g/100 mL. HPLC supported purifications were conducted using Agilent 1100 and Agilent 1200 systems. Supercritical fluid chromatography was carried out using a Waters Investigator System.

### Parasites, mice and infections

Clonal pleomorphic *T. brucei* AnTat 1.1E parasites were a kind gift from N. Van Meirvenne (Institute for Tropical Medicine, Belgium) and stored at −80°C. mice. Female wild-type (WT) C57Bl/6 mice (7–8 weeks old) were obtained from Janvier and infected with 5×10^3^ AnTat1.1E trypanosomes (intraperitoneally (i.p.) in 200 µL HBSS (Hanks’ balanced salt solution, ThermoFisher Scientific).

### Sera isolation

Blood was collected from CO_2_ euthanized non-infected and *T.brucei* infected mice via cardiac puncture, centrifuged (15 minutes, 10.000xg, 20°C), and serum was kept at -80°C. Human sera were obtained from the WHO and the ICAReB platform at the Institute Pasteur.

### Ethics statement

All experiments complied with the ECPVA guidelines (CETS n° 123) and were approved by the VUB Ethical Committee (Permit Number: 14-220-29).

#### PREPARATION OF GLYCAN MICROARRAY

The synthetic glycans were dissolved in phosphate buffer (50 mM NaH2PO4, pH 8.5 for amine linker compounds) or PBS buffer (pH 7.4 for thiols, including an equimolar amount of TCEP·HCl). The compounds were immobilized in four copies employing a piezoelectric spotting device (S3, Scienion) on maleimide-functionalized slides or on epoxy slides (sciCHIPEPOXY, Scienion), in 50% relative humidity at 23°C. The printed slides were stored for 18 h in a humified chamber to complete the immobilization reaction. Afterwards the slides were stored in a cooled environment. Prior to the experiment, the slides were washed three times with water and the remaining maleimido or epoxy groups were quenched by incubating the slides in an aqueous solution of 100 mM ethanolamine and 50 mM Na2HPO4·12H2O with pH 9.01 for 1 h at 25°C. The slides were rinsed three times with water and dried by centrifugation. Microarrays were blocked with BSA (2.5%, w/v) in PBS for 1 h at room temperature. Blocked slides were washed twice with PBS, centrifuged and incubated with a 1:15 dilution of mouse or human sera in PBS for 1 h. After washing with PBS, microarrays were incubated with goat anti-mouse IgG H+L Alexa 645 (Molecular Probes, 1:400), donkey anti-mouse IgM Alexa 594 (Dianova, 1:200), goat anti-human IgG-Fc Alexa488 (Dianova, 1:400) or goat antihuman IgM Alexa 594 (Molecular Probes, 1:200) in PBS containing 1% BSA for 1 h. The slides were then washed with PBS and double-distilled water, subsequently dried by centrifugation and analyzed using a fluorescence microarray scanner (Genepix® 4300A, Molecular Devices).

### Data processing and Statistical analysis

Data were imported to GenePix pro and a mask was superimposed on the fluorescent area separating it from the background. Difference of fluorescence (inside) and background (outside) was calculated for each position and normalized against the mean of all samples corresponding to the same synthetic structure. The resulting data of duplicate scans were imported to GraphPad Prism and outliers removed. ANOVA with multiple comparisons and an initial significance test by Tukey was performed. Then an unpaired T-test with the following definition of significance was performed: not significant (ns) = p > 0,05; * = p < 0,05; ** = p < 0,01; *** = p < 0,001; **** = p < 0,0001. Receiver operating characteristic (ROC) curves were generated, when p < 0,05 existed for at least three groups of one synthetic structure in either murine IgM or IgG scan and for one disease stage for human IgM or IgG infected with Tbg and Tbr respectively. Sensitivity and Specificity are shown for the scenario of the highest likelihood ratio, *i.e.* how much more is it likely, that a specimen with a positive test has the disease versus a person with a negative test having the disease. The likelihood ratio equals sensitivity/(1.0-specificity).

### Compound printing pattern

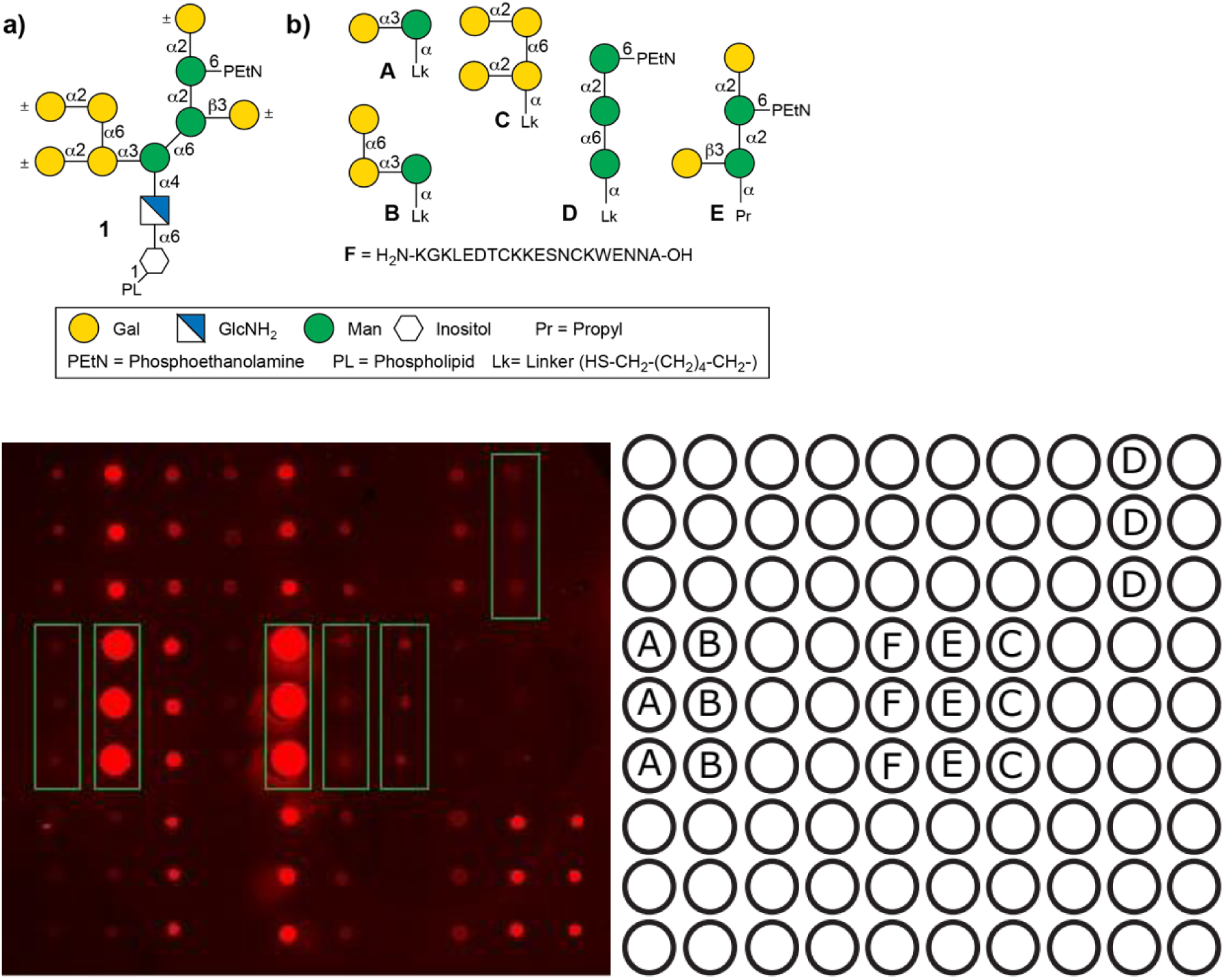

### Readout Microarray

#### IgM Mouse

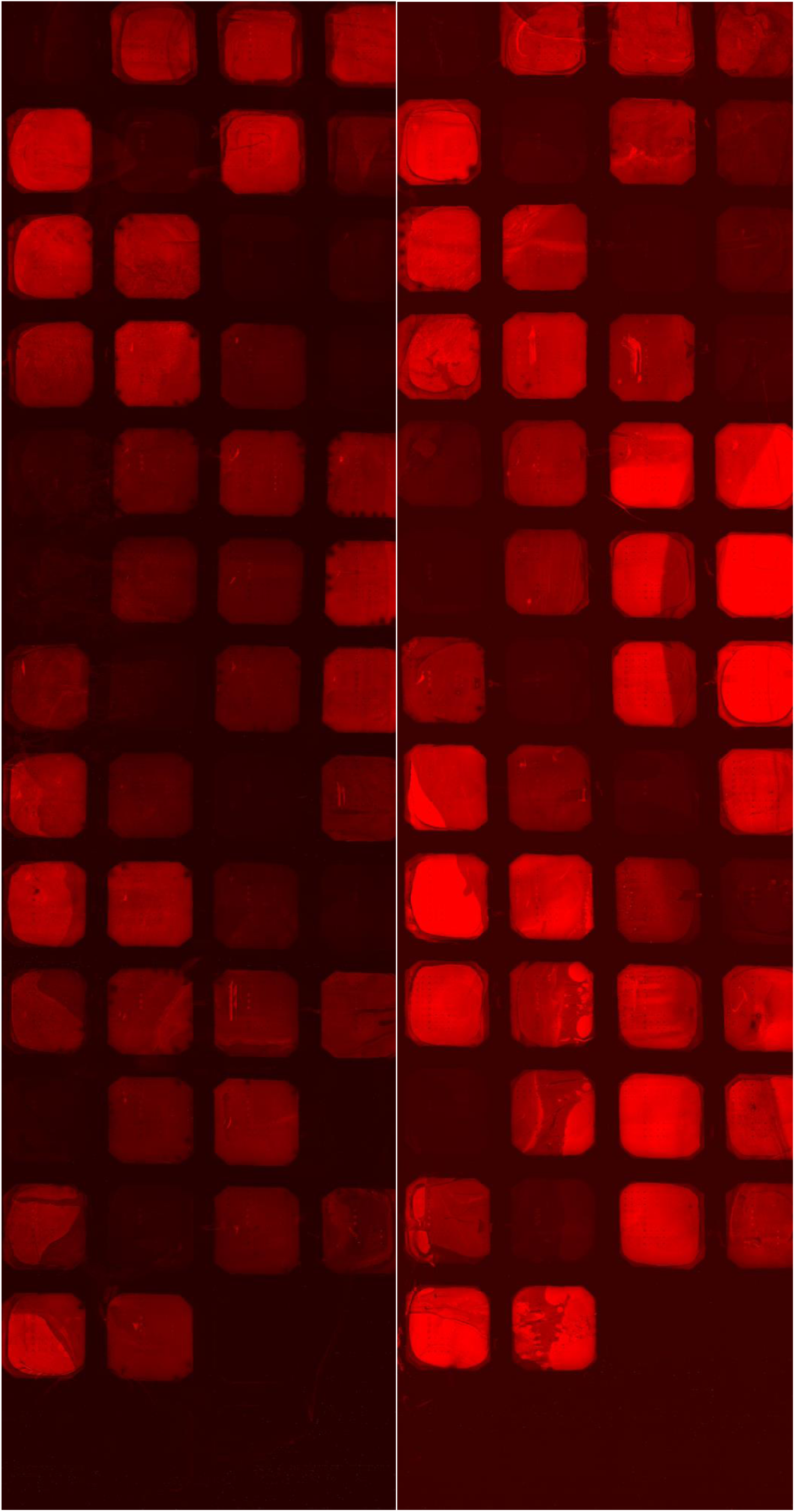

#### IgG Mouse

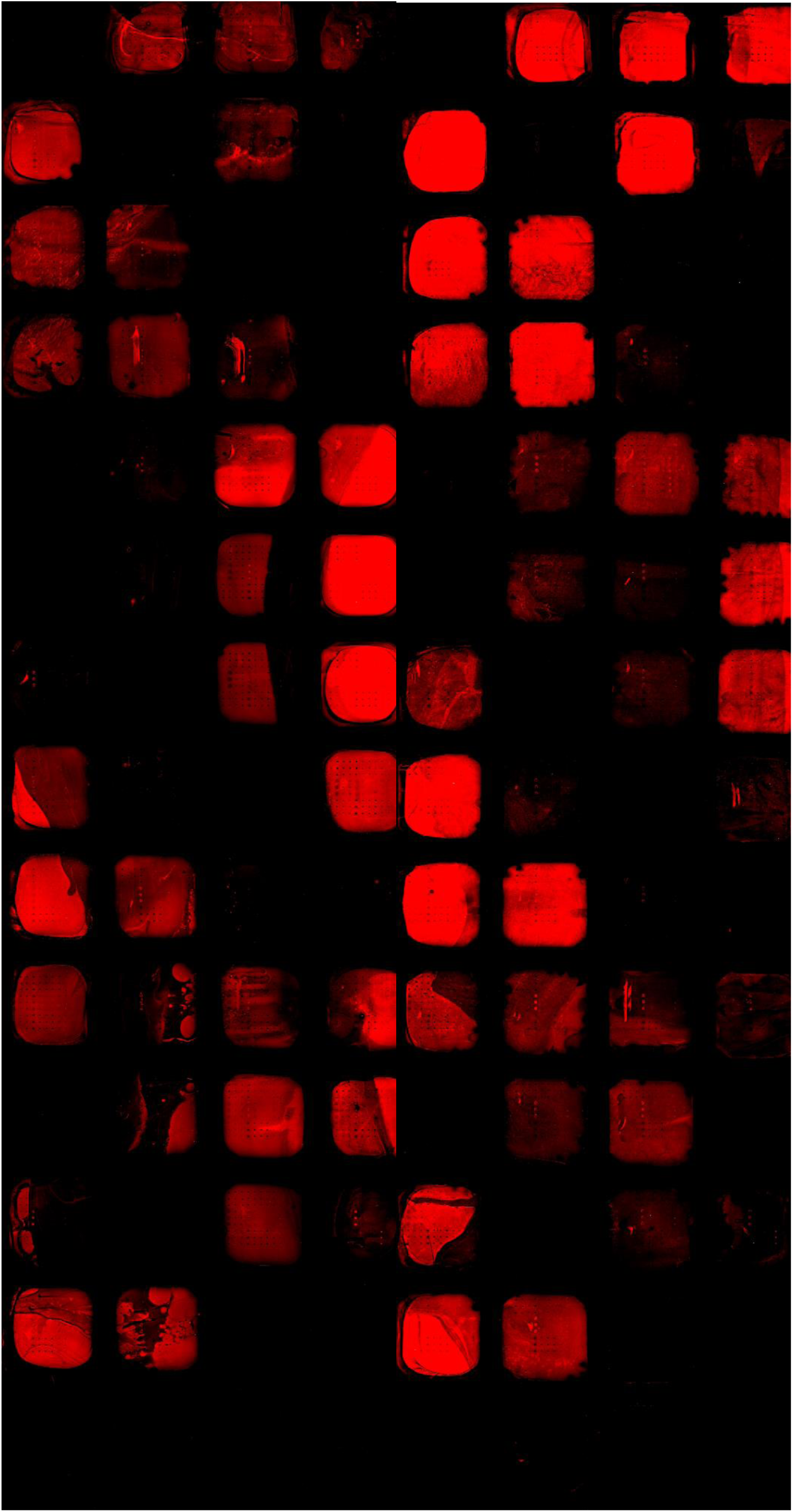

#### IgM human

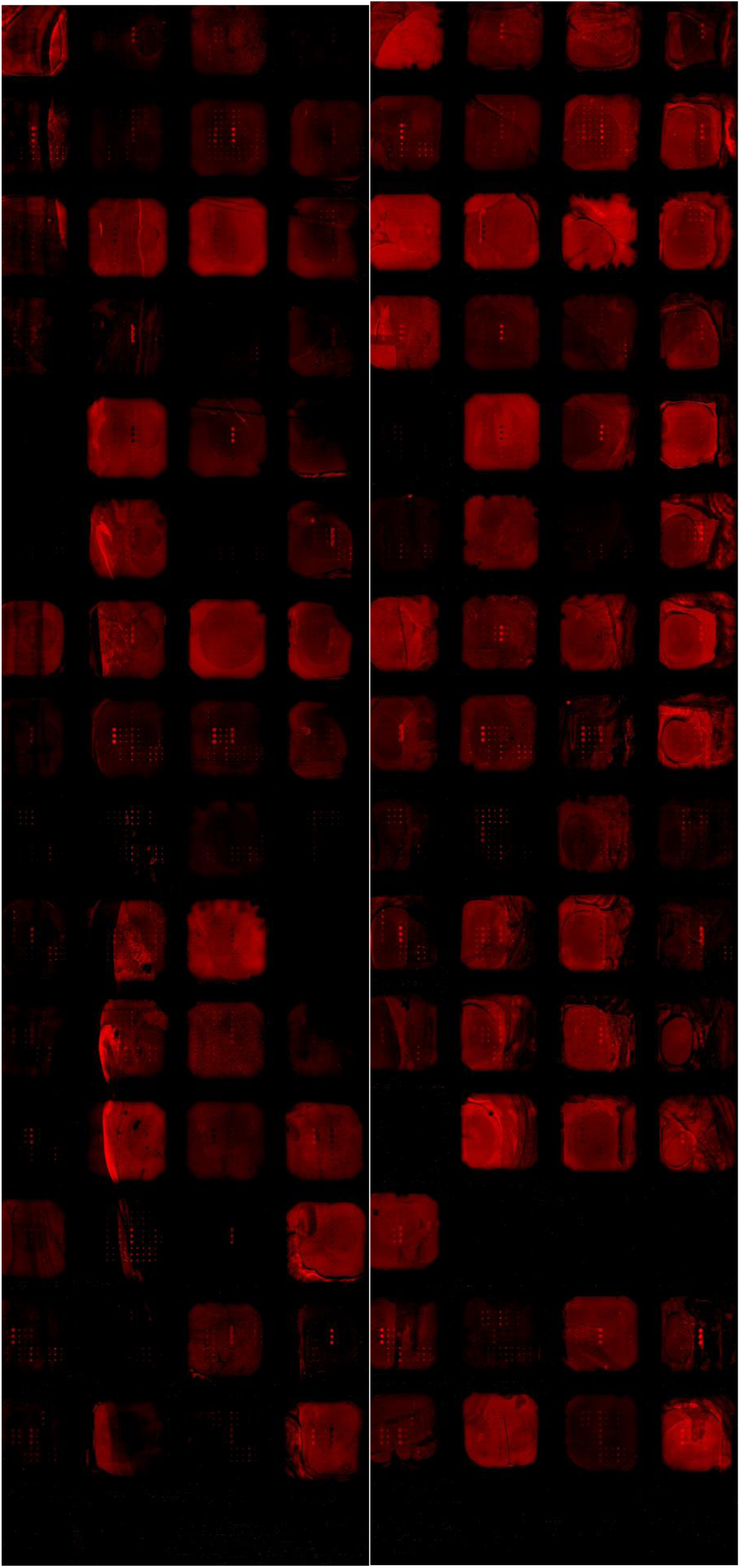

#### IgG human

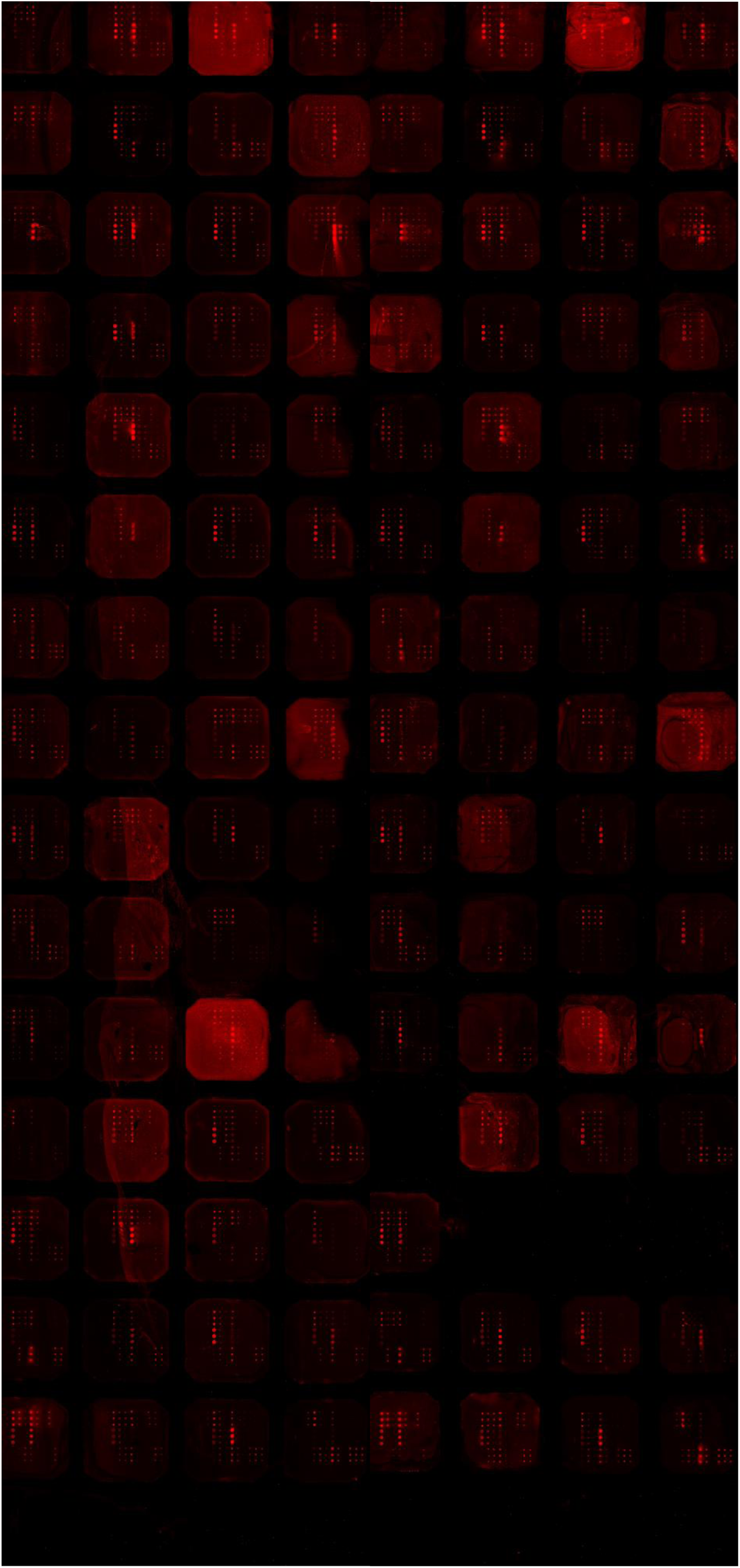

### Data Evaluation: Box-Plot and ROC curve for all data sets

Comparative Box-Plot of normalized fluorescence levels caused by antibody recognition by sera of different diseases stages, i.e. mouse naïve, d7, d14, d21 and d21 and human stage 1 and stage 2. Using GraphPadPrism 7.0, a significance test was performed using Tukey, followed by a two-way ANOVA and the calculation of receiver operation characteristics (ROC) curves. The area under the ROC curve was determined for the capability of the diagnostic tool to distinguish between A) non-infected and infected individuals and B) whether IgM and/or IgG are suitable antibody classes for this purpose.

#### IgM Mouse

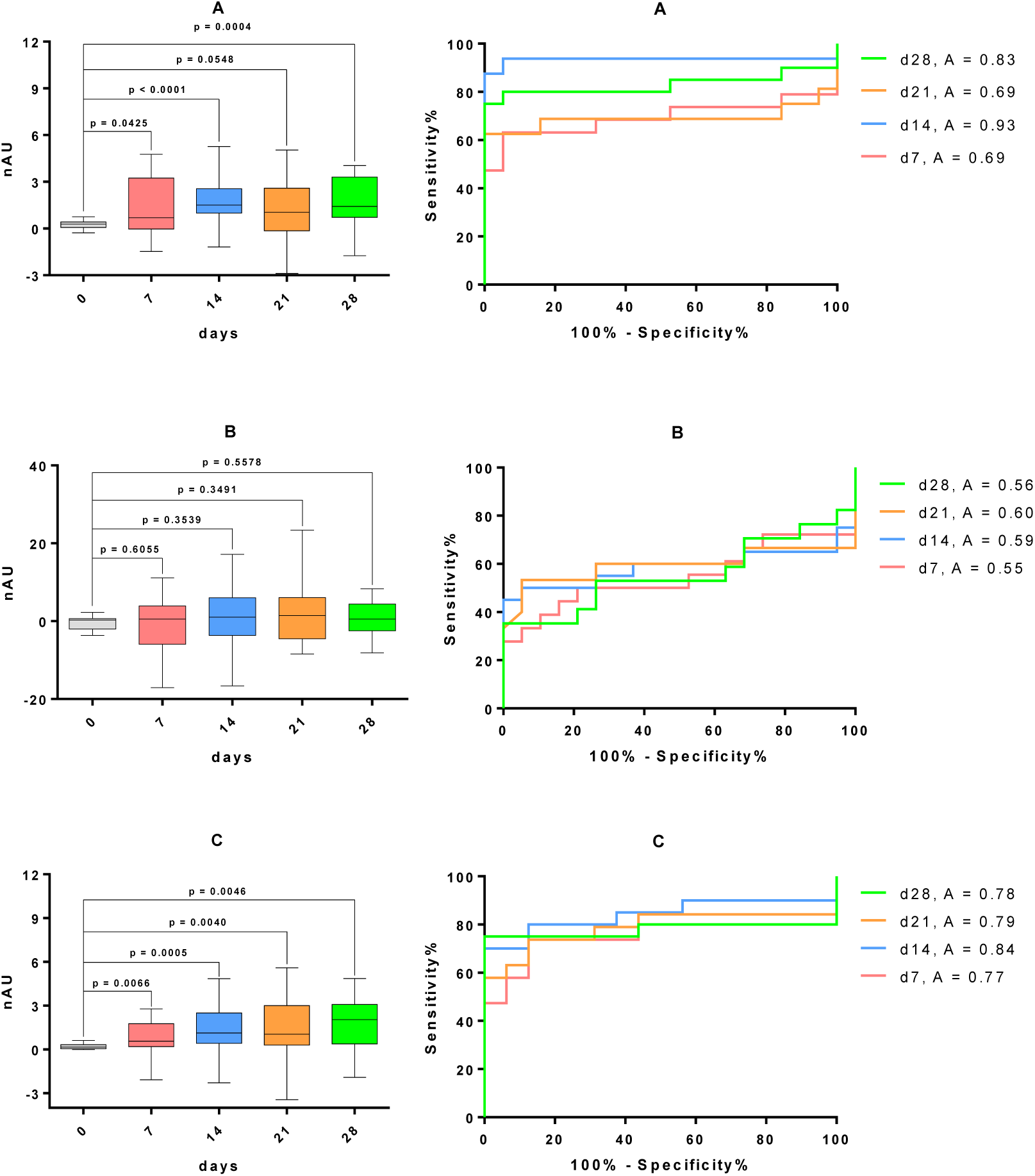
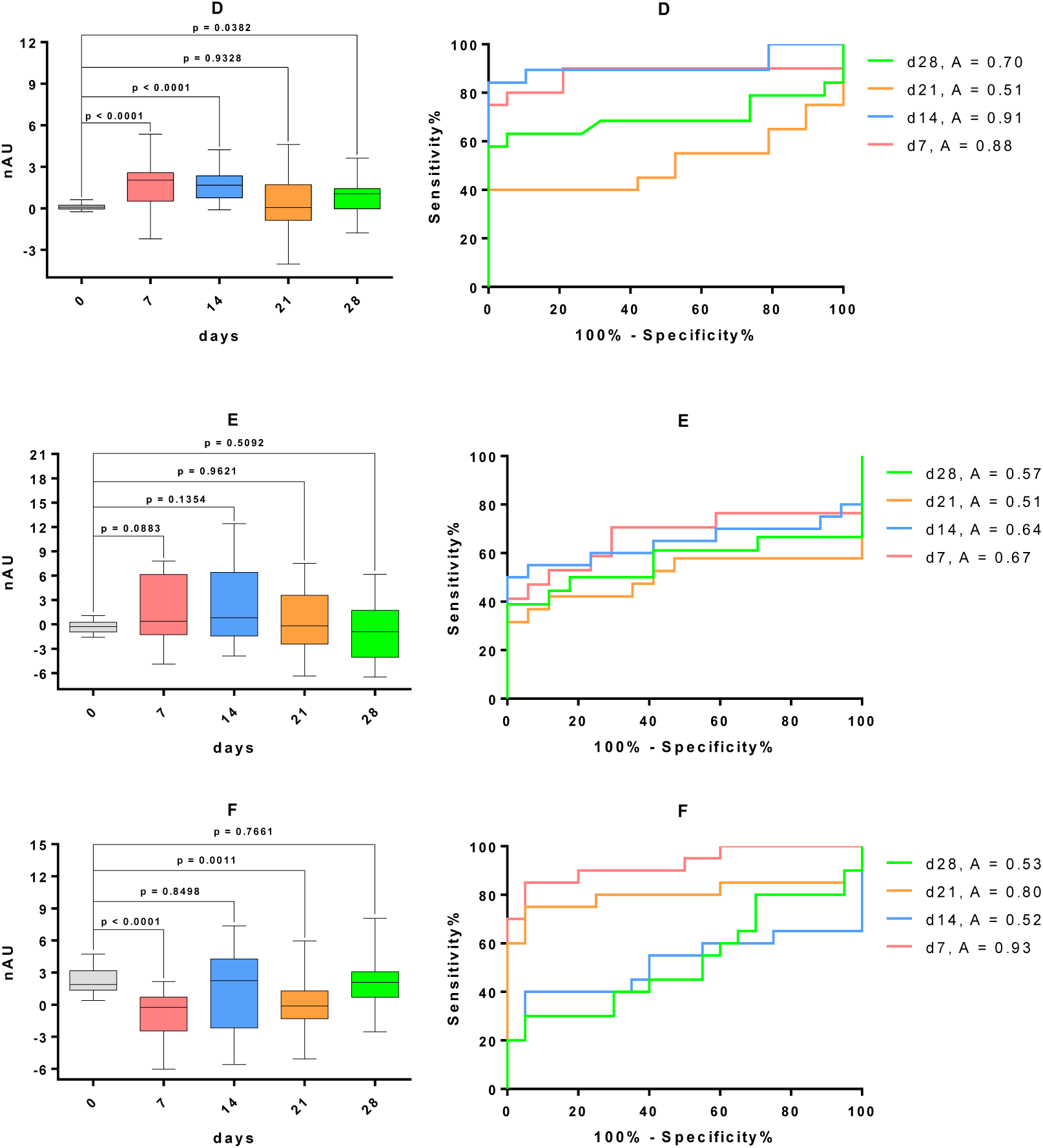

#### IgG Mouse

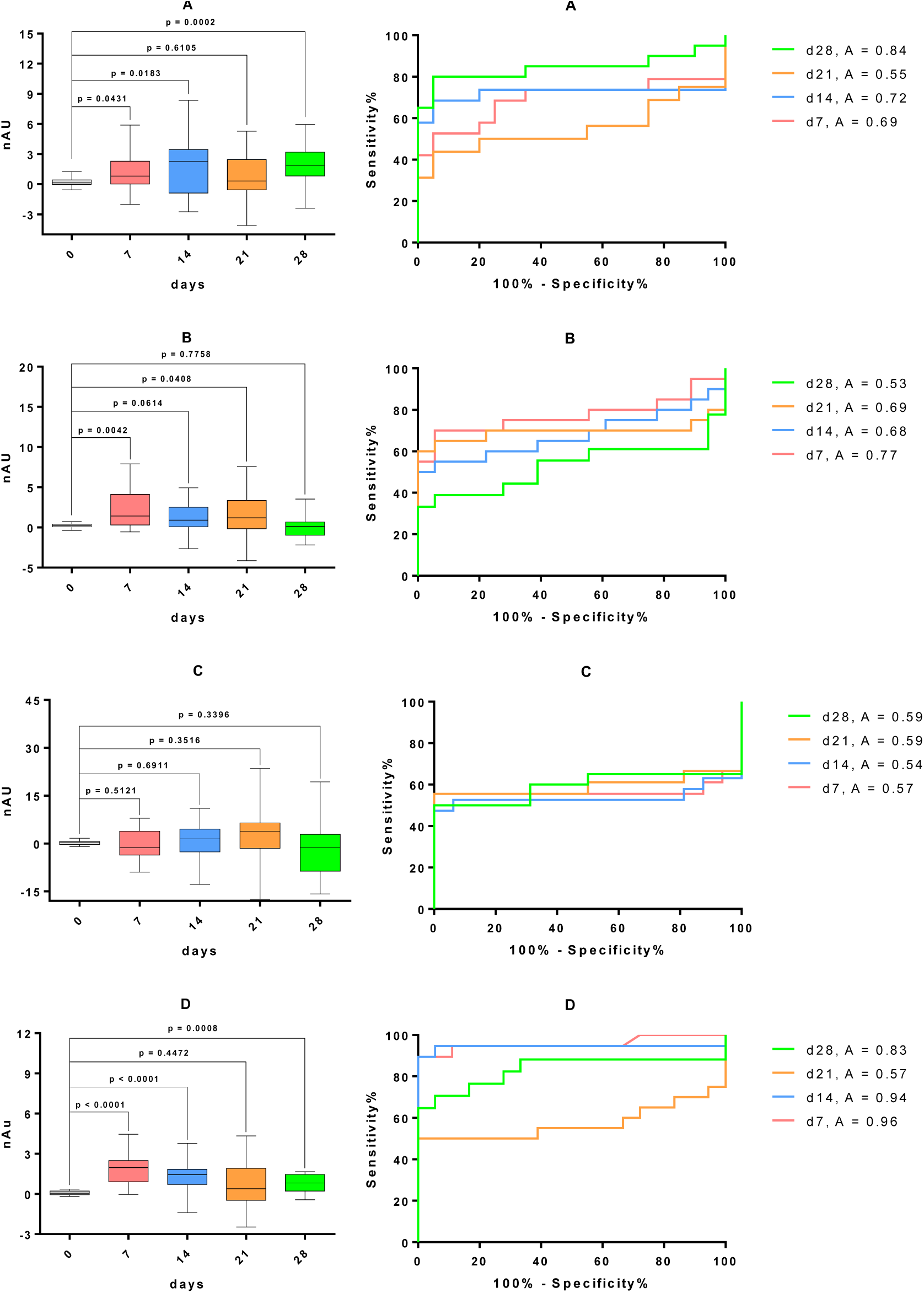
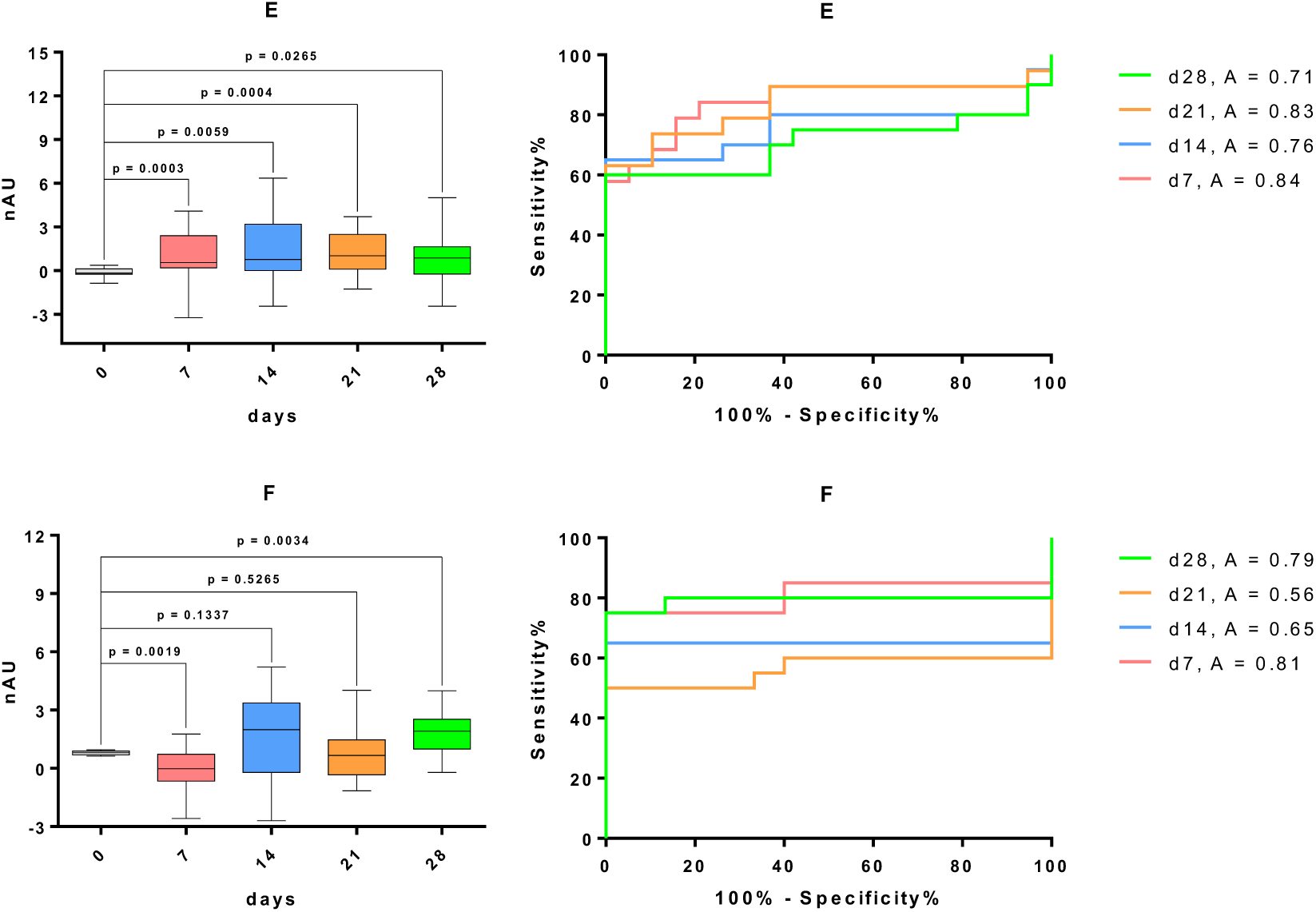

#### IgM human

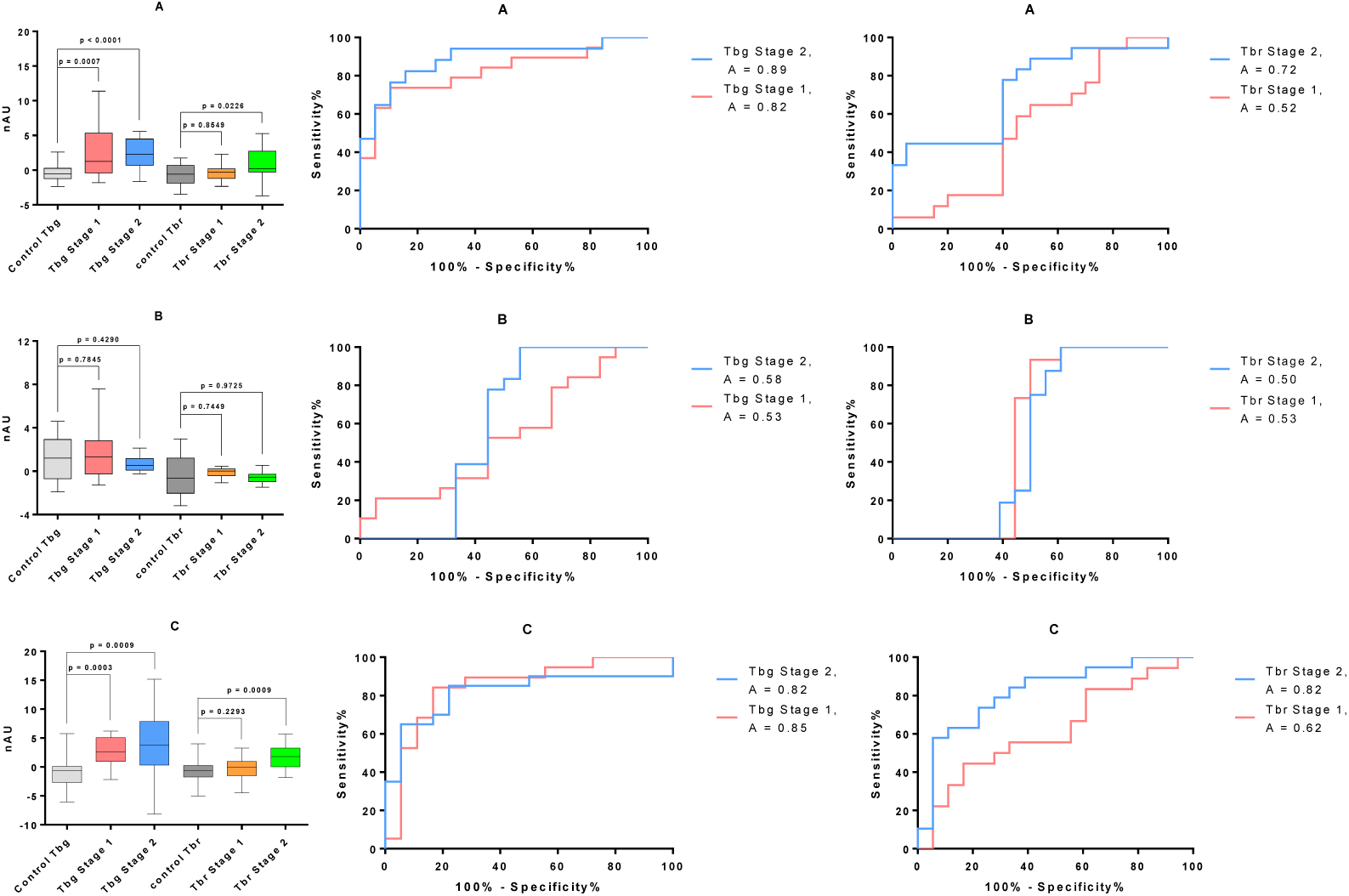
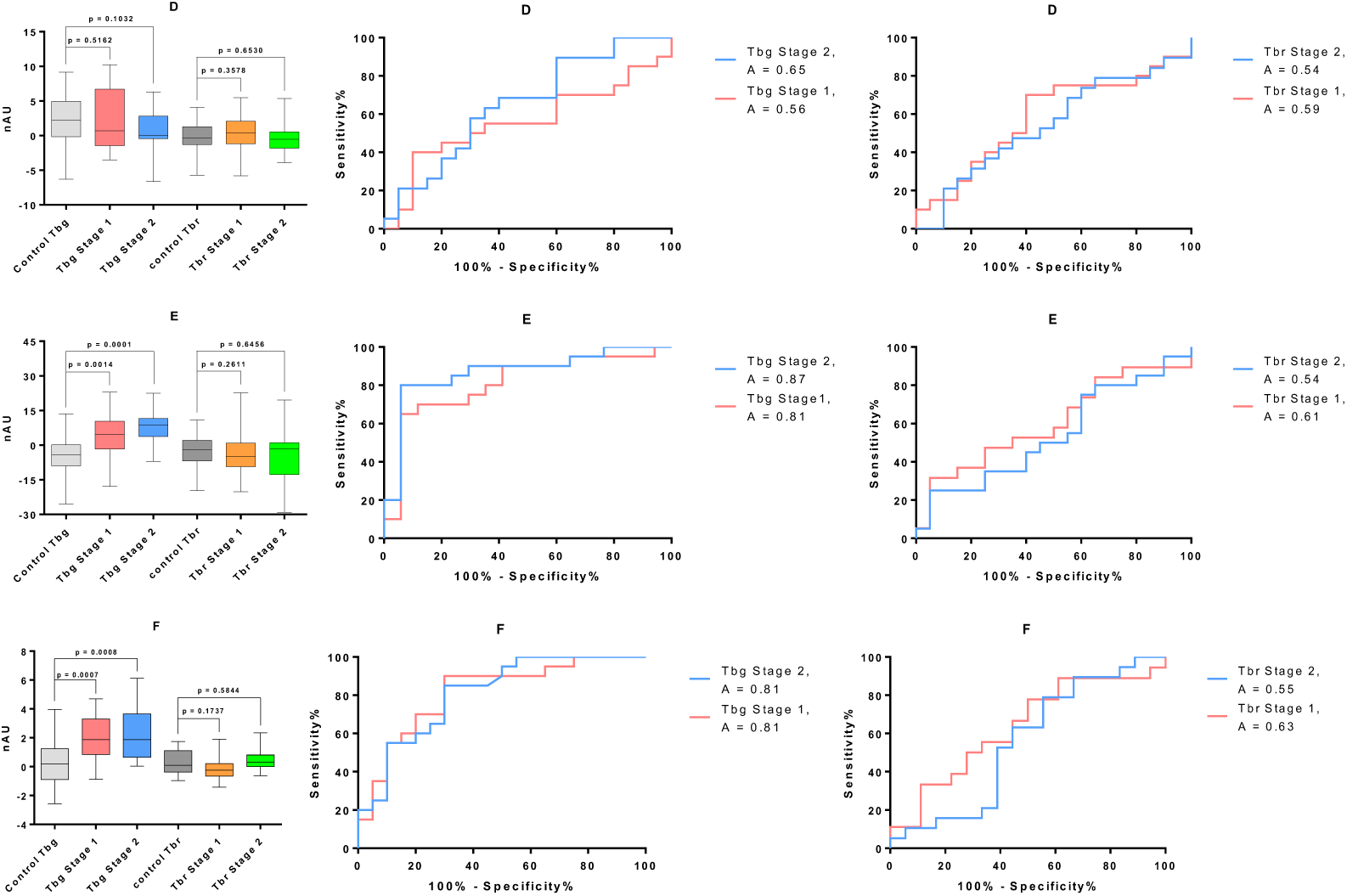

#### IgG human

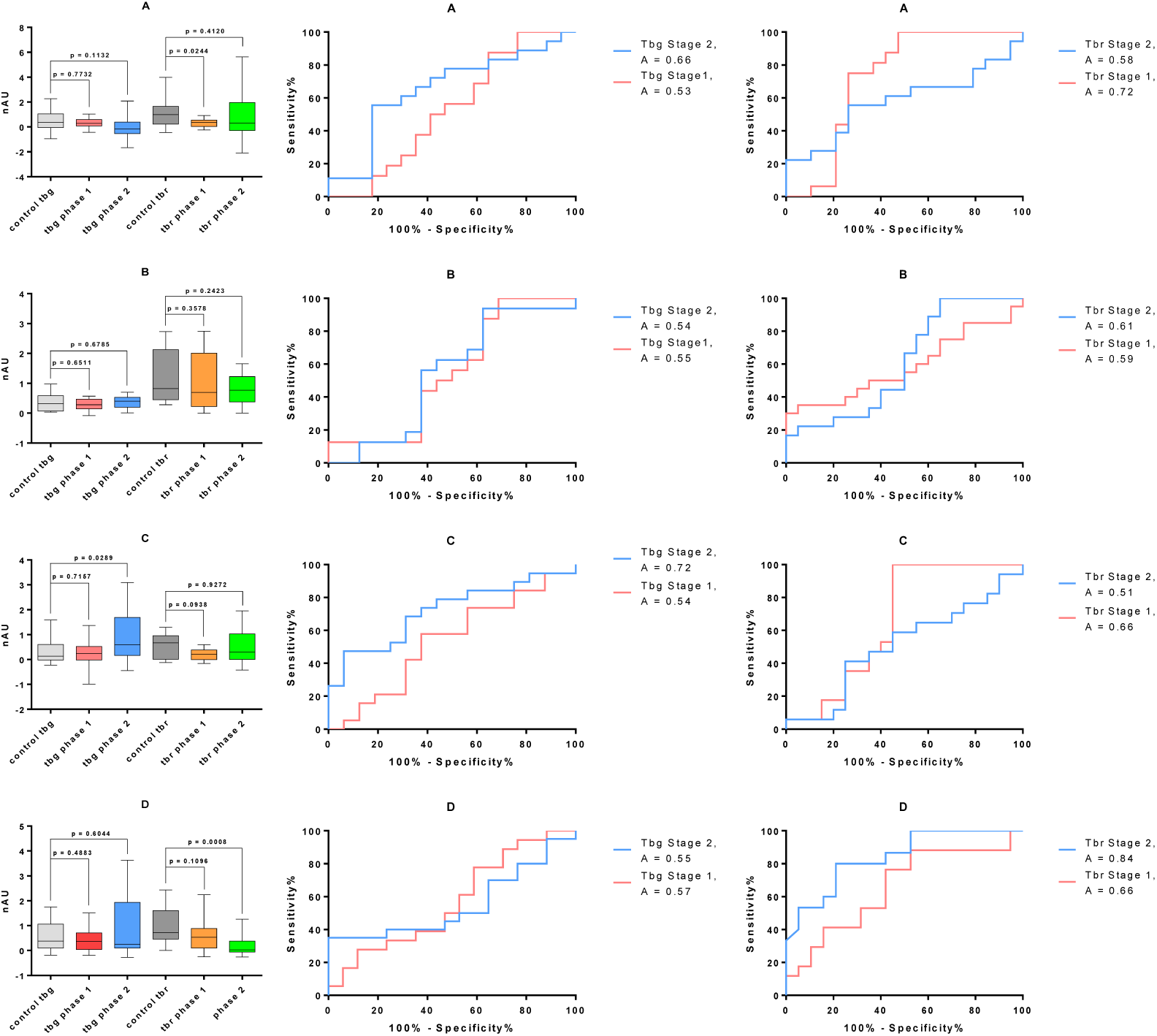
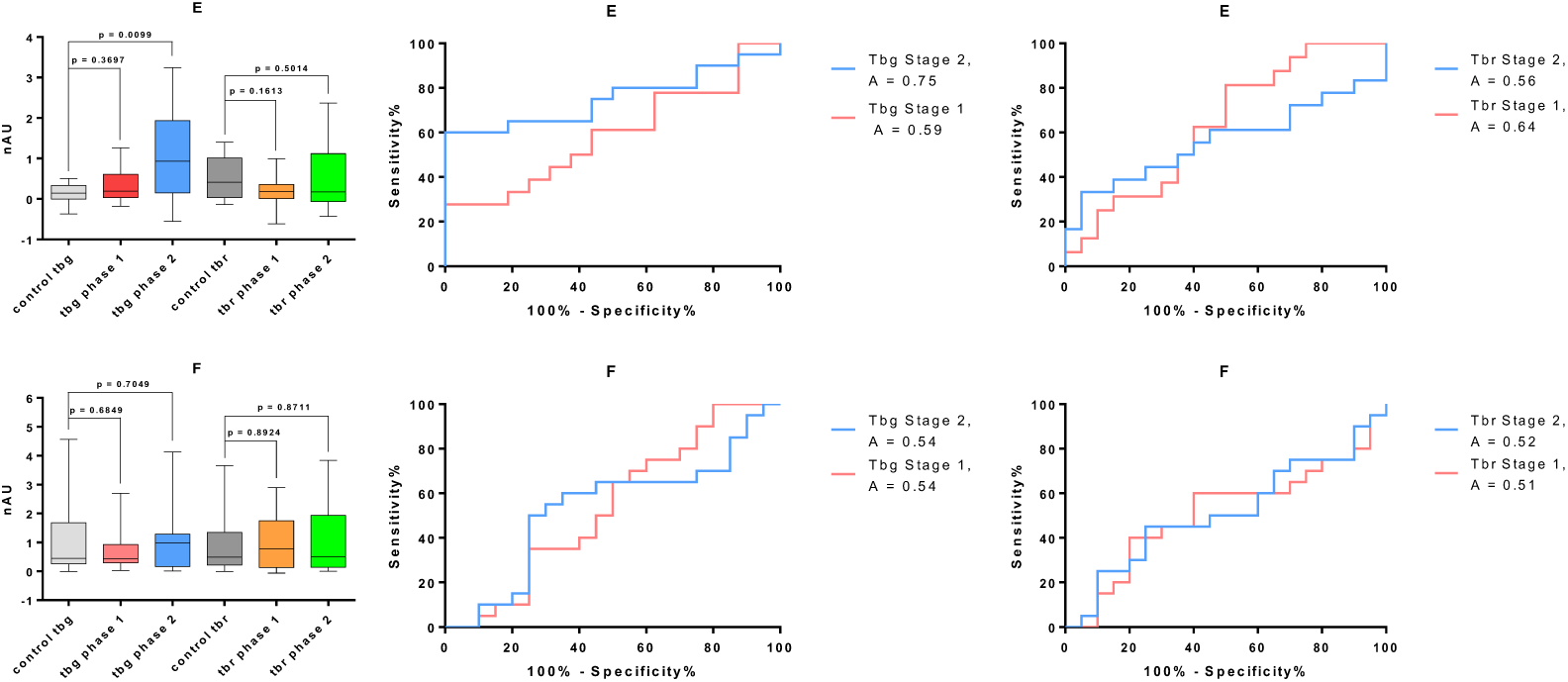

### Diagnostic power

Sensitivity, specificity and the respective 95% confidence intervals (CI) using the criteria of highest likelihood ratio for *T. brucei gambiense* and *T. brucei rhodesiense* infections are calculated as descriptors during generation of ROC curves. In a clinical setting, high likelihood ratios are preferred. This expectation allows for a moderate sensitivity as long as high specificity is maintained.

#### IgM mouse

| Structure/day | Sensitivity% | 95% CI | Specificity% | 95% CI | Likelihood ratio |
| --- | --- | --- | --- | --- | --- |
| Gal-Man d7 | 63,16 | 38,36% to 83,71% | 94,74 | 73,97% to 99,87% | 12,00 |
| Gal-Man d14 | 93,75 | 69,77% to 99,84% | 94,74 | 73,97% to 99,87% | 17,81 |
| Gal-Man d21 | 62,50 | 35,43% to 84,80% | 94,74 | 73,97% to 99,87% | 11,88 |
| Gal-Man d28 | 80,00 | 56,34% to 94,27% | 94,74 | 73,97% to 99,87% | 15,20 |
| Gal-Gal-Man d7 | 33,33 | 13,34% to 59,01% | 94,74 | 73,97% to 99,87% | 6,33 |
| Gal-Gal-Man d14 | 50,00 | 27,20% to 72,80% | 94,74 | 73,97% to 99,87% | 9,50 |
| Gal-Gal-Man d21 | 53,33 | 26,59% to 78,73% | 94,74 | 73,97% to 99,87% | 10,13 |
| Gal-Gal-Man d28 | 35,29 | 14,21% to 61,67% | 94,74 | 73,97% to 99,87% | 6,71 |
| Gal-Gal-(Gal)-Gal d7 | 57,89 | 33,50% to 79,75% | 93,75 | 69,77% to 99,84% | 9,26 |
| Gal-Gal-(Gal)-Gal d14 | 70,00 | 45,72% to 88,11% | 93,75 | 69,77% to 99,84% | 11,20 |
| Gal-Gal-(Gal)-Gal d21 | 63,16 | 38,36% to 83,71% | 93,75 | 69,77% to 99,84% | 10,11 |
| Gal-Gal-(Gal)-Gal d28 | 75,00 | 50,90% to 91,34% | 93,75 | 69,77% to 99,84% | 12,00 |
| P-Man-Man-Man d7 | 80,00 | 56,34% to 94,27% | 94,74 | 73,97% to 99,87% | 15,20 |
| P-Man-Man-Man d14 | 84,21 | 60,42% to 96,62% | 94,74 | 73,97% to 99,87% | 16,00 |
| P-Man-Man-Man d21 | 40,00 | 19,12% to 63,95% | 94,74 | 73,97% to 99,87% | 7,60 |
| P-Man-Man-Man d28 | 63,16 | 38,36% to 83,71% | 94,74 | 73,97% to 99,87% | 12,00 |
| VSG117ctdp d7 | 85,00 | 62,11% to 96,79% | 95,00 | 75,13% to 99,87% | 17,00 |
| VSG117ctdp d14 | 40,00 | 19,12% to 63,95% | 95,00 | 75,13% to 99,87% | 8,00 |
| VSG117ctdp d21 | 75,00 | 50,90% to 91,34% | 95,00 | 75,13% to 99,87% | 15,00 |
| VSG117ctdp 28 | 30,00 | 11,89% to 54,28% | 95,00 | 75,13% to 99,87% | 6,00 |
| Gal-(p)-Man-(Gal)-Man d7 | 47,06 | 22,98% to 72,19% | 94,12 | 71,31% to 99,85% | 8,00 |
| Gal-(p)-Man-(Gal)-Man d14 | 55,00 | 31,53% to 76,94% | 94,12 | 71,31% to 99,85% | 9,35 |
| Gal-(p)-Man-(Gal)-Man d21 | 36,84 | 16,29% to 61,64% | 94,12 | 71,31% to 99,85% | 6,26 |
| Gal-(p)-Man-(Gal)-Man d28 | 38,89 | 17,30% to 64,25% | 94,12 | 71,31% to 99,85% | 6,61 |

#### IgG Mouse

| Structure/day | Sensitivity% | 95% CI | Specificity% | 95% CI | Likelihood ratio |
| --- | --- | --- | --- | --- | --- |
| Gal-Man d7 | 52,63 | 28,86% to 75,55% | 95,00 | 75,13% to 99,87% | 10,53 |
| Gal-Man d14 | 68,42 | 43,45% to 87,42% | 95,00 | 75,13% to 99,87% | 13,68 |
| Gal-Man d21 | 43,75 | 19,75% to 70,12% | 95,00 | 75,13% to 99,87% | 8,75 |
| Gal-Man d28 | 80,00 | 56,34% to 94,27% | 95,00 | 75,13% to 99,87% | 16,00 |
| Gal-Gal-Man d7 | 70,00 | 45,72% to 88,11% | 94,44 | 72,71% to 99,86% | 12,60 |
| Gal-Gal-Man d14 | 55,00 | 31,53% to 76,94% | 94,44 | 72,71% to 99,86% | 9,90 |
| Gal-Gal-Man d21 | 65,00 | 40,78% to 84,61% | 94,44 | 72,71% to 99,86% | 11,70 |
| Gal-Gal-Man d28 | 38,89 | 17,30% to 64,25% | 94,44 | 72,71% to 99,86% | 7,00 |
| Gal-Gal-(Gal)-Gal d7 | 55,56 | 30,76% to 78,47% | 93,75 | 69,77% to 99,84% | 8,89 |
| Gal-Gal-(Gal)-Gal d14 | 52,63 | 28,86% to 75,55% | 93,75 | 69,77% to 99,84% | 8,42 |
| Gal-Gal-(Gal)-Gal d21 | 55,56 | 30,76% to 78,47% | 93,75 | 69,77% to 99,84% | 8,89 |
| Gal-Gal-(Gal)-Gal d28 | 50,00 | 27,20% to 72,80% | 93,75 | 69,77% to 99,84% | 8,00 |
| P-Man-Man-Man d7 | 89,47 | 66,86% to 98,70% | 94,44 | 72,71% to 99,86% | 16,11 |
| P-Man-Man-Man d14 | 94,74 | 73,97% to 99,87% | 94,44 | 72,71% to 99,86% | 17,05 |
| P-Man-Man-Man d21 | 50,00 | 27,20% to 72,80% | 94,44 | 72,71% to 99,86% | 9,00 |
| P-Man-Man-Man d28 | 70,59 | 44,04% to 89,69% | 94,44 | 72,71% to 99,86% | 12,71 |
| VSG117ctdp d7 | 75,00 | 50,90% to 91,34% | 93,33 | 68,05% to 99,83% | 11,25 |
| VSG117ctdp d14 | 65,00 | 40,78% to 84,61% | 93,33 | 68,05% to 99,83% | 9,75 |
| VSG117ctdp d21 | 50,00 | 27,20% to 72,80% | 93,33 | 68,05% to 99,83% | 7,50 |
| VSG117ctdp 28 | 75,00 | 50,90% to 91,34% | 93,33 | 68,05% to 99,83% | 11,25 |
| Gal-(p)-Man-(Gal)-Man d7 | 63,16 | 38,36% to 83,71% | 94,74 | 73,97% to 99,87% | 12,00 |
| Gal-(p)-Man-(Gal)-Man d14 | 65,00 | 40,78% to 84,61% | 94,74 | 73,97% to 99,87% | 12,35 |
| Gal-(p)-Man-(Gal)-Man d21 | 63,16 | 38,36% to 83,71% | 94,74 | 73,97% to 99,87% | 12,00 |
| Gal-(p)-Man-(Gal)-Man d28 | 60,00 | 36,05% to 80,88% | 94,74 | 73,97% to 99,87% | 11,40 |

#### IgM human

| Structure/Disease Stage | Sensitivity% | 95% CI | Specificity% | 95% CI | Likelihood ratio |
| --- | --- | --- | --- | --- | --- |
| Tb gambiense |  |  |  |  |  |
| Gal-Man Stage 1 | 63,16 | 38,36% to 83,71% | 94,74 | 73,97% to 99,87% | 12,00 |
| Gal-Man Stage 2 | 64,71 | 38,33% to 85,79% | 94,74 | 73,97% to 99,87% | 12,29 |
| Gal-Gal-Man Stage 1 | 21,05 | 6,052% to 45,57% | 94,44 | 72,71% to 99,86% | 3,79 |
| Gal-Gal-Man Stage 2 | 77,78 | 52,36% to 93,59% | 55,56 | 30,76% to 78,47% | 1,75 |
| Gal-Gal-(Gal)-Gal Stage 1 | 52,63 | 28,86% to 75,55% | 94,44 | 72,71% to 99,86% | 9,47 |
| Gal-Gal-(Gal)-Gal Stage 2 | 65,00 | 40,78% to 84,61% | 94,44 | 72,71% to 99,86% | 11,70 |
| p-Man-Man-Man Stage 1 | 40,00 | 19,12% to 63,95% | 90,00 | 68,30% to 98,77% | 4,00 |
| p-Man-Man-Man Stage 2 | 21,05 | 6,052% to 45,57% | 95,00 | 75,13% to 99,87% | 4,21 |
| Gal-(p)-Man-(Gal)-Man Stage 1 | 65,00 | 40,78% to 84,61% | 94,12 | 71,31% to 99,85% | 11,05 |
| Gal-(p)-Man-(Gal)-Man Stage 2 | 80,00 | 56,34% to 94,27% | 94,12 | 71,31% to 99,85% | 13,60 |
| VSG117ctdp Stage 1 | 35,00 | 15,39% to 59,22% | 95,00 | 75,13% to 99,87% | 7,00 |
| VSG117ctdp Stage 2 | 55,00 | 31,53% to 76,94% | 90,00 | 68,30% to 98,77% | 5,50 |
| Tb rhodesiense |  |  |  |  |  |
| Gal-Man Stage 1 | 52,94 | 27,81% to 77,02% | 55,00 | 31,53% to 76,94% | 1,18 |
| Gal-Man Stage 2 | 44,44 | 21,53% to 69,24% | 95,00 | 75,13% to 99,87% | 8,89 |
| Gal-Gal-Man Stage 1 | 93,33 | 68,05% to 99,83% | 50,00 | 26,02% to 73,98% | 1,87 |
| Gal-Gal-Man Stage 2 | 100,00 | 79,41% to 100,0% | 38,89 | 17,30% to 64,25% | 1,64 |
| Gal-Gal-(Gal)-Gal Stage 1 | 22,22 | 6,409% to 47,64% | 94,44 | 72,71% to 99,86% | 4,00 |
| Gal-Gal-(Gal)-Gal Stage 2 | 57,89 | 33,50% to 79,75% | 94,44 | 72,71% to 99,86% | 10,42 |
| p-Man-Man-Man Stage 1 | 70,00 | 45,72% to 88,11% | 60,00 | 36,05% to 80,88% | 1,75 |
| p-Man-Man-Man Stage 2 | 21,05 | 6,052% to 45,57% | 90,00 | 68,30% to 98,77% | 2,11 |
| Gal-(p)-Man-(Gal)-Man Stage 1 | 31,58 | 12,58% to 56,55% | 95,00 | 75,13% to 99,87% | 6,32 |
| Gal-(p)-Man-(Gal)-Man Stage 2 | 25,00 | 8,657% to 49,10% | 95,00 | 75,13% to 99,87% | 5,00 |
| VSG117ctdp Stage 1 | 33,33 | 13,34% to 59,01% | 88,89 | 65,29% to 98,62% | 3,00 |
| VSG117ctdp Stage 2 | 10,53 | 1,301% to 33,14% | 94,44 | 72,71% to 99,86% | 1,90 |

#### IgG human

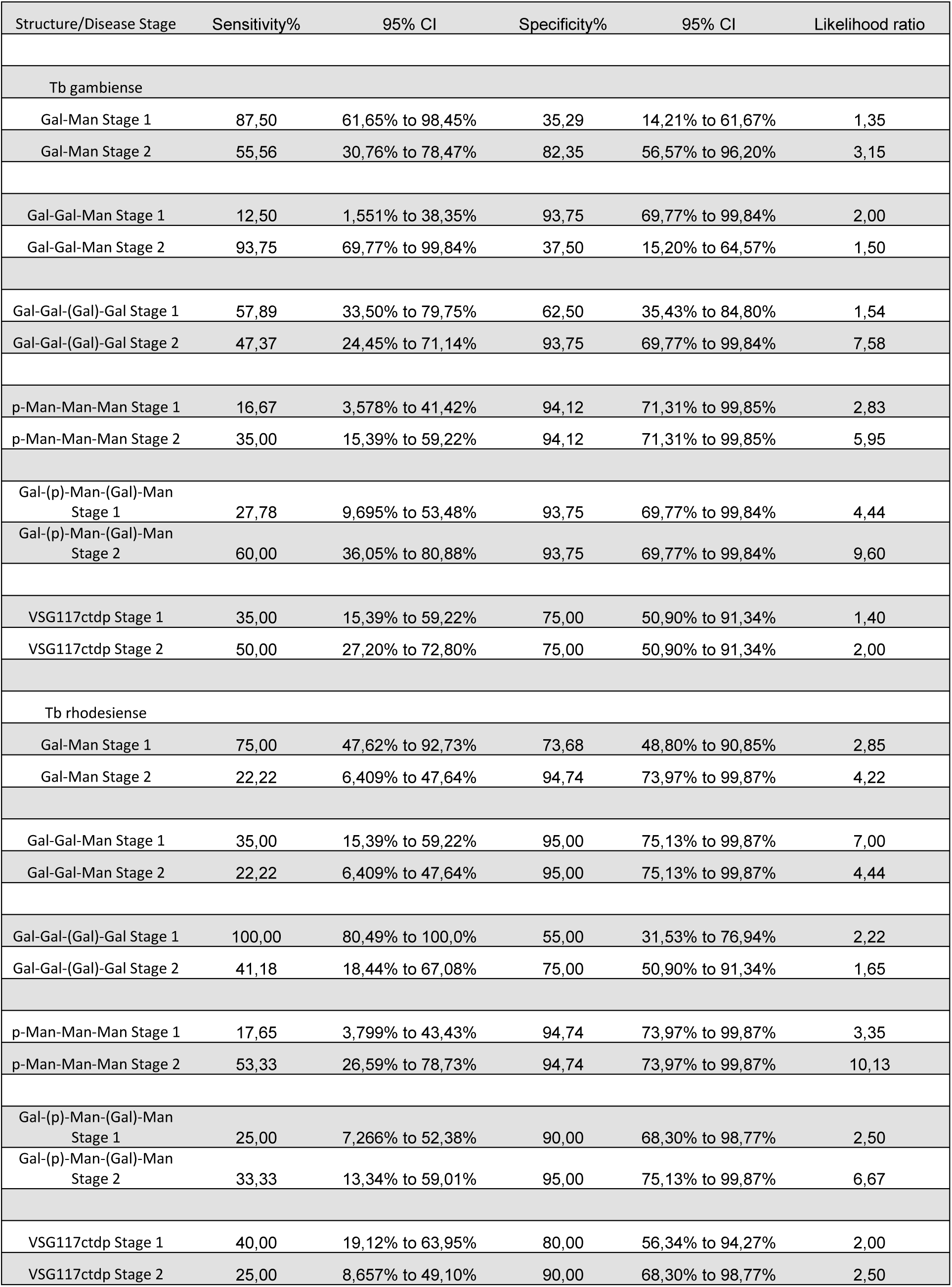

| Structure/Disease Stage | Sensitivity% | 95% CI | Specificity% | 95% CI | Likelihood ratio |
| --- | --- | --- | --- | --- | --- |
| Tb gambiense |  |  |  |  |  |
| Gal-Man Stage 1 | 87,50 | 61,65% to 98,45% | 35,29 | 14,21% to 61,67% | 1,35 |
| Gal-Man Stage 2 | 55,56 | 30,76% to 78,47% | 82,35 | 56,57% to 96,20% | 3,15 |
| Gal-Gal-Man Stage 1 | 12,50 | 1,551% to 38,35% | 93,75 | 69,77% to 99,84% | 2,00 |
| Gal-Gal-Man Stage 2 | 93,75 | 69,77% to 99,84% | 37,50 | 15,20% to 64,57% | 1,50 |
| Gal-Gal-(Gal)-Gal Stage 1 | 57,89 | 33,50% to 79,75% | 62,50 | 35,43% to 84,80% | 1,54 |
| Gal-Gal-(Gal)-Gal Stage 2 | 47,37 | 24,45% to 71,14% | 93,75 | 69,77% to 99,84% | 7,58 |
| p-Man-Man-Man Stage 1 | 16,67 | 3,578% to 41,42% | 94,12 | 71,31% to 99,85% | 2,83 |
| p-Man-Man-Man Stage 2 | 35,00 | 15,39% to 59,22% | 94,12 | 71,31% to 99,85% | 5,95 |
| Gal-(p)-Man-(Gal)-Man Stage 1 | 27,78 | 9,695% to 53,48% | 93,75 | 69,77% to 99,84% | 4,44 |
| Gal-(p)-Man-(Gal)-Man Stage 2 | 60,00 | 36,05% to 80,88% | 93,75 | 69,77% to 99,84% | 9,60 |
| VSG117ctdp Stage 1 | 35,00 | 15,39% to 59,22% | 75,00 | 50,90% to 91,34% | 1,40 |
| VSG117ctdp Stage 2 | 50,00 | 27,20% to 72,80% | 75,00 | 50,90% to 91,34% | 2,00 |
| Tb rhodesiense |  |  |  |  |  |
| Gal-Man Stage 1 | 75,00 | 47,62% to 92,73% | 73,68 | 48,80% to 90,85% | 2,85 |
| Gal-Man Stage 2 | 22,22 | 6,409% to 47,64% | 94,74 | 73,97% to 99,87% | 4,22 |
| Gal-Gal-Man Stage 1 | 35,00 | 15,39% to 59,22% | 95,00 | 75,13% to 99,87% | 7,00 |
| Gal-Gal-Man Stage 2 | 22,22 | 6,409% to 47,64% | 95,00 | 75,13% to 99,87% | 4,44 |
| Gal-Gal-(Gal)-Gal Stage 1 | 100,00 | 80,49% to 100,0% | 55,00 | 31,53% to 76,94% | 2,22 |
| Gal-Gal-(Gal)-Gal Stage 2 | 41,18 | 18,44% to 67,08% | 75,00 | 50,90% to 91,34% | 1,65 |
| p-Man-Man-Man Stage 1 | 17,65 | 3,799% to 43,43% | 94,74 | 73,97% to 99,87% | 3,35 |
| p-Man-Man-Man Stage 2 | 53,33 | 26,59% to 78,73% | 94,74 | 73,97% to 99,87% | 10,13 |
| Gal-(p)-Man-(Gal)-Man Stage 1 | 25,00 | 7,266% to 52,38% | 90,00 | 68,30% to 98,77% | 2,50 |
| Gal-(p)-Man-(Gal)-Man Stage 2 | 33,33 | 13,34% to 59,01% | 95,00 | 75,13% to 99,87% | 6,67 |
| VSG117ctdp Stage 1 | 40,00 | 19,12% to 63,95% | 80,00 | 56,34% to 94,27% | 2,00 |
| VSG117ctdp Stage 2 | 25,00 | 8,657% to 49,10% | 90,00 | 68,30% to 98,77% | 2,50 |

### Synthetic structures

*6-(benzylthio)hexyl 2,3,4-O-tri-benzyl-6-O-chloroacetyl-α-D-galactopyranosyl-(1→3)-2-O-acetyl-4-O-benzyl-6-O-tert-butyldiphenylsilyl-α-D-mannopyranoside* **(4)**

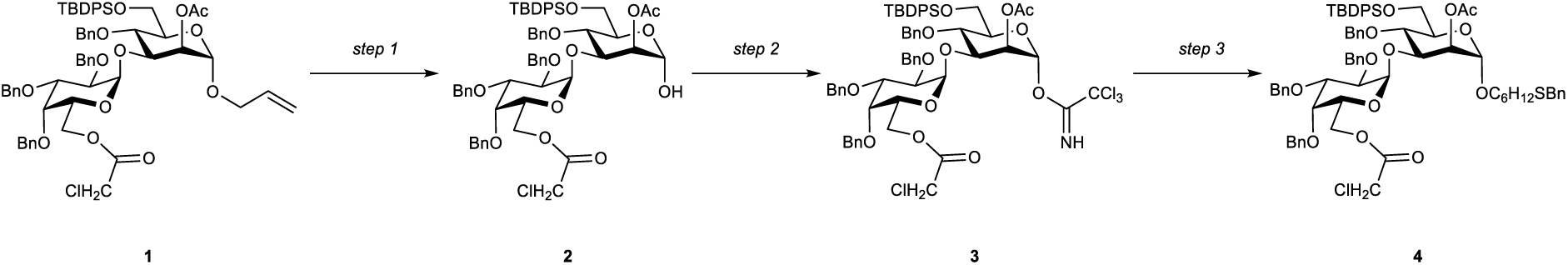

#### Step 1

10 mg of [IrCOD(PMePh_2_)_2_]PF_6_ were added to 2 ml THF. Hydrogen was bubbled through the suspension until the catalyst dissolved. The solution was transferred to a second flask, where it dissolved disaccharide **1**[1] (0.050 mmol, 0.055 g). The reaction was stirred at room temperature overnight. THF was evaporated under reduced pressure and the residue was dissolved in an 8:1 mixture of acetone and water. Mercury oxide (0.005 mmol, 1.0 mg) and mercury chloride (0.250 mmol, 67.8 mg) were added and the solution was stirred for one hour at room temperature. The reaction was quenched by adding saturated NaHCO_3_ -solution and the resulting mixture was extracted with DCM. The combined organic phases were dried over Na_2_SO_4_, filtered and concentrated under reduced pressure. The residue was purified by silica column chromatography using hexane and ethyl acetate as eluent. Product **2** was obtained in 65% yield (0.032 mmol, 0.034 g) as colorless oil. **R_f_** = 0.1 (4:1, Hex/AcOEt); **^1^H-NMR** (400 MHz, CDCl_3_): δ = 7.66 – 7.62 (m, 2H, H_Ar._), 7.55 (dt, *J* = 6.8 Hz, 1.4 Hz, 2H, H_Ar._), 7.35 – 7.17 (m, 16H, H_Ar._), 7.16 – 6.98 (m, 10H, H_Ar._), 5.25 (d, *J* = 3.6 Hz, 1H, Gal-1), 5.13 – 5.09 (m, 2H, Man-1, Man-2), 5.00 (d, *J* = 11.6 Hz, 1H, -CH_2_-), 4.86 (d, *J* = 11.6 Hz, 1H, -CH_2_-), 4.77 – 4.67 (m, 2H, -CH_2_-), 4.62 (d, *J* = 12.0 Hz, 1H, -CH_2_-), 4.57 – 4.48 (m, 3H, -CH_2_-), 4.27 – 4.17 (m, 2H), 4.10 – 4.04 (m, 2H), 4.01 – 3.96 (m, 4H), 3.93 – 3.78 (m, 5H), 3.73 (dd, *J* = 11.4, 1.7 Hz, 1H), 2.02 (s, 3H, -CH_3_), 1.01 (s, 9H, -C(-CH_3_)_3_) ppm; **^13^C-NMR** (101 MHz, CDCl_3_): δ = 170.8 (C=O), 167.3 (C=O), 138.8 (C_Ar._), 138.6 (C_Ar._), 138.4 (C_Ar._), 138.1 (C_Ar._), 136.1 (C_Ar._), 135.6 (C_Ar._), 134.0 (C_Ar._), 133.2 (C_Ar._), 129.8 (C_Ar._), 129.7 (C_Ar._), 128.6 (C_Ar._), 128.6 (C_Ar._), 128.3 (C_Ar._), 128.3 (C_Ar._), 128.0 (C_Ar._), 127.8 (C_Ar._), 127.7 (C_Ar._), 127.7 (C_Ar._), 127.6 (C_Ar._), 127.6 (C_Ar._), 127.5 (C_Ar._), 127.3 (C_Ar._), 126.9 (C_Ar._), 126.7 (C_Ar._), 99.7 (Gal-1), 92.4 (Man-1), 78.8, 77.5, 77.4, 77.2, 76.8, 75.6, 75.0, 74.7, 74.5, 74.5, 74.3, 73.5, 73.1, 73.0, 72.6, 69.2, 65.6, 62.8, 40.9, 27.0, 21.2, 19.6 ppm; **ESI-MS**: m/z M_calcd_ for C_60_H_67_ClO_13_Si = 1058.4039; M_found_ = 1081.3901 [M+Na]^+^; **[α]_D_^20^** = 40.07 (c = 0.1 g/L in CHCl_3_); **FTIR**: ν = 2934.63, 1745.15, 1455.60, 1241.04, 1061.56 cm^-1^.

#### Step 2

Hemiacetal **2** (0.032 mmol, 0.034 g) was dissolved in DCM at 0°C. Trichloroacetonitrile (0.256 mmol, 25.7 µl) and DBU (0.003 mmol, 0.5 µl) were added and the reaction was stirred until TLC indicated full conversion. The resulting mixture was concentrated under reduced pressure and was purified by silica column chromatography using hexane and ethyl acetate as eluent. Product **3** was obtained in 84% yield (0.027 mmol, 0.033 g) as colorless oil and was used directly for the next step. **R_f_** = 0.45 (4:1, Hex/AcOEt); **^1^H-NMR** (400 MHz, CDCl_3_): δ = 8.62 (s, 1H, =NH), 7.63 (dt, *J* = 6.9 Hz, 1.5 Hz, 2H, H_Ar._), 7.58 – 7.53 (m, 2H, H_Ar._), 7.37 – 7.01 (m, 26H), 6.29 (d, *J* = 2.0 Hz, 1H, Man-1), 5.24 (d, *J* = 2.3 Hz, 1H, Gal-1), 5.13 – 5.07 (m, 2H, Man-2, -CH_2_-), 4.89 (d, *J* = 11.4 Hz, 1H, -CH_2_-), 4.82 – 4.63 (m, 4H, -CH_2_-), 4.58 – 4.50 (m, 3H), 4.19 – 4.16 (m, 2H), 4.10 – 3.75 (m, 11H), 2.05 (s, 3H, -CH_3_), 1.00 (s, 9H, -C(-CH_3_)_3_) ppm.

#### Step 3

Imidate **3** (0.027 mmol, 0.033 g) and 6-Thiobenzylhexanol (0.012 g, 0.054 mmol) were coevaporated three times with toluene, dried under high vacuum and dissolved in DCM. Freshly activated powdered molecular sieves were added and the suspension was stirred for 10 minutes. TMSOTf (0.008 mmol, 1.5 µl) was added at 0°C. After TLC indicated full conversion, the reaction was quenched by adding saturated NaHCO_3_-solution and extracted with DCM. The combined organic phases were dried over MgSO_4_, filtered and evaporated. The residue was purified by silica gel chromatography giving product **5** quantitative yield (0.027 mmol, 0.034 g) as colorless oil. **R_f_** = 0.45 (4:1, Hex/AcOEt); **^1^H-NMR** (400 MHz, CDCl_3_): δ = 7.64 (dt, *J* = 6.7 Hz, 1.5 Hz, 2H, H_Ar._), 7.57 (dt, *J* = 6.8 Hz, 1.4 Hz, 2H, H_Ar._), 7.35 – 7.06 (m, 29H, H_Ar._), 7.00 – 6.96 (m, 2H, H_Ar._), 5.19 (d, *J* = 3.6 Hz, 1H, Gal-1), 5.04 (dd, *J* = 3.4 Hz, 1.7 Hz, 1H, Man-1), 4.98 (d, *J* = 11.4 Hz, 1H, -CH_2_-), 4.87 (d, *J* = 11.6 Hz, 1H, -CH_2_-), 4.77 – 4.68 (m, 4H, -CH_2_-, Man-1), 4.66 (d, *J* = 12.1 Hz, 1H, -CH_2_-), 4.55 – 4.50 (m, 2H, -CH_2_-), 4.47 (d, *J* = 11.5 Hz, 1H, -CH_2_-), 4.22 – 4.08 (m, 2H), 4.04 – 3.73 (m, 13H), 3.66 – 3.40 (m, 2H), 3.29 (dt, *J* = 9.7 Hz, 6.8 Hz, 1H), 2.37 – 2.27 (m, 4H, -CH_2_-), 2.00 (s, 3H, -CH_3_), 1.57 – 1.12 (m, 8H), 0.99 (s, 9H) ppm; **^13^C-NMR** (101 MHz, CDCl_3_): δ = 170.8 (C=O), 167.0 (C=O), 138.8 (C_Ar._), 138.6 (C_Ar._), 138.4 (C_Ar._), 138.2 (C_Ar._), 136.1 (C_Ar._), 135.7 (C_Ar._), 133.9 (C_Ar._), 133.2 (C_Ar._), 129.7 (C_Ar._), 129.7 (C_Ar._), 128.9 (C_Ar._), 128.6 (C_Ar._), 128.6 (C_Ar._), 128.5 (C_Ar._), 128.3 (C_Ar._), 128.3 (C_Ar._), 128.3 (C_Ar._), 128.0 (C_Ar._), 127.8 (C_Ar._), 127.7 (C_Ar._), 127.7 (C_Ar._), 127.6 (C_Ar._), 127.6 (C_Ar._), 127.5 (C_Ar._), 127.3 (C_Ar._), 127.1 (C_Ar._), 127.0 (C_Ar._), 127.0 (C_Ar._), 100.0 (Gal-1), 97.1 (Man-1), 78.8, 77.5, 77.4, 77.2, 76.8, 76.6, 75.7, 74.8, 74.7, 74.5, 74.3, 73.5, 73.1, 72.8, 72.6, 69.1, 67.7, 65.4, 64.6, 62.9, 40.9, 36.4, 36.4, 31.4, 29.9, 29.4, 29.2, 28.8, 28.6, 28.6, 26.9, 25.8, 25.7, 21.2, 19.5 ppm; **ESI-MS**: m/z M_calcd_ for C_73_H_85_ClO_13_SSi: 1264.5169; M_found_ = 1287.5035 [M+Na]^+^; **[α]_D_^20^** = 65.75 (c = 0.1 g/L in CHCl_3_); **FTIR**: ν = 3032.53, 2932.25, 2859.17, 2211.79, 2162.52, 1743.16, 1496.93, 1454.99, 1429.28, 1364.17, 1239.58, 1135.16, 1101.34, 1059.38, 1028.84, 824.05, 798.05, 737.48, 700.11 cm^-1^.

*6-Mercaptohexyl α-D-galactopyranosyl-(1→3)-α-D-mannopyranoside **(A)***

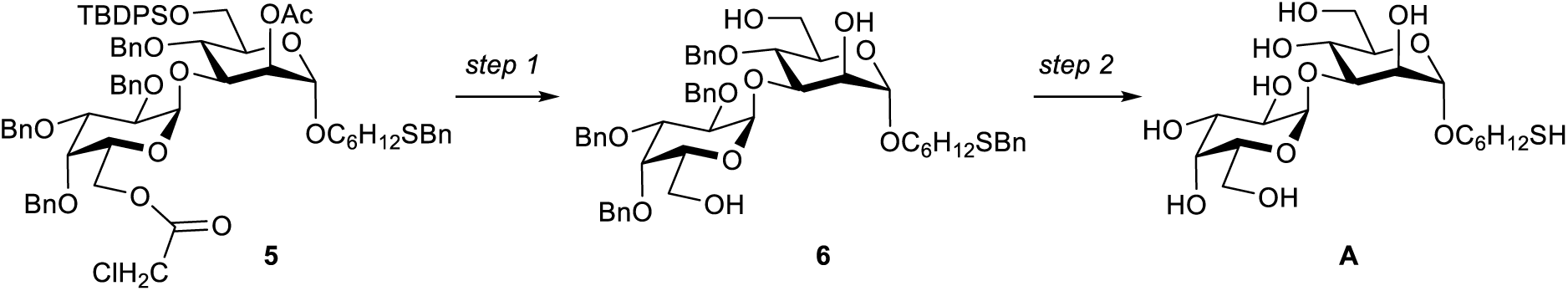

#### Step 1

Disaccharide **5** (0.027 mmol, 0.034 g) was dissolved in a mixture of DCM and Methanol. Acetyl chloride (0.1 ml) was added dropwise and the reaction mixture was stirred until TLC indicated full conversion. The green reaction was diluted with DCM, quenched with saturated NaHCO_3_-solution and extracted with DCM. The combined organic phases were dried over MgSO_4_, filtered and evaporated. The residue was purified by silica gel chromatography giving product **6** in 48% (0.013 mmol, 0.012 g) as colorless oil. **R_f_** = 0.2 (2:1, Hex/AcOEt); **^1^H-NMR** (400 MHz, CDCl_3_): δ = 7.34 – 7.07 (m, 25H, H_Ar._), 5.06 – 5.01 (m, 2H, -CH_2_-, Gal-1), 4.86 (d, *J* = 11.7 Hz, 1H, -CH_2_-), 4.77 – 4.58 (m, 5H, -CH_2_-, Man-1), 4.53 – 4.48 (m, 2H, -CH_2_-), 4.09 – 3.99 (m, 3H), 3.90 – 3.79 (m, 3H), 3.78 – 3.65 (m, 3H), 3.62 (s, 2H), 3.60 – 3.43 (m, 3H), 3.32 – 3.21 (m, 2H), 2.33 (t, *J* = 7.3 Hz, 2H), 1.52 – 1.41 (m, 4H), 1.31 -1.17 (m, 4H) ppm; **^13^C-NMR** (101 MHz, CDCl_3_): δ = 138.5 (C_Ar._), 138.4 (C_Ar._), 138.4 (C_Ar._), 138.2 (C_Ar._), 128.8 (C_Ar._), 128.5 (C_Ar._), 128.4 (C_Ar._), 128.4 (C_Ar._), 128.3 (C_Ar._), 128.3 (C_Ar._), 127.9 (C_Ar._), 127.8 (C_Ar._), 127.7 (C_Ar._), 127.6 (C_Ar._), 127.6 (C_Ar._), 127.5 (C_Ar._), 126.8 (C_Ar._), 100.0 (Gal-1), 98.7 (Man-1), 82.5, 78.5, 77.3, 77.2, 77.0, 76.9, 76.7, 75.3, 74.9, 74.3, 73.3, 73.2, 73.1, 71.7, 71.4, 69.7, 67.7, 63.1, 61.9, 36.2, 31.2, 29.2, 29.0, 28.6, 25.7 ppm; **ESI-MS**: m/z M_calcd_ for C_53_H_64_O_11_S = 908.4169; M_found_ = 931.4075 [M+Na]^+^; **[α]_D_^20^** = 30.76 (c = 0.1 g/L in CHCl_3_); **FTIR**: ν = 3393.73, 3032.10, 2930.62, 2163.30, 2036.82, 1497.01, 1454.56, 1352.26, 1096.52, 1068.04, 1043.31, 738.63, 698.46, 682.08, 660.41 cm^-1^.

#### Step 2

Ammonia (10 ml) was condensed in a flask at -78°C and two drops of methanol were added. Sodium was added in small pieces until a dark blue color was established. Triol **6** (0.013 mmol, 0.012 g) was dissolved in 1 ml THF and added to the ammonia solution. The reaction was stirred for 1 h, subsequently adding more sodium when the blue color disappeared. The reaction was quenched by adding methanol and ammonia was blown off using a stream of nitrogen. The pH of the resulting solution was adjusted to 7-8 with glacial acetic acid. The residue was concentrated under reduced pressure and purified by size exclusion using 5% ethanol in water as eluent and RP-HPLC (hypercarb column 150×10mm, ThermoFisher, 5 µ, acetonitrile in water 0-100% in 60 min). Product **A** was obtained in 50% yield (6.540 µmol, 0.3 mg) as white solid. **^1^H-NMR** (400 MHz, D_2_O): δ = 5.12 (d, *J* = 4.1 Hz, 1H, Gal-1), 4.72 (1H, Man-1), 4.01 – 3.21 (m, 16H), 1.51 – 1.42 (m, 4H), 1.24 – 1.05 (m, 4H) ppm; **^13^C-NMR** (151 MHz, D_2_O): δ = 100.0 (Gal-1), 99.4 (Man-1), 72.7, 71.2, 69.1, 68.6, 66.4, 66.0, 64.6, 30.7, 21.9 ppm; **ESI-MS**: m/z M_calcd_ for C_18_H_34_O_11_S = 458.1822; M_found_ = 937.3439 [2M+Na]^+^

*2,3,4-O-tri-benzyl-6-O-chloroacetyl-α-D-galactopyranosyl-(1→6)-2,3,4-O-tri-benzyl-α-D-galactopyranosyl-(1→3)-2-O-acetyl-4-O-benzyl-6-O-tert-butyldiphenylsilyl-α-D-mannopyranosyl trichloracetimidate* (**9**)

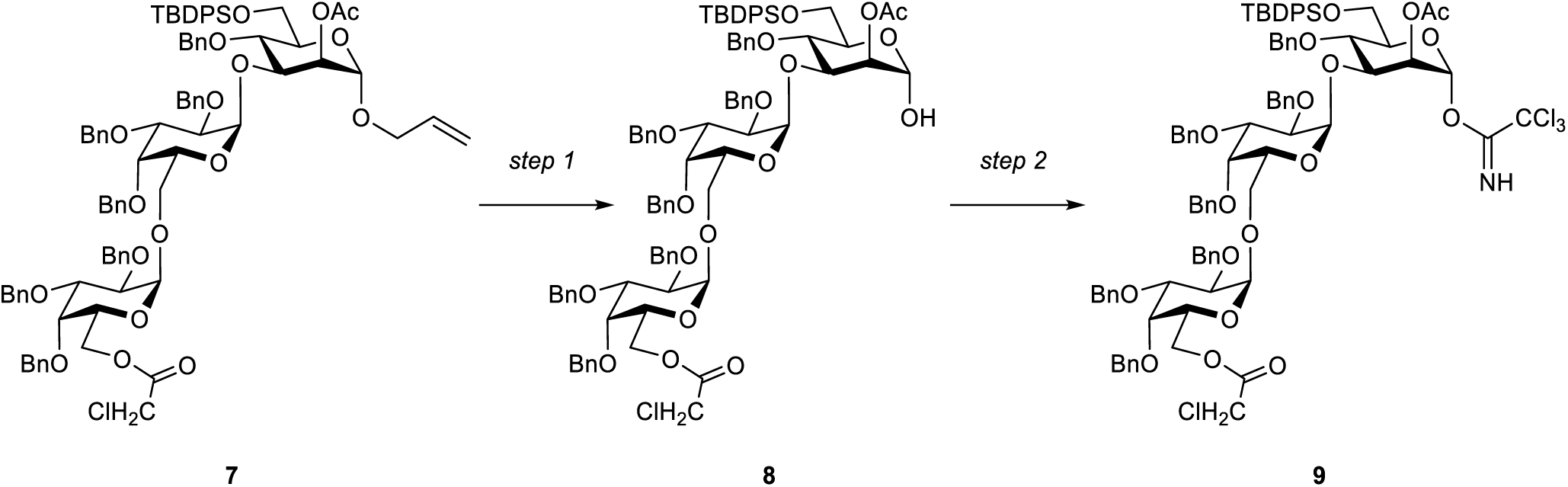

#### Step 1

Hydrogen was bubbled through a suspension of 10 mg of [IrCOD(PMePh_2_)_2_]PF_6_ in 2 ml THF until the catalyst was dissolved. The solution was transferred to a second flask containing trisaccharide **7**[1] (0.014 mmol, 0.022 g) and the resulting reaction mixture was stirred under hydrogen atmosphere at room temperature overnight. The THF was evaporated under reduced pressure and the residue was dissolved in an 8:1 mixture of acetone and water. Mercury oxide (0.001 mmol, 0.2 mg) and mercury chloride (0.070 mmol, 19.0 mg) were added and the mixture was stirred for one hour at room temperature. The reaction was quenched by adding saturated NaHCO_3_ -solution and the resulting mixture was extracted with DCM. The combined organic phases were dried over Na_2_SO_4_, filtered and concentrated under reduced pressure. The residue was purified by silica column chromatography using hexane and ethyl acetate as eluent. Product **8** was obtained in 45% yield (6.430 µmol, 9.6 mg) as colorless oil. **R_f_** = 0.1 (4:1, Hex/AcOEt); **^1^H-NMR** (400 MHz, CDCl_3_): δ = 7.64 (d, *J* = 7.3 Hz, 2H, H_Ar._), 7.55 (d, *J* = 7.3 Hz, 2H, H_Ar._), 7.39 – 6.99 (m, 41H, H_Ar._), 5.29 (s, 1H, Gal-1), 5.06 (s, 1H, Man-1), 4.95 (d, *J* = 11.8 Hz, 1H, Gal‘-1), 4.88 – 4.80 (m, 3H, Gal-1, Gal‘-6, -CH_2_-), 4.76 – 4.42 (m, 13H, -CH_2_-), 4.36 – 4.22 (m, 2H), 4.11 (s, 1H, Man-1), 4.06 – 3.61 (m, 16H), 3.08 (d, *J* = 10.1 Hz, Man-6), 2.00 (s, 3H, -CH_3_), 0.98 (s, 9H, -C(-CH_3_)_3_) ppm; **^13^C-NMR** (101 MHz, CDCl_3_): <colcnt=9> δ = 170.8 (C=O), 167.2 (C=O), 138.9 (C_Ar._), 138.7 (C_Ar._), 138.6 (C_Ar._), 138.5 (C_Ar._), 138.3 (C_Ar._), 138.2 (C_Ar._), 138.1 (C_Ar._), 137.5 (C_Ar._), 136.2 (C_Ar._), 135.7 (C_Ar._), 129.6 (C_Ar._), 128.6 (C_Ar._), 128.6 (C_Ar._), 128.5 (C_Ar._), 128.5 (C_Ar._), 128.4 (C_Ar._), 128.3 (C_Ar._), 128.2 (C_Ar._), 128.2 (C_Ar._), 128.1 (C_Ar._), 128.0 (C_Ar._), 127.8 (C_Ar._), 127.7 (C_Ar._), 127.7 (C_Ar._), 127.7 (C_Ar._), 127.6 (C_Ar._), 127.6 (C_Ar._), 127.4 (C_Ar._), 127.3 (C_Ar._), 126.9 (C_Ar._), 99.1 (Gal-1), 97.3 (Gal-1), 92.0 (Man-1), 79.0, 78.6, 77.5, 77.4, 77.2, 76.8, 75.7, 75.2, 74.9, 74.6, 74.5, 73.9, 73.4, 73.4, 73.3, 73.1, 72.8, 72.6, 70.0, 68.0, 64.9, 62.9, 40.8 (-CClH_2_), 26.9 (-*C*(-CH_3_)_3_), 21.4 (-C(-*C*H_3_)_3_), 19.6 (-CH_3_) ppm; **ESI-MS**: m/z M_calcd_ for C_87_H_95_ClO_18_Si = 1490.5976; M_found_ = 1513.5905 [M+Na]^+^; **[α]_D_^20^** = 54.35 (c = 0.1 g/L in CHCl_3_); **FTIR**: ν = 3453.56, 3033.13, 2930.22, 2172.92, 2128.27, 2037.40, 1966.26, 1738.57, 1497.85, 1455.58, 1429.05, 1361.71, 1241.71, 1103.51, 1059.90, 1028.48, 825.28, 739.79, 698.76, 664.27 cm^-1^.

#### Step 2

Hemiacetal **8** (6.430 μmol, 9.6 mg) was dissolved in DCM at 0°C. Trichloroacetonitrile (0.051 mmol, 5.1 µl) and DBU (0.643 μmol, 0.1 µl) were added and the reaction was stirred for until TLC indicated full conversion. The resulting mixture was concentrated under reduced pressure and was purified by silica column chromatography using hexane and ethyl acetate as eluent. Imidate **9** was obtained in 95% yield (6.110 µmol, 10.0 mg) as colorless oil and used for the next step. **R_f_** = 0.6 (4:1, Hex/AcOEt); **^1^H-NMR** (400 MHz, CDCl_3_): δ = 8.58 (s, 1H, =NH), 7.67 – 7.55 (m, 5H), 7.38 – 7.01 (m, 40H), 6.25 (s, 1H, Man-1), 5.24 – 5.13 (m, 2H), 4.94 – 4.38 (m, 22H), 4.18 – 3.76 (m, 24H), 3.60 – 3.43 (m, 3H), 1.99 (s, 3H, -CH_3_), 1.00 (s, 9H, -C(-CH_3_)_3_) ppm.

*6-Mercaptohexyl α-D-galactopyranosyl-(1→6)-α-D-galactopyranosyl-(1→3)-α-D-mannopyranoside* **(B)**

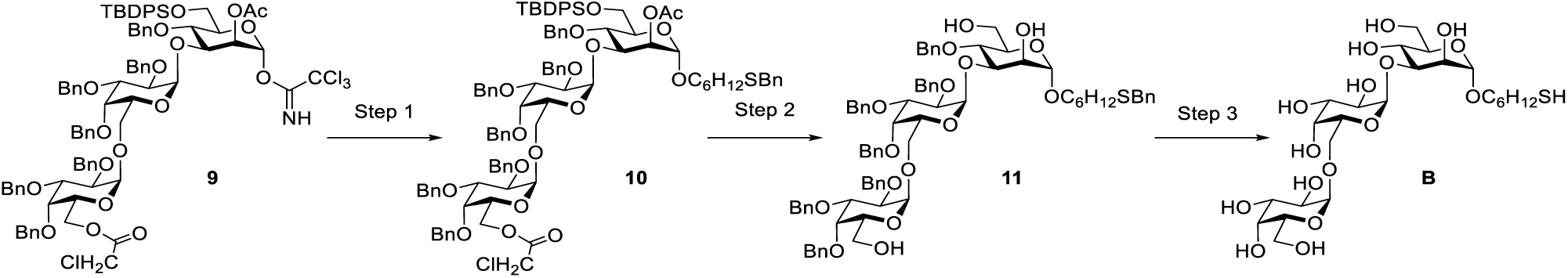

#### Step 1

Imidate **9** (6.110 μmol, 10.0 mg) and 6-thiobenzyl-1-hexanol (0.031 mmol, 7.0 mg) were coevaporated three times with toluene, dried under high vacuum and dissolved in DCM. Freshly activated powdered molecular sieves were added and the suspension was stirred for 10 minutes. TMSOTf (0.002 μmol, 0.4 µl) was added and the reaction was stirred at 0°C. After TLC indicated full conversion, the reaction was quenched by adding saturated NaHCO_3_-solution and extracted with DCM. The combined organic phases were dried over MgSO_4_, filtered and evaporated. The residue was purified by silica gel chromatography giving trisaccharide **10** in 67% yield (4.120 µmol, 7.0 mg) as colorless oil. The compound was directly used for the next step. **R_f_** = 0.50 (4:1, Hex/AcOEt); **^1^H-NMR** (400 MHz, CDCl_3_): δ = 7.74 – 7.60 (m, 4H, H_Ar._), 7.42 – 7.07 (m, 46H, H_Ar._), 5.45 – 4.44 (m, 18H), 4.25 – 3.25 (m, 23H), 2.41 – 2.31 (m, 2H), 2.02 (s, 1.5H, -CH_3_), 1.99 (s, 1.5H, -CH_3_), 1.63 – 1.18 (m, 8H), 1.04 (s, 9H, -C(-CH_3_)_3_) ppm.

#### Step 2

Trisaccharide **10** (4.120 μmol, 7.0 mg) was dissolved in a mixture of DCM and Methanol. Acetyl chloride (0.1 mL) was added dropwise and the reaction was stirred until TLC indicated full conversion. The green reaction solution was diluted with DCM, quenched with saturated NaHCO_3_-solution and extracted with DCM. The combined organic phases were dried over MgSO_4_, filtered and evaporated. The residue was purified by silica gel chromatography giving triol **11** in 83% yield (3.430 µmol, 4.6 mg) as colorless oil. **R_f_** = 0.2 (2:1, Hex/AcOEt); **^1^H-NMR** (400 MHz, CDCl_3_): δ = 7.72 – 7.69 (m, 1H, H_Ar._), 7.53 - 7.51 (m, 1H, H_Ar._), 7.42 – 7.18 (m, 38H, H_Ar._), 5.12 – 3.08 (m, 39H), 2.36 – 2.33 (m, 2H), 1.42 – 1.19 (m, 8H) ppm; **ESI-MS**: m/z M_calcd_ for C_80_H_92_O_16_S = 1340.6106; M_found_ = 1363.5980 [M+Na]^+^; **[α]_D_^20^** = 10.54 (c = 0.1 g/L in CHCl_3_); **FTIR**: ν = 3400.89, 2925.13, 2855.51, 2310.45, 2219.20, 2196.67, 2163.72, 2143.15, 2053.58, 2024.55, 1986.08, 1941.06, 1725.99, 1455.76, 1376.12, 1096.72, 826.02, 737.91, 698.77, 663.35 cm^-1^.

#### Step 3

Ammonia was condensed (10 ml) at -78°C and two drops of methanol were added. Sodium was added in small pieces until a dark blue color was established. Triol **11** (3.430 μmol, 4.6 mg) was dissolved in 1 ml THF and added to the ammonia solution. The reaction was stirred for 1 h, subsequently adding more sodium when the blue color disappeared. The reaction was quenched by adding methanol and ammonia was blown off using a stream of nitrogen. The pH of the resulting solution was adjusted to 7-8 with glacial acetic acid. The residue was concentrated under reduced pressure and purified by size exclusion using 5% ethanol in water as eluent to give the trisaccharide **B** in 71% yield (2.440 µmol, 3.0 mg). **^1^H-NMR** (600 MHz, D_2_O): δ = 5.12 (d, *J* = 4.1 Hz, 1H, Gal-1), 4.71 (s, 1H, Gal’-1), 4.65 (d, *J* = 1.9 Hz, 1H, Man-1), 4.02 – 3.18 (m, 14H), 2.64 (t, *J* = 7.2 Hz, 2H), 1.60 - 1.45 (m, 4H), 1.31 – 1.23 (m, 4H) ppm; **^13^C-NMR** (151 MHz, D_2_O): δ = 100.7, 99.5 (x2), 78.4, 72.6, 71.3, 69.7, 69.2, 68.6, 67.7, 65.9, 61.4, 61.2, 60.7, 38.0, 28.1, 27.0, 24.9 ppm; **ESI-MS**: m/z M_calcd_ for C_24_H_44_O_16_S = 620.2350; M_found_ = 1261.4429 [2M+Na]^+^.

*6-Mercaptohexyl α-D-galactopyranosyl-(1→2)-α-D-galactopyranosyl-(1→6)-2-O-(α-D-galactopyranosyl)-α-D-galactopyranoside* **(C)**

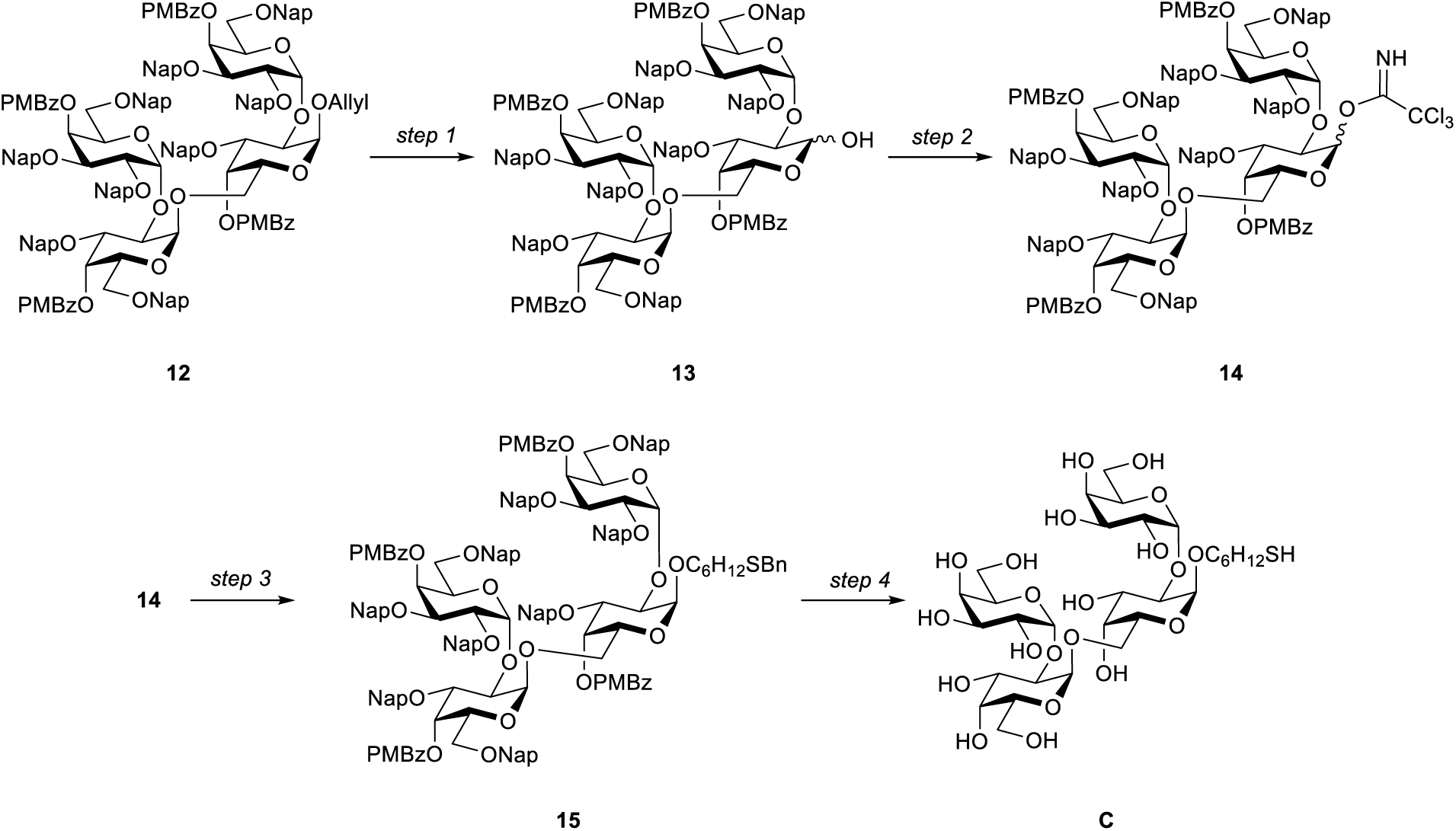

#### Step 1

In a round bottom flask, 10 mg of [IrCOD(PMePh_2_)_2_]PF_6_ were added to 2 mL THF. Hydrogen was bubbled through the suspension until the catalyst dissolved. This solution was transferred to a second flask, where it dissolved of tetrasaccharide **12**[1] (0.080 mmol, 0.200 g). The reaction was stirred at room temperature overnight. THF was evaporated under reduced pressure and the residue was dissolved in an 8:1 mixture of acetone and water. mercury oxide (0.8 µmol, 1.7 mg) and mercury chloride (0.40 mmol, 108.6 mg) were added and the solution was stirred for one hour at room temperature. The reaction was quenched by adding sat. NaHCO_3_ solution and the resulting mixture was extracted three times with DCM. The combined organic phases were dried over Na_2_SO_4_, filtered and concentrated under reduced pressure. The residue was purified by silica column chromatography using hexane and ethyl acetate as eluent. Product **13** was obtained in 81% yield (0.064 mmol, 0.159 g) as colorless oil and directly used for the next step. **R_f_** = 0.1 (3:1, Hex/AcOEt).

#### Step 2

Hemiacetal **13** (0.024 mmol, 0.059 g) was dissolved in DCM at 0°C. Trichloroacetonitril (0.192 mmol, 19.3 µl) and DBU (2.400 µmol, 0.4 µl) were added and the reaction was stirred until TLC indicated full conversion. The resulting mixture was concentrated under reduced pressure and purified by silica column chromatography using hexane and ethyl acetate as eluent. After several reaction cycles Product **14** was obtained in 11% yield (2.680 µmol, 7.000 mg) as colorless oil and used in the next step. **R_f_** = 0.6 (3:1, Hex/AcOEt).

#### Step 3

Imidate **14** (0.019 mmol, 0.050 g) and 6-Thiobenzylhexanol (0.057 mmol. 0.013 g) were co-evaporated three times with toluene and dried under high vacuum. The compound mixture was dissolved in the solvent and 4 Å MS was added. TMSOTf (5.700 µmol, 1.0 µl) was added and the reaction was stirred until TLC indicated full conversion. The reaction mixture was diluted with DCM and quenched by adding sat. NaHCO_3_ solution. The mixture was extracted three times with DCM and the combined organic phases were dried over Na_2_SO_4_, filtered and concentrated under reduced pressure. The residue was purified by silica column chromatography using hexane and ethyl acetate as eluent. Product **15** was obtained in 20% yield (3.740 µmol, 10.0 mg) as colorless oil. **R_f_** = 0.55 (4:1, Hex/AcOEt) **^1^H-NMR** (400 MHz, CDCl_3_): δ = 7.89 – 6.56 (m, 82H, H_Ar._), 5.85 – 5.77 (m, 2H), 5.52 – 5.46 (m, 2H), 5.16 – 5.04 (m, 5H), 4.95 – 3.03 (m, 53H), 2.30 -2.25 (m, 2H), 1.50 -1.13 (m, 8H) ppm. **^13^C-NMR from HSQC** (101 MHz, CDCl_3_): δ = 131.8, 131.7, 127.7 (x), 127.0 (x), 125.9 (x), 113.4 (x), 96.8, 95.7, 95.3, 94.4, 78.8 (x2), 78.4 (x2), 76.5, 76.4, 76.0, 75.9, 75.3, 74.5 (x6), 74.4 (x2), 73.5 (x8), 73.2, 72.7 (x5), 72.3 (x3), 71.9 (x4), 71.8 (x2), 71.6, 68.5 (x4), 68.3 (x2), 68.1 x4), 67.8 (x2), 60.2, 55.3 (x4), 29.8, 22.6, 20.9, 14.1 (x2) ppm.

#### Step 4

10 mL ammonia were condensed in a flask and methanol (2 drops) was added. Sodium was added in small pieces until a dark blue color established. Tetrasaccharide **15** (3.700 µmol, 10.0 mg) was dissolved in THF and added to the ammonium solution at -78°C. At this temperature the reaction was stirred for 1 h. The reaction was quenched by adding methanol and ammonia was blown off using a stream of nitrogen. The pH of the resulting solution was adjusted with glacial acetic acid to 7-8. The reaction concentrated under reduced pressure and the residue was purified by size exclusion using 5% ethanol in water as eluent. A final HPLC purification on a hypercarb column (150×10mm, ThermoFisher, 5 µ) using a 0-100% gradient of acetonitrile in water in 60 min delivered product **C** in 20% (7.480 µmol, 5.860 mg) as white solid. **^1^H-NMR** (400 MHz, D_2_O): δ = 5.14 – 4.99 (m, 4H), 4.07 – 3.44 (m, 16H), 2.82 – 2.77 (m, 2H), 1.68 – 1.28 (m, 4H) ppm. **ESI-MS**: m/z M_calcd_ for C_30_H_54_O_21_S = 782.2878; M_found_ = 1581.6877 [M_2_+NH_4_]^+^.

*6-(Benzylthio)-hexyl 2-O-acetyl-3,4-di-O-benzyl-6-hydroxy-α-D-mannopyranoside (**18**)*

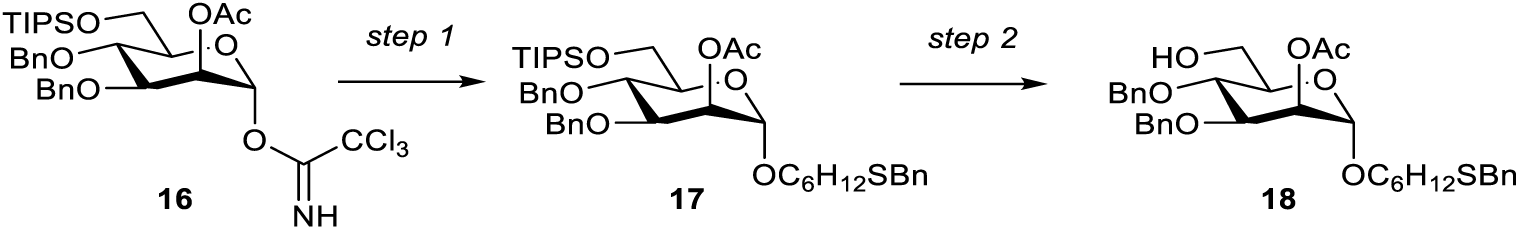

#### Step 1

A mixture of mannosyl trichloroacetamidate donor **16**[2] (1.138 mmol, 0.800 g) and 6-Thiobenzylhexanol (3.41 mmol, 0.766 g) was coevaporated three times with toluene and dried under high vacuum for 1 h. After that the reaction mixture was dissolved in DCM followed by the addition of molecular sieves. The reaction stirred for 10 min at 0°C and TMSOTf (0.228 mmol, 41.0 µl) was added. After 1.30 h, the reaction mixture was quenched using triethylamine, filtered and concentrated. The residue was purified using silica gel column chromatography to obtain **17** in 72% yield (0.819 mmol, 625 mg) as colorless oil. **R_f_** = 0.6 (4:1, Hex/AcOEt), **^1^H-NMR** (400 MHz, CDCl_3_): δ = 7.26 – 7.08 (m, 15H), 5.20 (dd, *J* = 3.3, 1.8 Hz, 1H), 4.79 (d, *J* = 10.7 Hz, 1H), 4.64 – 4.58 (m, 2H), 4.53 (d, *J* = 10.7 Hz, 1H), 4.45 (d, *J* = 11.1 Hz, 1H), 3.95 – 3.74 (m, 4H), 3.59 (d, *J* = 2.4 Hz, 2H), 3.52 (qd, *J* = 6.7, 3.7 Hz, 2H), 3.22 (dt, *J* = 9.7, 6.6 Hz, 1H), 2.29 (dd, *J* = 9.3, 5.4 Hz, 2H), 2.00 (s, 3H), 1.42 (dt, *J* = 14.5, 7.7 Hz, 4H), 1.25 – 1.16 (m, 4H), 0.97 (d, *J* = 4.6 Hz, 21H) ppm; **^13^C-NMR** (101 MHz, CDCl_3_): δ = 170.6, 138.6, 138.5, 138.0, 128.8, 128.4, 128.4, 128.4, 128.3, 128.1, 128.0, 127.7, 127.6, 126.8, 97.3, 78.2, 75.3, 74.2, 72.8, 71.8, 69.0, 67.4, 62.7, 36.2, 31.2, 29.2, 29.0, 28.6, 25.7, 21.1, 18.0, 17.9, 12.0 ppm; **ESI-MS**: m/z M_calcd_ for C_44_H_64_O_7_SSi = 764.4142; M_found_ = 787.4054 [M+Na]^+^.

#### Step 2

To a solution of **17** (0.784 mmol, 0.6 mg) in ACN (10 ml) and DCM (5 ml), water (100 μl) and Sc(OTf)_3_ (2.353 mmol, 1.2 mg) were added and the solution was heated to 50°C for 3 h. The reaction was quenched with pyridine (100 μl) and the solvents were removed *in vacuo*. The residue was co-evaporated with toluene and purified by silica gel column chromatography to obtain **18** in 75% yield (0.588 mmol, 0.360 g) as colorless oil. **R_f_** = 0.2 (4:1, Hex/EtOAc); **^1^H-NMR** (400 MHz, CDCl_3_): δ = 7.48 – 6.88 (m, 15H), 5.34 (dd, *J* = 3.4, 1.8 Hz, 1H), 4.90 (d, *J* = 10.9 Hz, 1H), 4.77 – 4.65 (m, 2H), 4.61 (d, *J* = 10.8 Hz, 1H), 4.53 (d, *J* = 11.2 Hz, 1H), 3.97 (dd, *J* = 9.3, 3.3 Hz, 1H), 3.79 (qd, *J* = 11.9, 5.4 Hz, 3H), 3.71 – 3.57 (m, 4H), 3.34 (dt, *J* = 9.6, 6.5 Hz, 1H), 2.39 (t, *J* = 7.3 Hz, 2H), 2.13 (s, 3H), 1.53 (q, *J* = 7.4 Hz, 4H), 1.39 – 1.23 (m, 4H) ppm; **^13^C-NMR from HSQC** (101 MHz, CDCl_3_, coupled): δ = 128.8, 128.4, 128.4, 128.4, 128.3, 128.1, 128.0, 127.7, 127.6, 126.8, 97.9, 78.1, 75.3, 74.3, 71.8, 71.7, 68.8, 67.8, 62.1, 36.2, 31.3, 29.3, 28.6, 26.0, 21.3 ppm; **ESI-MS**: m/z M_calcd_ for C_35_H_44_O_7_S = 608.2808; M_found_ = 631.2714 [M+Na]^+^.

6-(Benzylthio)-hexyl 2-*O*-acetyl-3,4-*O*-benzyl-6-hydroxy-α-D-mannopyranosyl-(1→2)-3,4,6-tri-*O*-benzyl-α-D-mannopyranosyl-(1→6)-2-*O*-acetyl-3,4-di-*O*-benzyl-α-D-mannopyranoside (**21**)

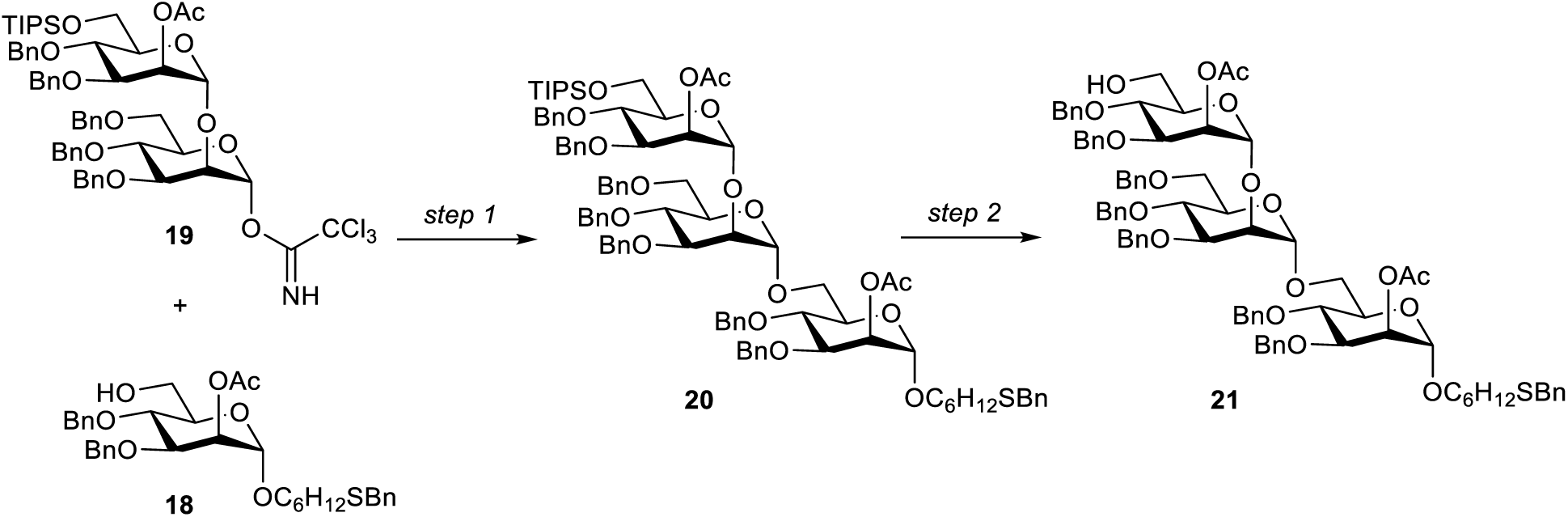

#### Step 1

A mixture of trichloroacetamidate donor **19**[2] (0.411 mmol, 0.466 g) and acceptor **18** (0.411 mmol, 0.250 g) was coevaporated three times with toluene and dried under high vacuum for 1 h. After that the reaction mixture was dissolved in a 2:1 mixture of thiophene and DCM followed by addition of molecular sieves. The reaction was stirred for 10 min at 0°C and TBSOTf (0.082 mmol, 20.0 µl) was added. After 1 h, the reaction was quenched using triethylamine, filtered and concentrated. The residue was purified using silica gel column chromatography to obtain trisaccharide **20** in 46% yield (0.189 mmol, 0.300 g) as colorless oil. **R_f_** = 0.6 (6:1, Hex/AcOEt); **^1^H-NMR** (400 MHz, CDCl_3_): δ = 7.38 – 6.88 (m, 40H), 5.46 – 5.20 (m, 2H), 5.06 (s, 1H), 4.86 – 4.73 (m, 4H), 4.69 – 4.47 (m, 7H), 4.46 – 4.29 (m, 5H), 4.08 (s, 1H), 3.99 – 3.81 (m, 7H), 3.77 (t, *J* = 9.4 Hz, 1H), 3.69 – 3.59 (m, 6H), 3.54 – 3.42 (m, 4H), 3.30 – 3.20 (m, 1H), 2.32 (t, *J* = 7.3 Hz, 2H), 2.02 (s, 3H), 2.00 (s, 3H), 1.52 – 1.40 (m, 4H), 1.28 – 1.16 (m, 4H), 1.00 (d, *J* = 4.6 Hz, 21H) ppm; **^13^C-NMR** (101 MHz, CDCl_3_): δ = 170.4, 170.2, 138.8, 138.6, 138.5, 138.4, 138.3, 138.1, 138.0, 137.8, 128.8, 128.5, 128.4, 128.4, 128.3, 128.2, 128.2, 127.9, 127.8, 127.8, 127.7, 127.6, 127.6, 127.5, 127.5, 127.3, 126.9, 98.9 (x2), 97.6, 79.6, 78.6, 77.9, 77.2, 75.2, 75.0, 74.9, 74.3, 73.9, 73.8, 73.3, 73.2, 73.1, 71.8, 71.7, 71.5, 70.5, 68.9, 68.9, 68.6, 67.7, 66.1, 62.5, 36.2, 31.2, 29.2, 29.0, 28.6, 25.8, 21.1, 21.0, 18.0, 17.9, 12.0 ppm; **ESI-MS**: m/z M_calcd_ for C_93_H_116_O_18_SSi = 1580.7652; M_found_ = 1603.7544 [M+Na]^+^.

#### Step 2

To a solution of trimannose **20** (0.190 mmol, 0.300 mg) in ACN (5 ml) and DCM (3 ml), water (50 μl) and Sc(OTf)_3_ (0.569 mmol, 0.280 g) were added and the solution was heated to 50 °C for 6 h. The reaction was quenched with pyridine (50 μl) and the solvents were removed *in vacuo*. The residue was co-evaporated with toluene and purified through silica gel column chromatography to obtain alcohol **21** in 70% yield (0.133 mmol, 190.0 mg) as colorless oil. **R_f_** = 0.2 (4:1, Hex/AcOEt); **^1^H-NMR** (400 MHz, CDCl_3_): δ = 7.37 – 7.06 (m, 40H), 5.49 (t, *J* = 2.4 Hz, 1H), 5.37 – 5.31 (m, 1H), 5.00 (s, 2H), 4.90 – 4.79 (m, 3H), 4.75 – 4.56 (m, 7H), 4.52 – 4.35 (m, 5H), 4.04 (d, *J* = 2.2 Hz, 1H), 3.99 – 3.80 (m, 6H), 3.79 – 3.47 (m, 12H), 3.32 (dd, *J* = 9.5, 6.5 Hz, 1H), 2.38 (t, *J* = 7.3 Hz, 2H), 2.10 (s, 3H), 2.09 (s, 3H), 1.51 (dt, *J* = 13.9, 7.0 Hz, 4H), 1.37 – 1.28 (m, 4H) ppm; **ESI-MS**: m/z M_calcd_ for C_84_H_96_O_18_S = 1424.6317; M_found_ = 1447.6229 [M+Na]^+^.

*Triethylammonium 2-O-acetyl-3,4-O-benzyl-6-O-(2-N-benzyloxycarbonyl)aminoethyl-phosphonato-α-D-mannopyranosyl-(1→2)-3,4,6-tri-O-benzyl-α-D-mannopyranosyl-(1→6)-1-O-(6-thiobenzyl)hexyl-2-O-acetyl-3,4-O-benzyl-α-D-mannopyranose* (**D**)

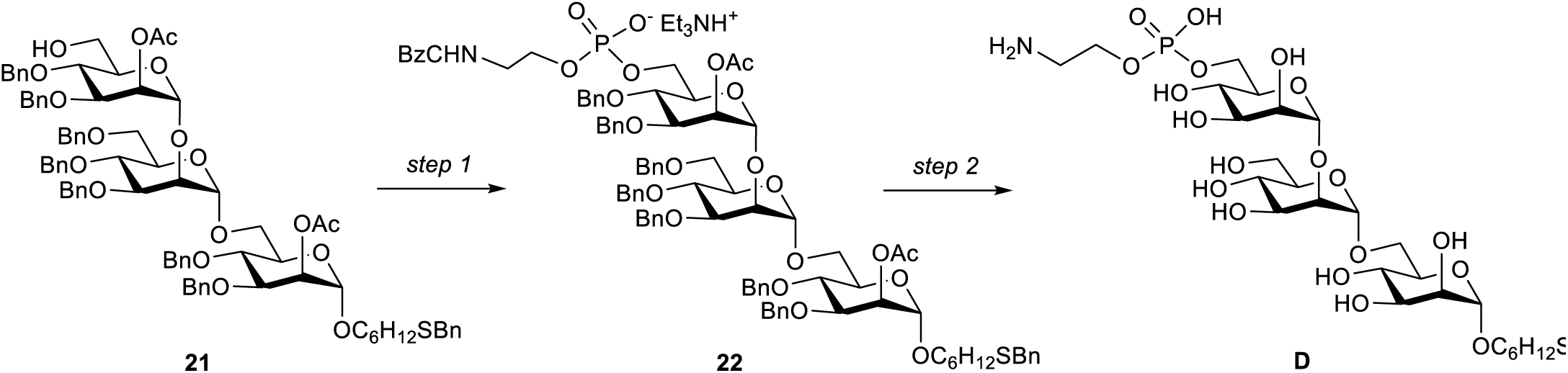

#### Step 1

Alcohol **21** (0.088 mmol, 0.125 g) and 2-amino-(benzyloxy-carbonyl)-H-phosphonate (0.438 mmol, 0.105 g) were co evaporated three times with pyridine. The residue was dissolved in pyridine (2 ml) and PivCl (0.263 mmol, 33.0 μl) was added dropwise. The solution was stirred for 2 h at room temperature before water (15 μl) and iodine (0.316 mmol, 80.0 mg) were added. The red solution was stirred for 1 h and quenched with sat. Na_2_S_3_O_3_ solution. The reaction mixture was diluted with chloroform and dried over Na_2_SO_4_. The reaction mixture was filtered and concentrated *in vacuo*. The crude residue was purified through flash column chromatography on deactivated (1% TEA in CHCl_3_) silica gel using methanol and chloroform an eluent to give phosphate **22** as yellow oil in 85% yield (0.075 mmol, 125.0 mg). **R_f_** = 0.4 (10% MeOH in DCM); **^1^H-NMR** (600 MHz, CDCl_3_): δ = 7.37 – 7.07 (m, 45H), 6.65 (t, *J* = 5.0 Hz, 1H), 5.50 (t, *J* = 2.1 Hz, 1H), 5.37 (dd, *J* = 3.4, 1.8 Hz, 1H), 5.09 – 5.00 (m, 3H), 4.94 (d, *J* = 2.0 Hz, 1H), 4.90 – 4.80 (m, 4H), 4.79 – 4.67 (m, 3H), 4.64 – 4.53 (m, 4H), 4.52 – 4.42 (m, 3H), 4.37 (dd, *J* = 11.7, 5.4 Hz, 2H), 4.27 (dt, *J* = 11.6, 3.8 Hz, 1H), 4.18 – 4.11 (m, 1H), 4.09 (t, *J* = 2.3 Hz, 1H), 4.02 – 3.86 (m, 7H), 3.82 (t, *J* = 9.6 Hz, 1H), 3.79 – 3.74 (m, 2H), 3.74 – 3.69 (m, 1H), 3.68 (s, 2H), 3.61 (dt, *J* = 9.8, 6.3 Hz, 2H), 3.56 (dd, *J* = 11.2, 4.7 Hz, 1H), 3.52 – 3.48 (m, 1H), 3.40 – 3.32 (m, 3H), 2.39 (t, *J* = 7.4 Hz, 2H), 2.11 (s, 3H), 2.07 (s, 3H), 1.57 – 1.48 (m, 4H), 1.37 – 1.23 (m, 4H) ppm; **^13^C-NMR** (151 MHz, CDCl_3_): δ = 170.4, 170.0, 156.6, 138.9, 138.6, 138.6, 138.5, 138.2, 138.1, 137.9, 137.0, 128.8, 128.4, 128.4, 128.3, 128.3, 128.2, 128.2, 128.2, 128.1, 128.0, 127.9, 127.8, 127.7, 127.7, 127.6, 127.5, 127.4, 127.4, 127.4, 127.3, 126.8, 99.5, 98.8, 97.6, 79.3, 78.6, 77.9, 74.9, 74.9, 74.6, 74.4, 74.0, 73.9, 73.1, 71.8, 71.8, 71.7, 71.6, 71.6, 70.6, 69.0, 68.9, 68.7, 67.8, 66.3, 66.2, 64.3, 64.3, 64.2, 42.5, 36.3, 31.3, 29.3, 29.1, 28.7, 25.8, 21.1, 21.1 ppm; **^31^P-NMR** (243 MHz, CDCl_3_): δ = 1.29 ppm; **ESI-MS**: m/z M_calcd_ for C_100_H_121_N_2_O_22_PS = 1764.7869; M_found_ = 1783.8053 [M+H_3_O]^+^.

#### Step 2

Phosphate **22** (0.030 mmol, 50.0 mg) was dissolved in THF (2 mL) and *tert-*butanol (2 drops). This solution was added to approximately 10 ml ammonia which was condensed at -78°C. Small fresh cut pieces of sodium were added till a dark blue color was established. The reaction was stirred for 35 min at -78°C. The reaction was quenched with MeOH (2 mL) and ammonia was blown off using a stream of nitrogen. The solution was adjusted with concentrated acetic acid to pH 7. Water was removed by freeze drying and the residue was purified using a Sephadex super fine G-25 (GE Healthcare) column (1 cmx20 cm) to yield compound **D** as a white solid mixture of free thiol and disulphide in 35% yield (0.011 mmol, 19.0 mg). **^1^H-NMR** (400 MHz, D_2_O): δ = 4.97 (s, 1.5H), 4.89 (s, 1.5H), 4.70 (s, 1.5H), 4.03 – 3.93 (m, 8H), 3.85 – 3.71 (m, 10H), 3.67 – 3.53 (m, 12H), 3.42 (dt, *J* = 10.9, 6.5 Hz, 2H), 3.14 (t, *J* = 4.9 Hz, 3H), 2.64 (t, *J* = 7.2 Hz, 1H), 2.41 (t, *J* = 7.1 Hz, 2H), 1.49 (q, *J* = 6.8 Hz, 6H), 1.32 – 1.12 (m, 6H) ppm; **^13^C-NMR** (101 MHz, D_2_O): δ = 102.3, 99.8, 97.9, 78.9, 72.7, 72.0, 71.9, 71.1, 70.7, 70.1, 70.0, 69.8, 67.87, 66.8, 66.7, 66.2, 66.1, 64.7, 61.8, 60.8, 40.0, 32.8, 28.3, 27.1, 24.8, 23.6 ppm; **^31^P-NMR** (162 MHz, D_2_O): δ = 0.25 ppm; **ESI-MS**: m/z M_calcd_ for C_26_H_50_NO_19_PS = 743.2435; M_found_ = 766.2354 [M+Na]^+^.

#### *VSG117 CTD peptide (KGKLEDTCKKESNCKWENNA)* (F)

In a fritted reaction vessel 200 mg trityl-ChemMatrix® resin (substitution grade 0.62 mmol/g) were swollen in anhydrous DCM for 2 hours. After DCM was drained, a solution of 10% AcBr in anhydrous DCM (14.00 ml) was added to the resin and the slurry was shaken for 4 hours. The resin was washed numerous times with anhydrous DCM and a solution of 350 mg Fmoc-Thr(OtBu)-OH and 400 μ l DIPEA in 10 ml anhydrous DCM was added to the resin and shaken for 16 hours. The resin was washed neatly with DCM and the efficiency of the coupling was determined by Fmoc quantification. The resin was capped using a mixture of methanol, DIPEA and DCM (2:1:17) for 10 minutes. Coupling reagents used were DIC and Oxyma. The coupling reagents were prepared as solutions in DMF with 1 M Oxyma (with 0.1 M DIPEA) and 0.5 M DIC. Amino acids were added as 0.2 M solutions in DMF. All amino acids were coupled twice in five-fold excess. The temperature during coupling is 50°C and the coupling time is 10 minutes, except arginine. Arginine was carried out at room temperature for 20 minutes and the second coupling was at 50°C for 10 minutes. For all amino acids Fmoc was removed three times using 20% piperidine in DMF without microwave for 5 minutes. TFA/TIPS/water (190:5:5/v:v:v) was added and the resin (100 mg when synthesis was started) was shaken for 3 hours. The cleavage solution was collected and the resin was washed with another 8 ml TFA. The solution was concentrated under nitrogen and crushed out with ice cold diethyl ether. The precipitate was centrifuged and washed two more times with ice cold diethyl ether. The resulting peptide was dried, dissolved, lyophilized and purified over RP-HPLC. MALDI: M_calcd_ for oxidized peptide = 2325.584; M_found_ = 2358.070

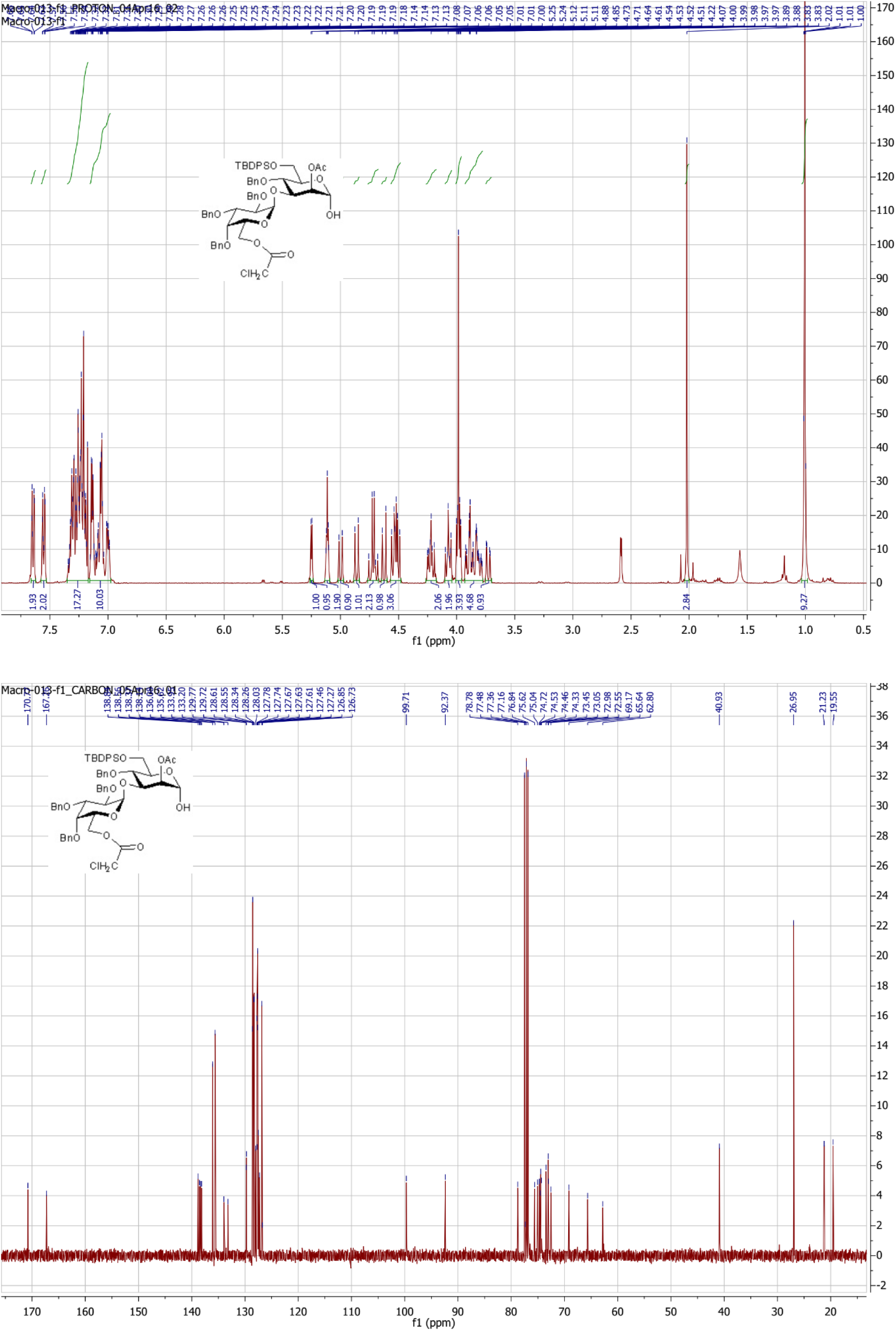
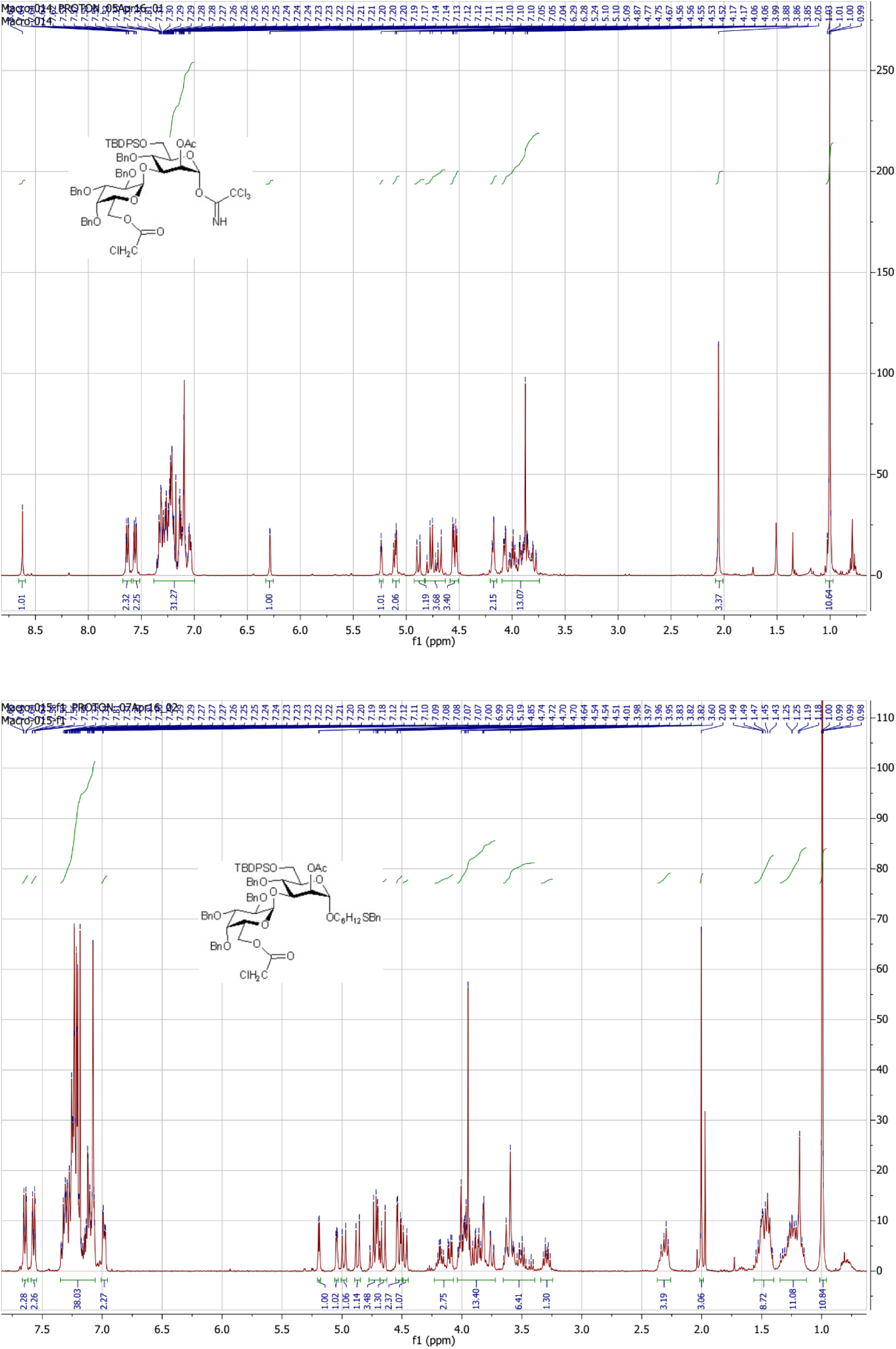
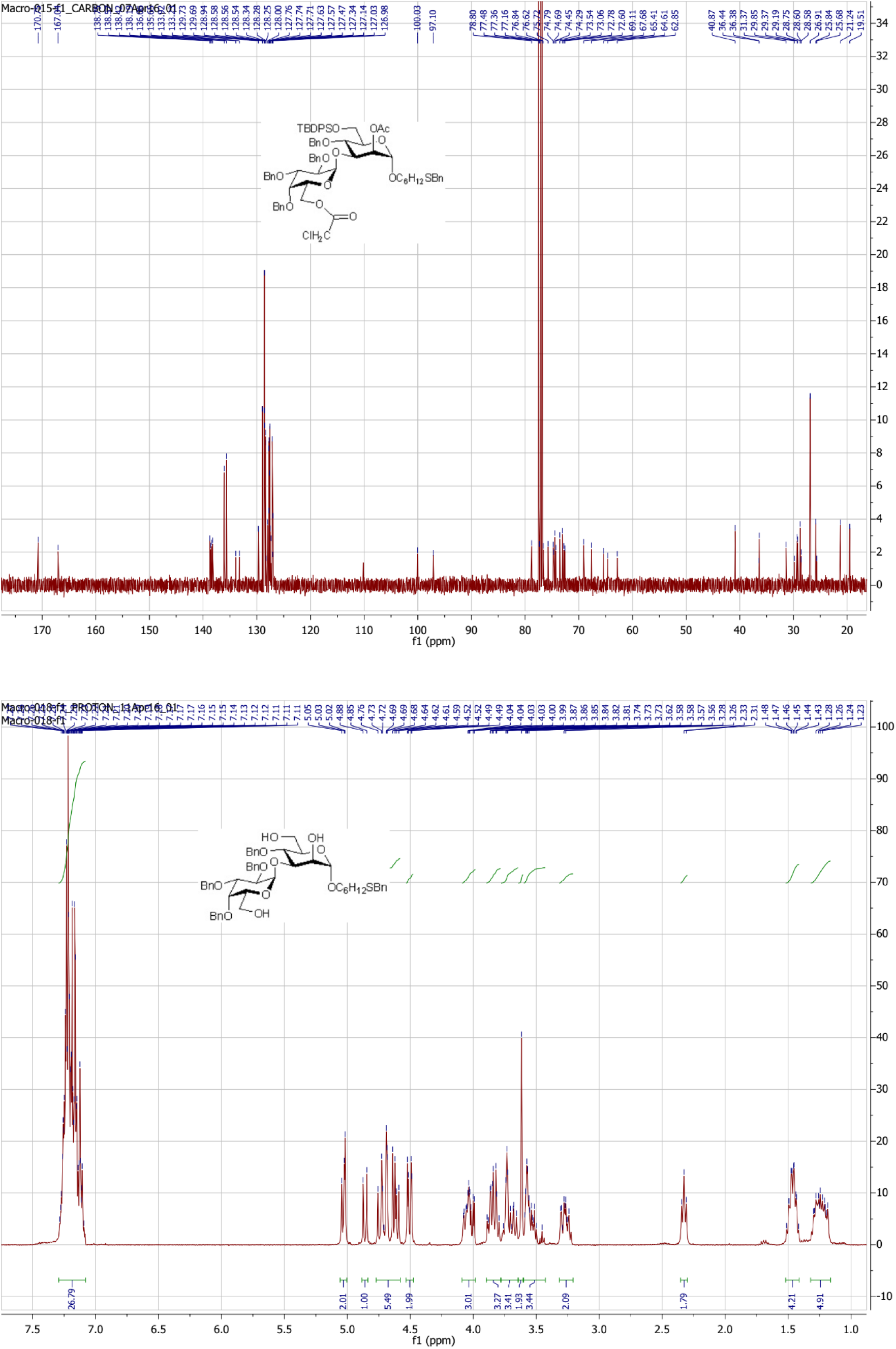
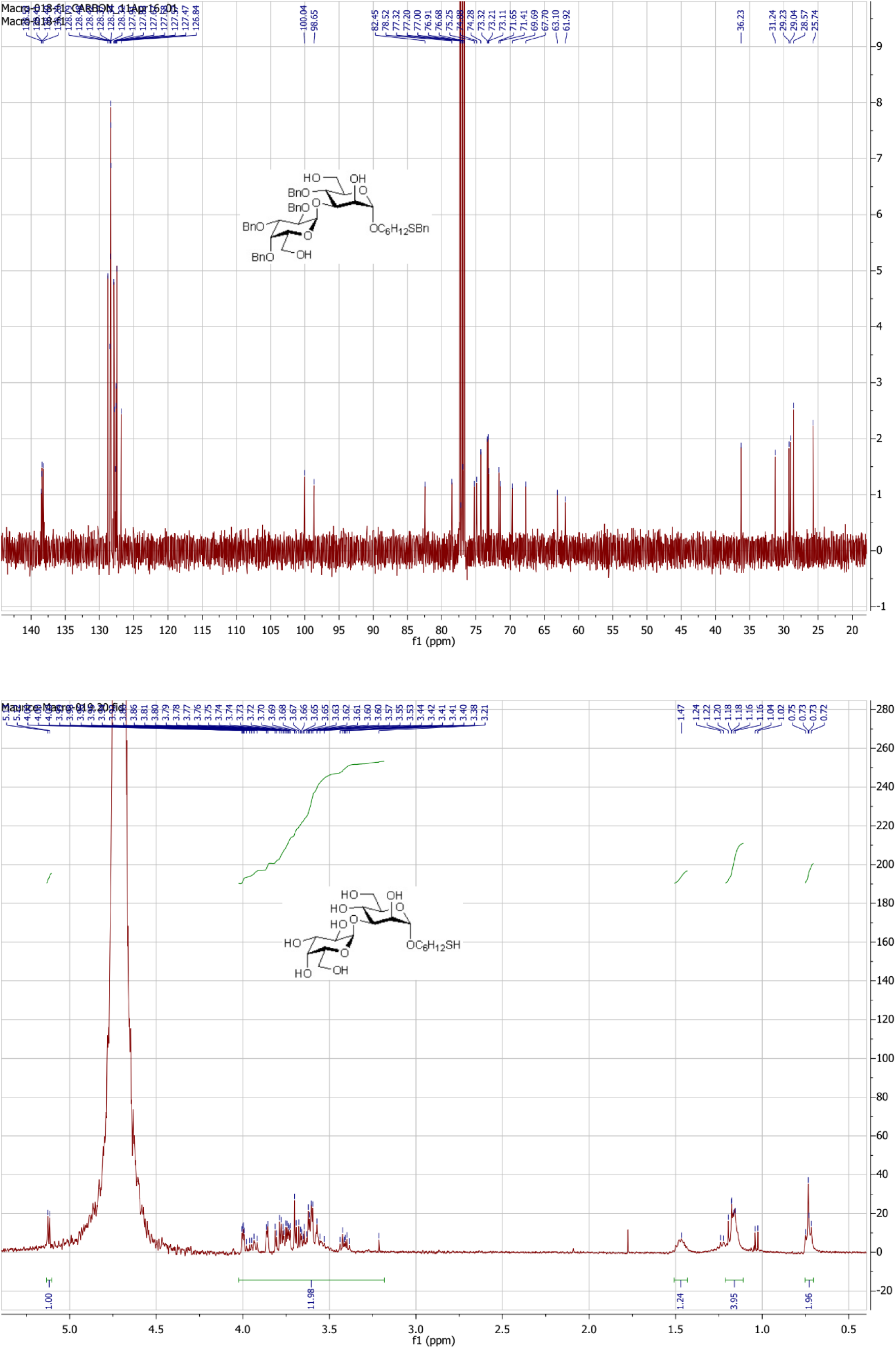
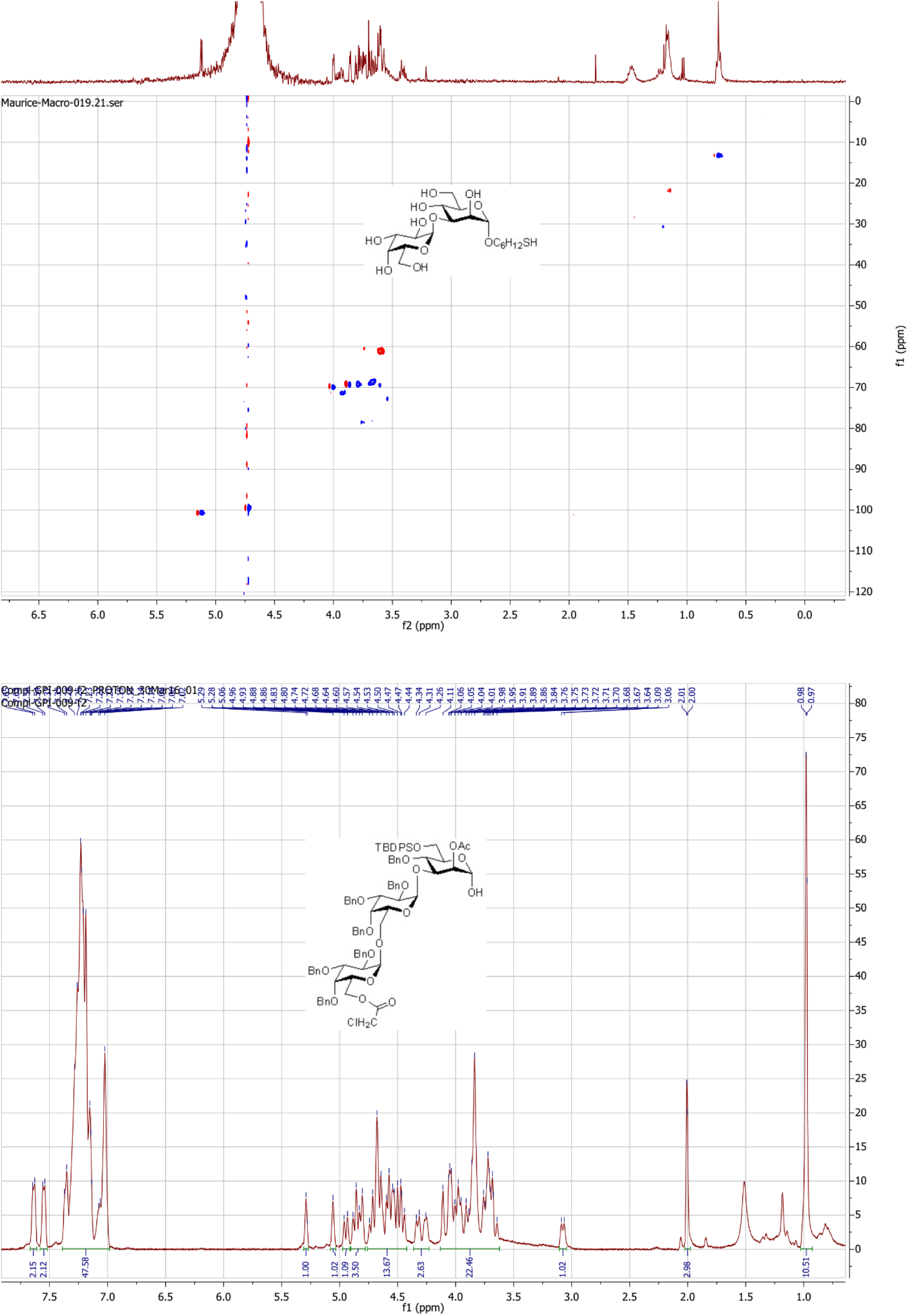
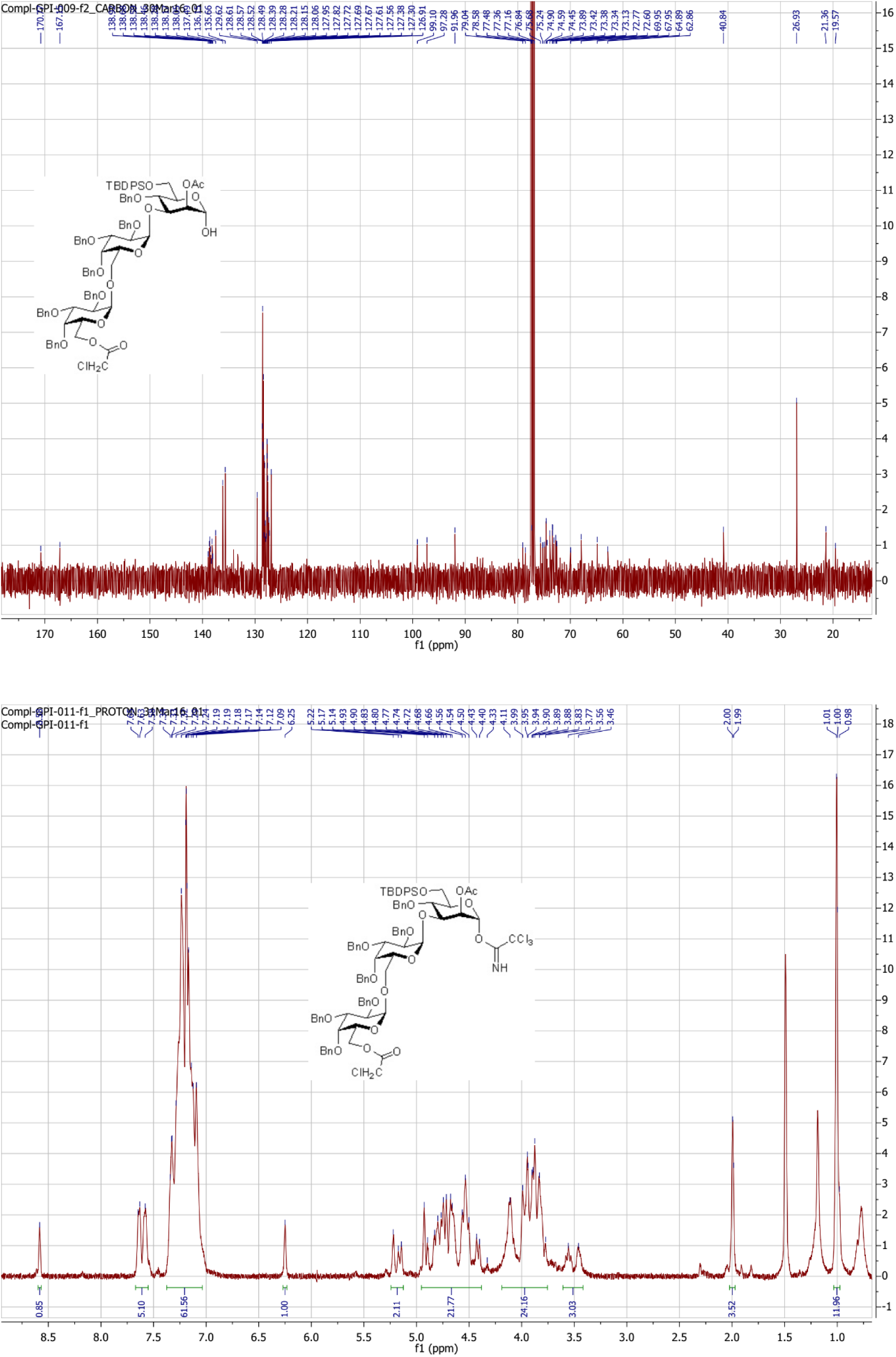
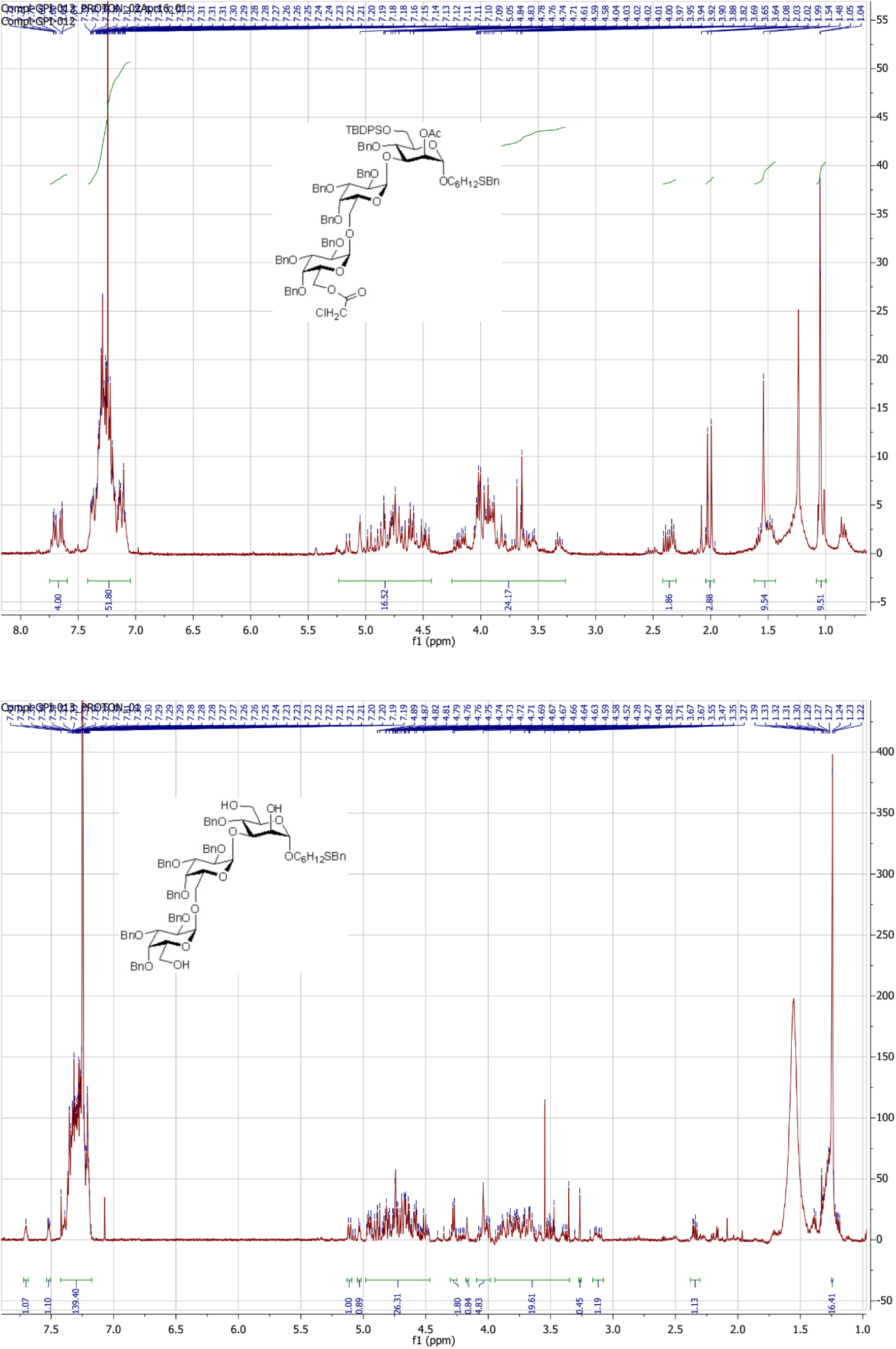
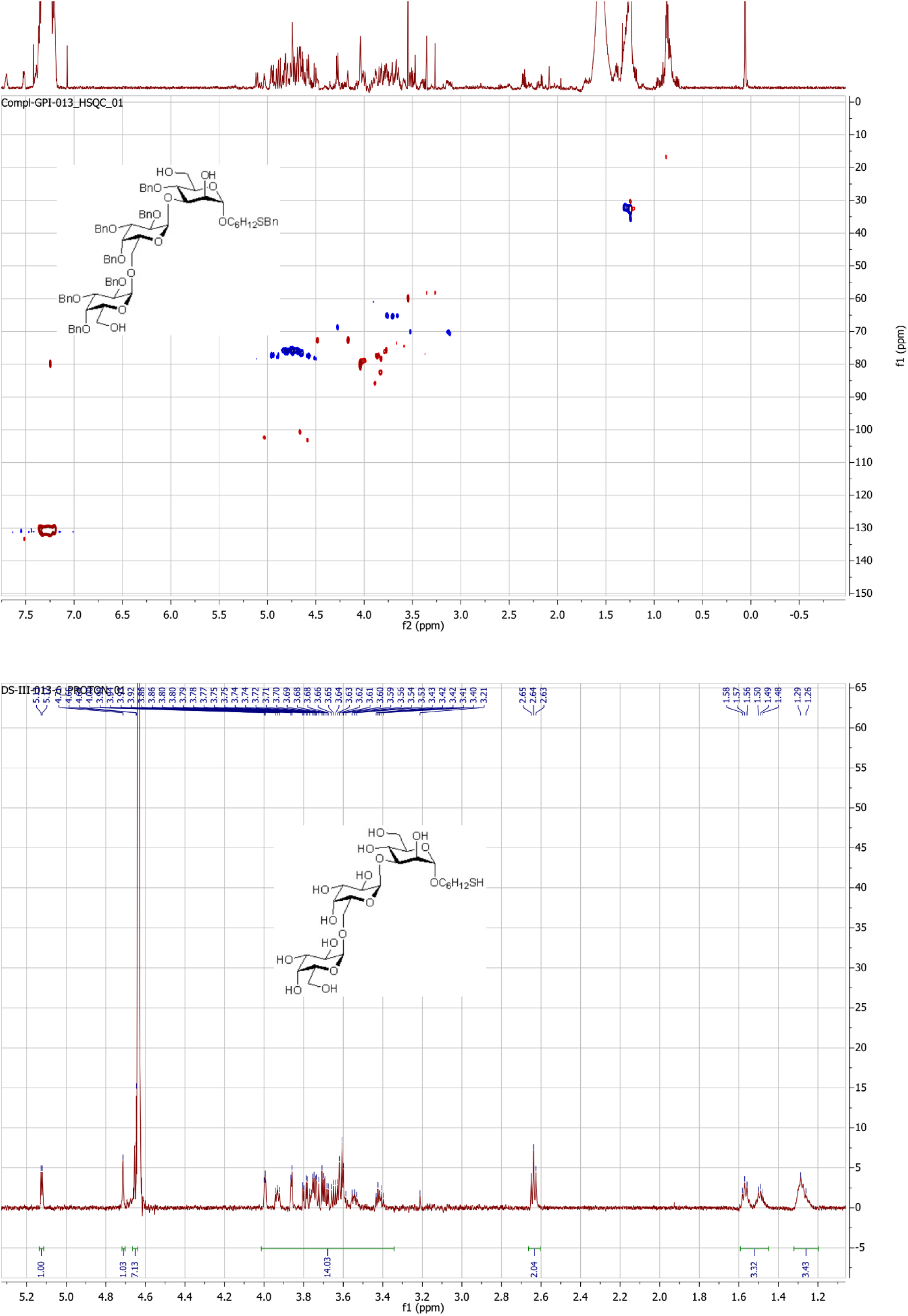
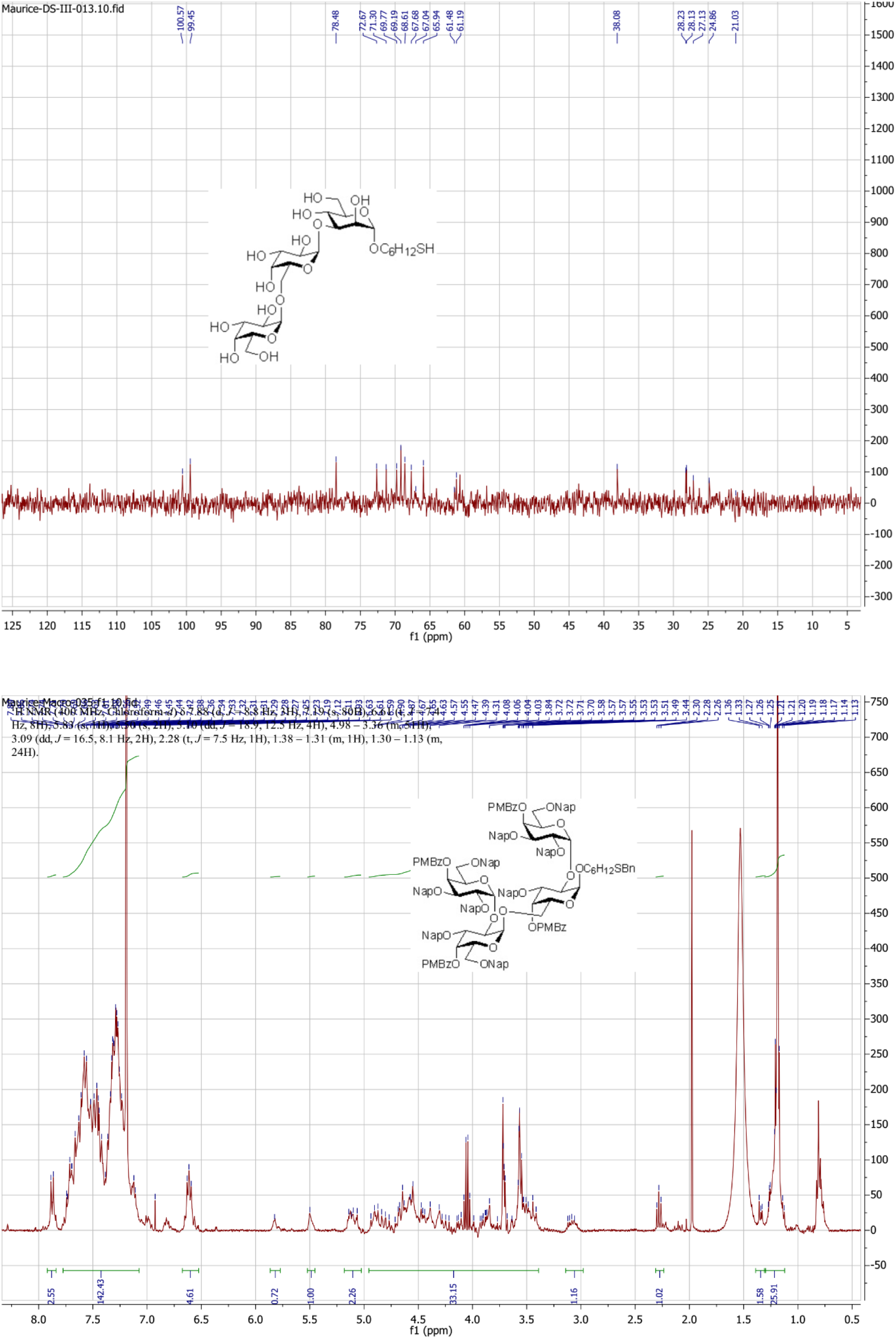
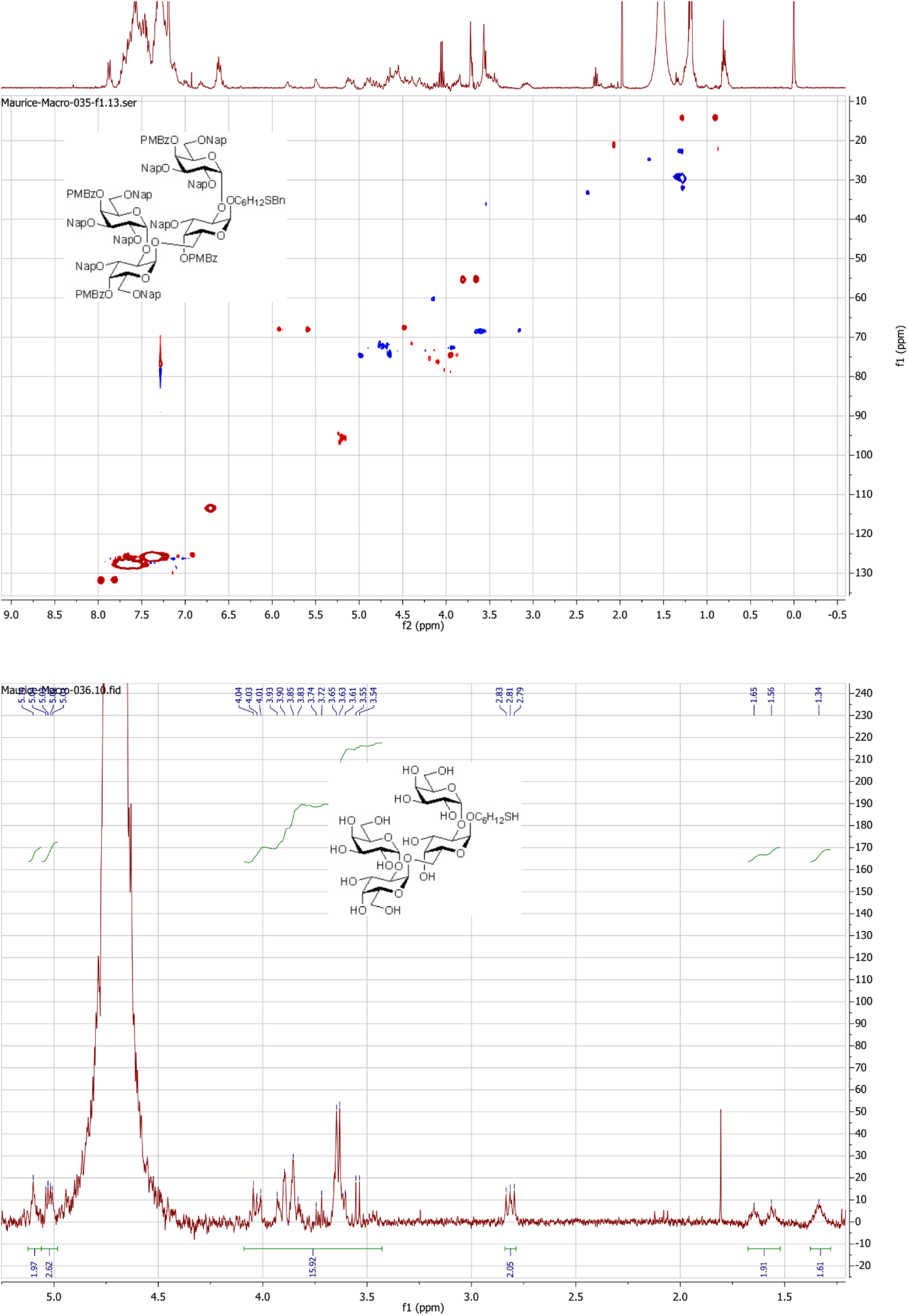
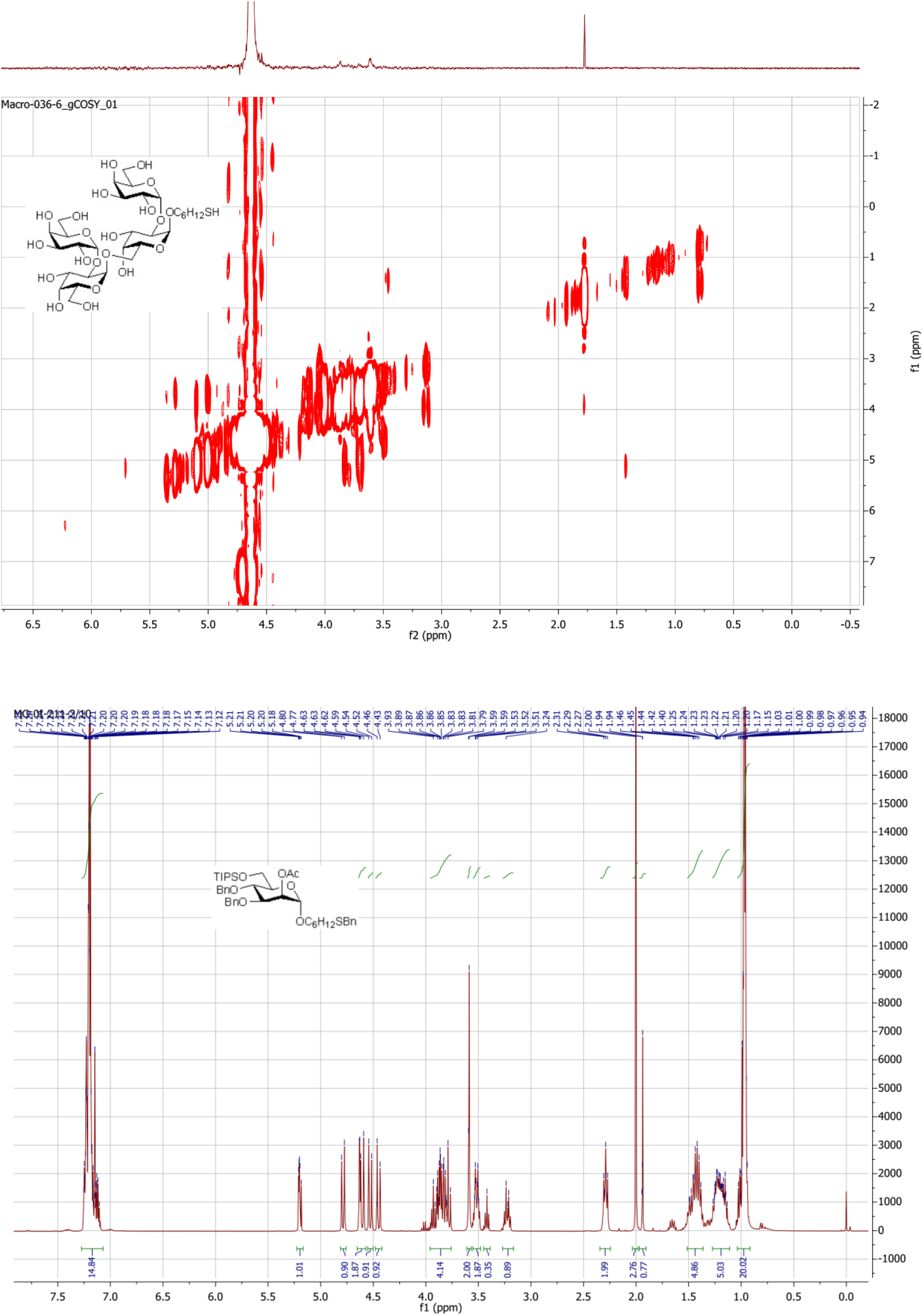
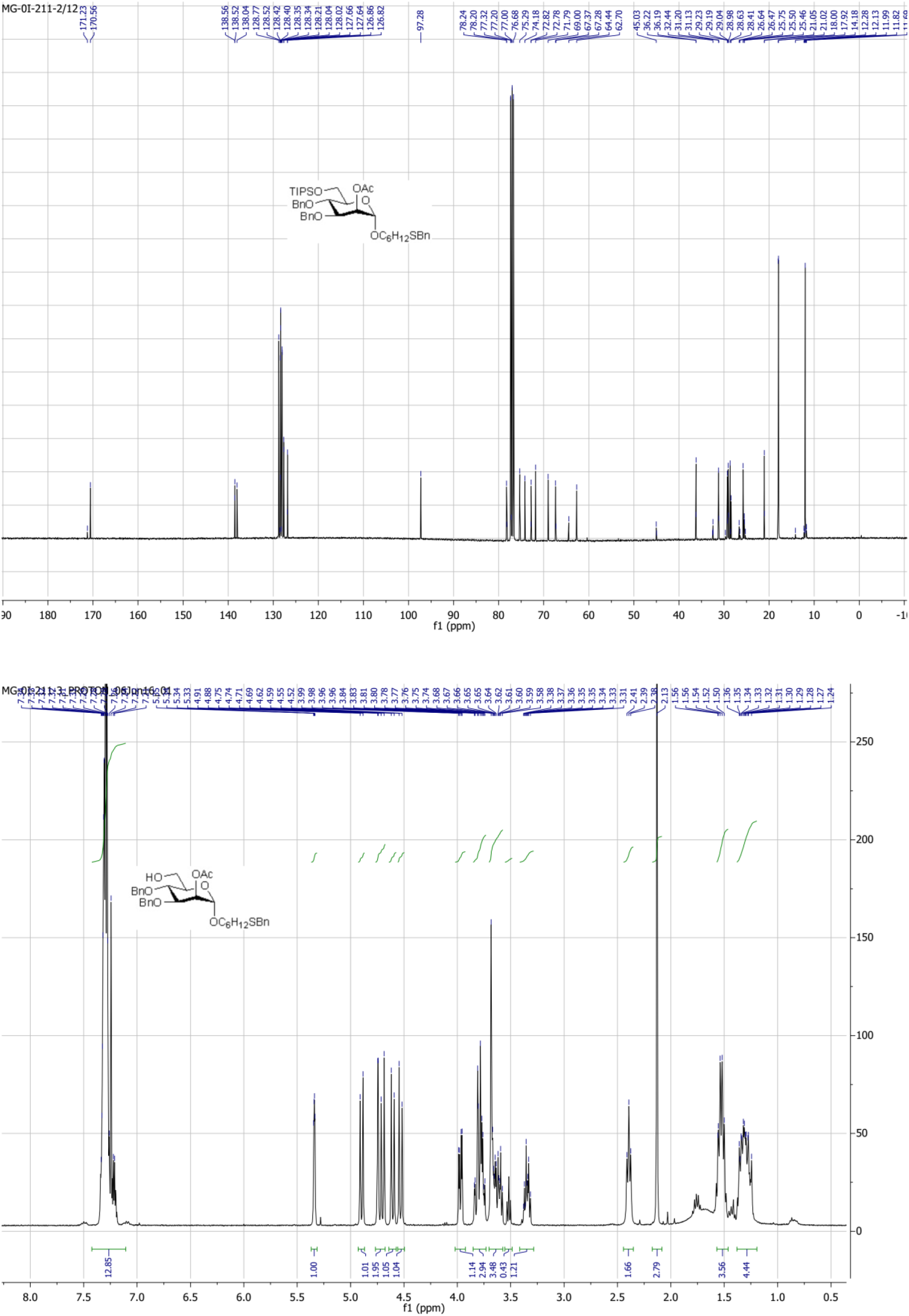
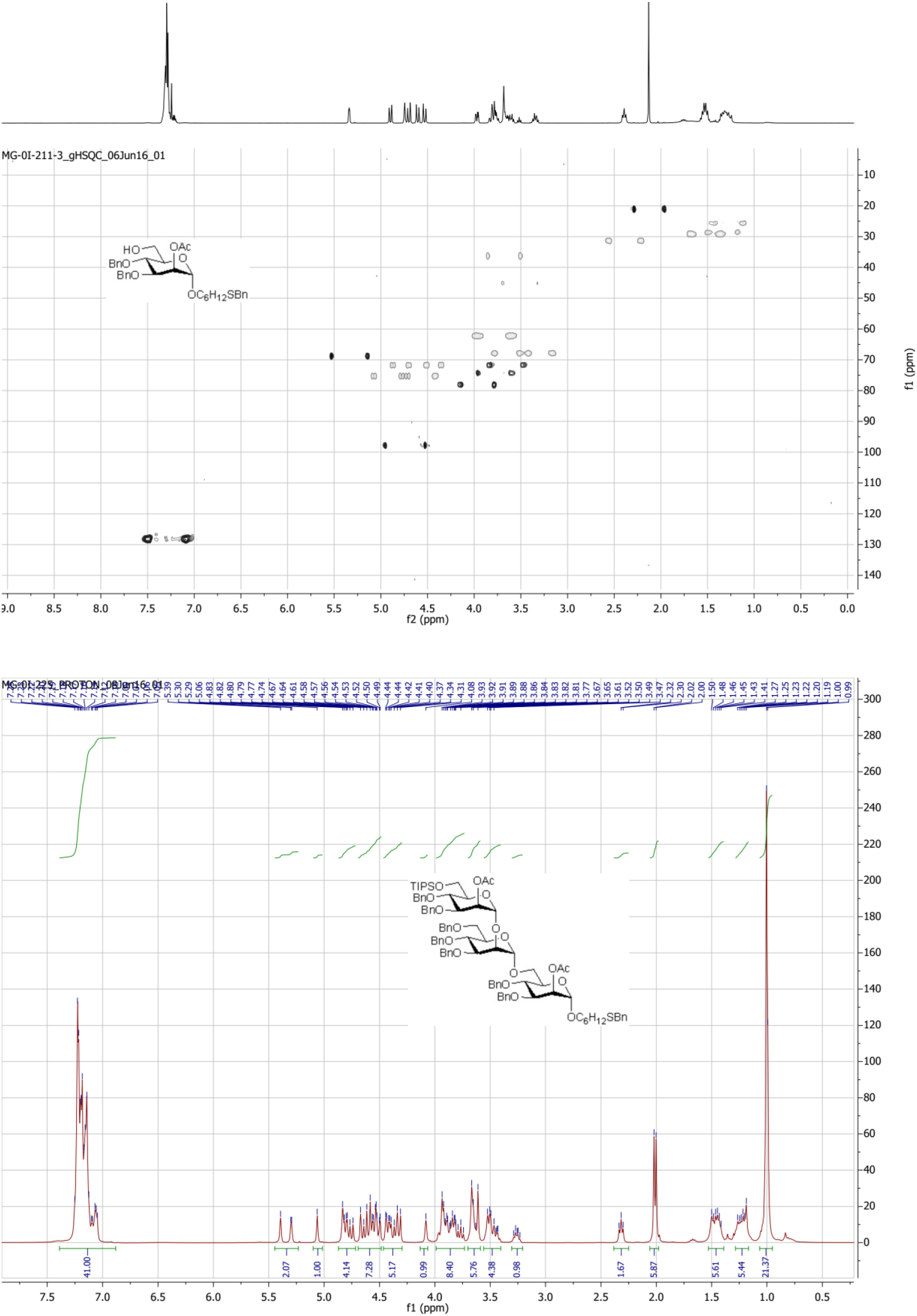
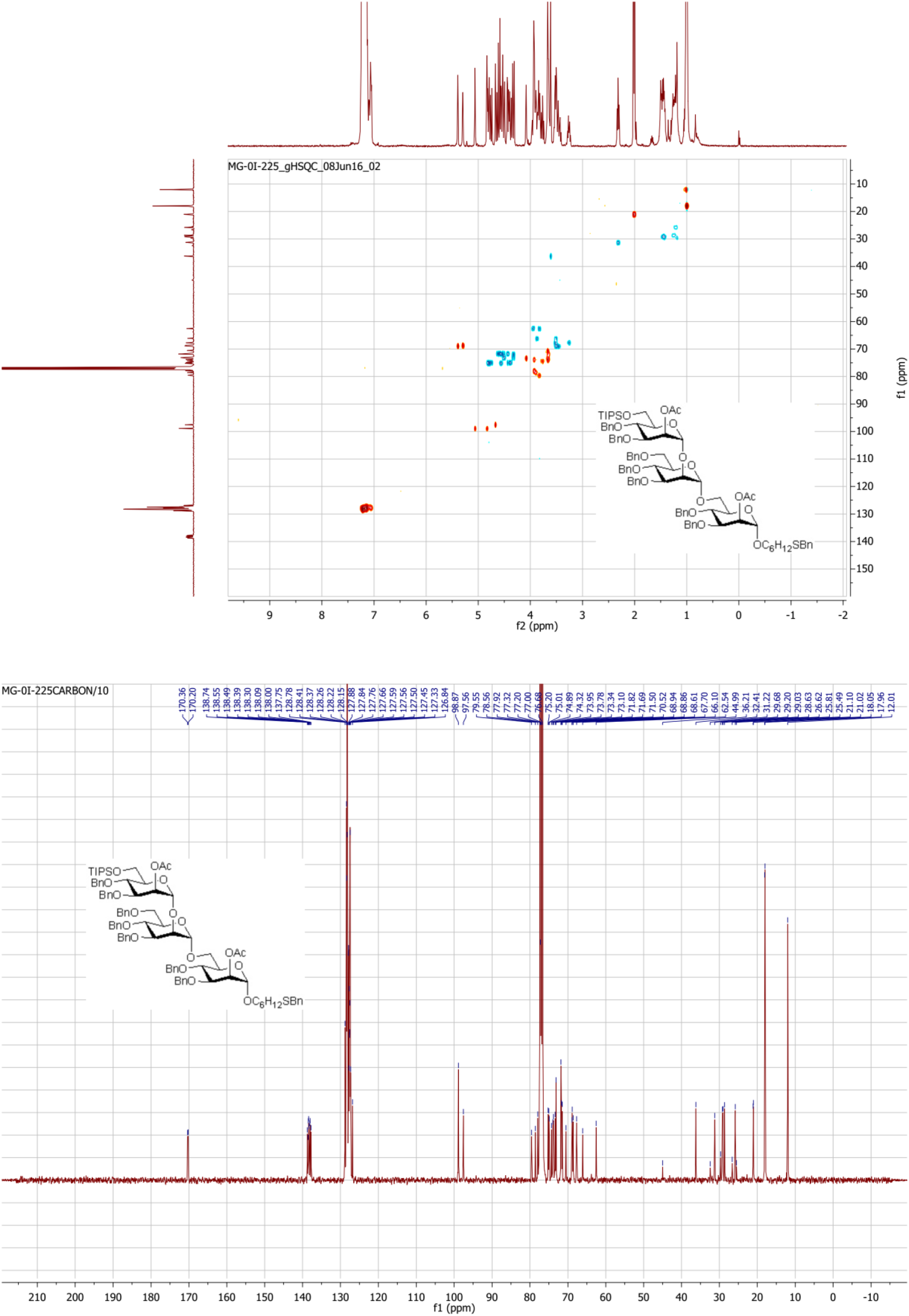
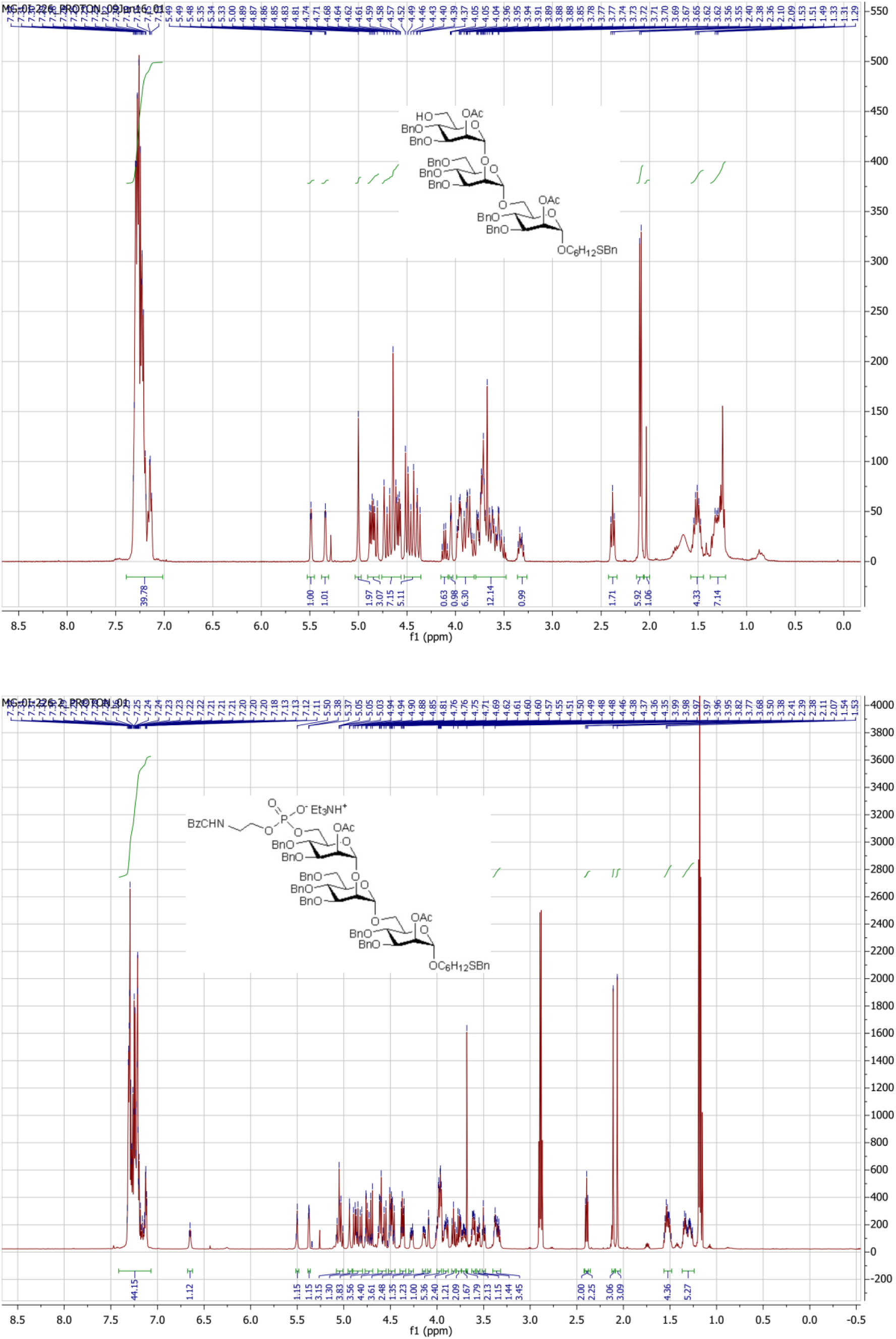
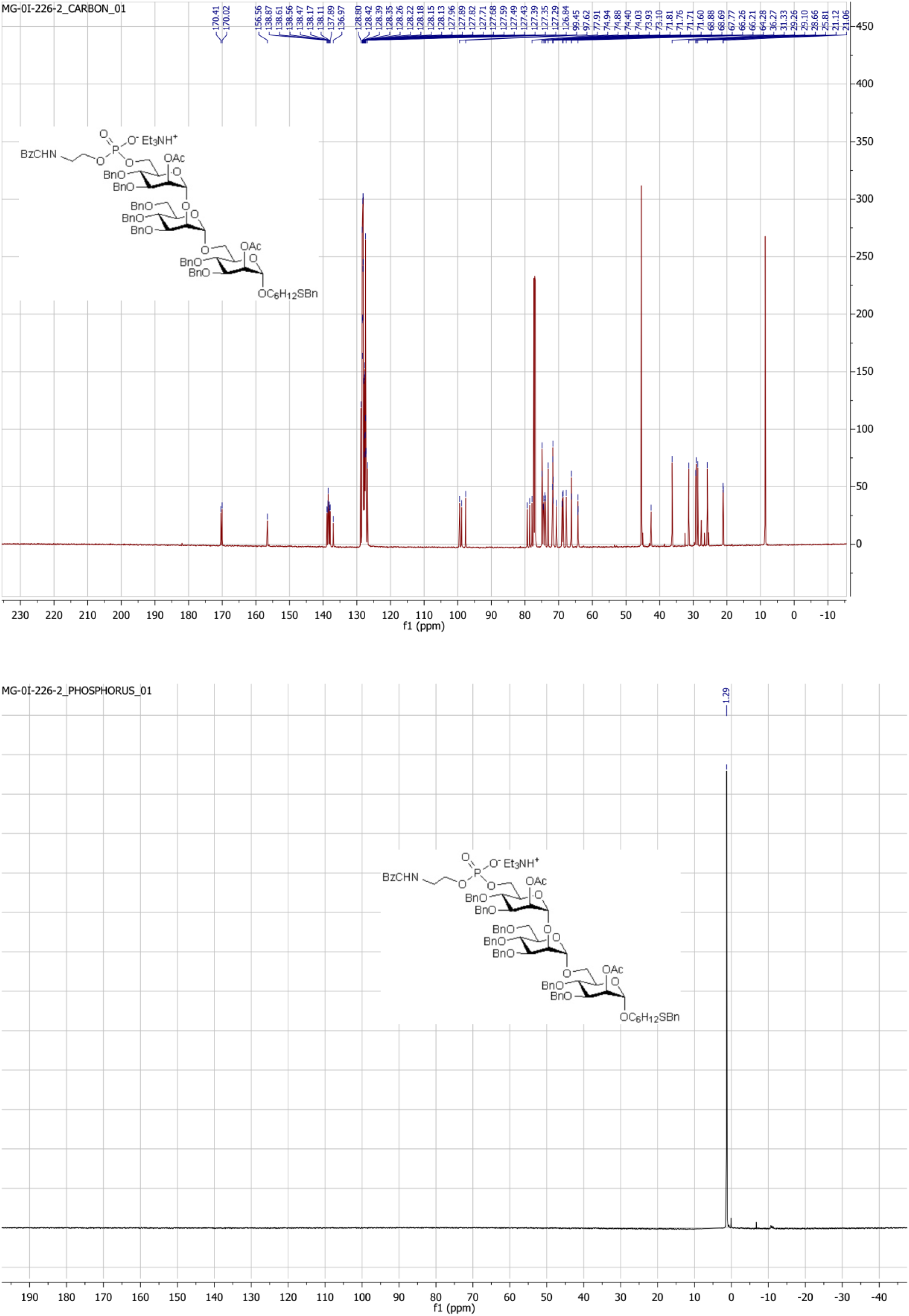
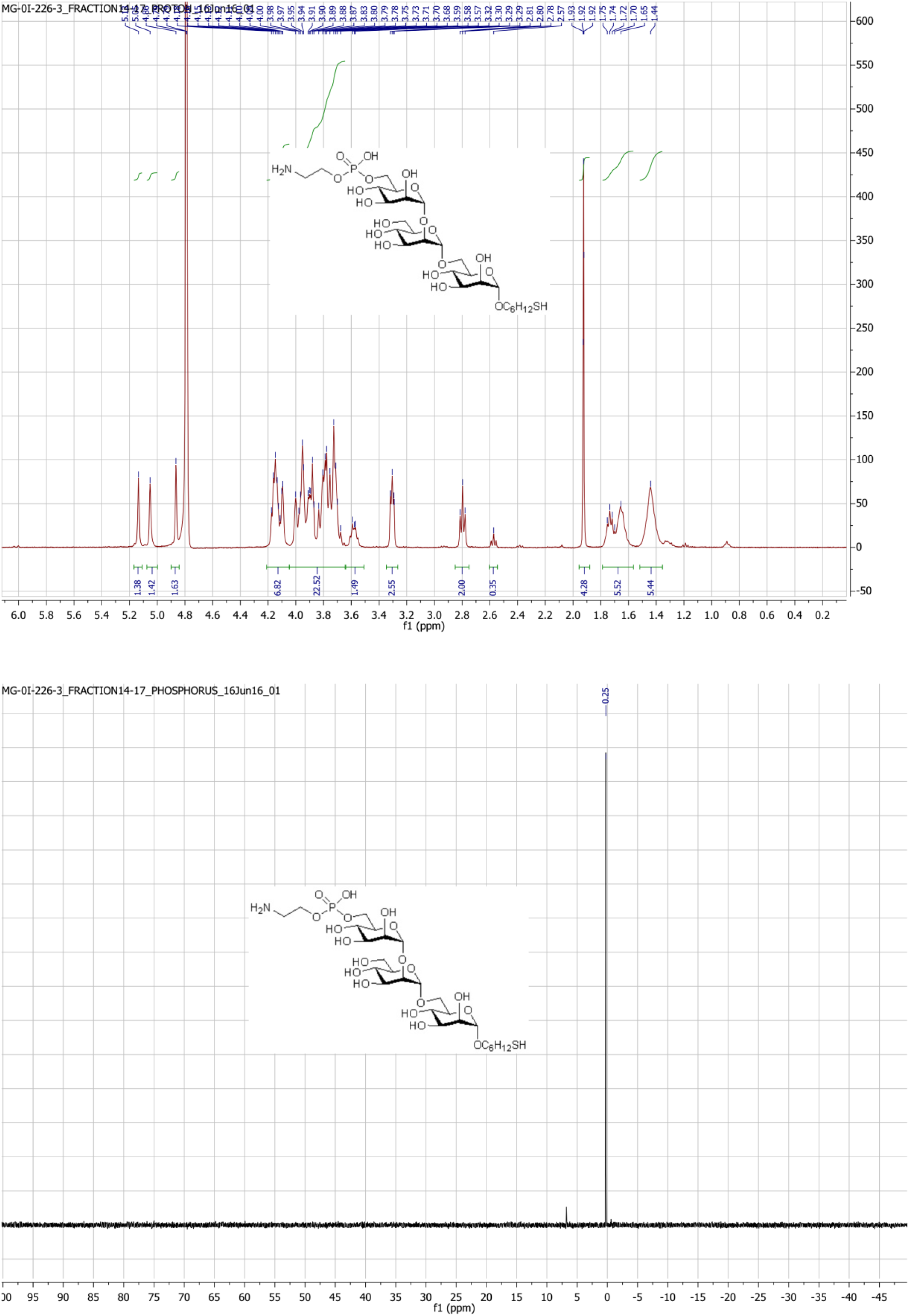
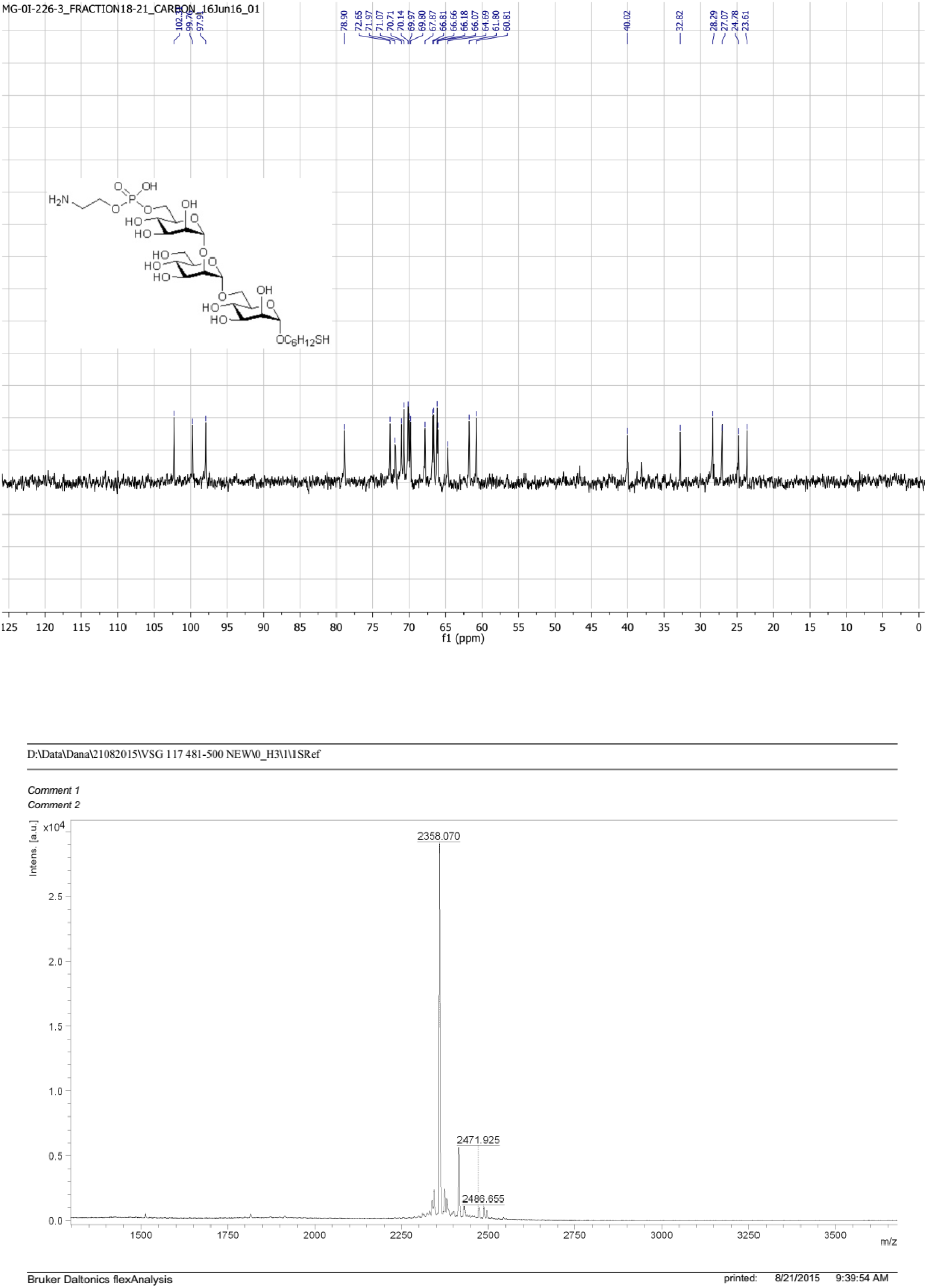

